# WATER reveals heterochrony of molecular programs underlies developmental failure caused by minor spliceosome inhibition

**DOI:** 10.64898/2026.05.08.723603

**Authors:** Saren M. Springer, Abigail R. Boria, Kyle D. Drake, Kevon O. Afriyie, Kaitlin N. Girardini, Taveena Konakanchi, Isabella Stevens, Nicolas Camacho, Tomas Lopes, Rahul N. Kanadia

**Affiliations:** Department of Physiology and Neurobiology, University of Connecticut, Storrs, CT 06269, USA; Institute for Systems Genomics, University of Connecticut, Storrs, CT 06269, USA; Stem Cell and Regenerative Biology Program and Division of Hematology/Oncology, Children’s Hospital Boston, Howard Hughes Medical Institute, Boston, MA 02115, USA; Harvard Medical School, Boston, MA 02115, USA; Department of Plant, Soil and Microbial Sciences, Michigan State University, Lansing, MI 48824, USA

## Abstract

The final limb structure reflects coordinated deployment of molecular programs, defined not only by which genes are expressed but when they are activated and silenced across time. Existing omics analyses obscure the temporal unfolding of these programs and conflate program identity with deployment timing by assuming temporal equivalence between conditions. We developed **WATER** (**W**eighted Windowed **A**ssignment of **T**emporal **E**xpression of **R**NA), a framework that reconstructs temporal gene expression trajectories independently within each condition, enabling direct comparison of temporal program architecture between wild-type and perturbed systems. Applying WATER to U11-null mouse forelimb development revealed that minor spliceosome inhibition redistributes genes across inappropriate temporal trajectories. Minor spliceosome inhibition causes splicing defects in minor intron-containing genes such as the PRC2 component *Eed*, leading to reduced H3K27me3 deposition and chromatin-transcription divergence. Single-cell RNA sequencing revealed persistence of progenitor states, impaired chondrogenic progression, and p53-dependent apoptotic checkpoint activation. Orthogonal WATER analysis of *Eed*-knockout stem cells recapitulated key features of chromatin gating failure, including temporal redistribution of skeletal development programs and progenitor state persistence, confirming that Eed loss alone is sufficient to produce temporal program redistribution independently of other splicing defects. Trp53 ablation in U11-null limbs partially rescued distal limb structures without correcting the underlying splicing defects, establishing that checkpoint activation amplifies rather than initiates the timing disruption. The limb retains much of its molecular toolkit but executes it in the wrong order, demonstrating that developmental failure arises from mistimed deployment of intact molecular programs. Thus, temporal program architecture is a fundamental organizing principle of morphogenesis.

## Introduction

The final form of a tissue does not reveal the path taken to generate it. Development depends not only on which genes are expressed, but on when they are activated and silenced as molecular programs unfold across time^1^. Classical developmental biology has focused on spatial patterning, defining how morphogen gradients and signaling centers specify where structures form^2–7^. Now, with the advent of omics data, we can extract the temporal organization of gene expression programs that govern the sequence of developmental transitions driving progression from progenitor maintenance to lineage commitment and tissue morphogenesis. As development proceeds and cell types diversify, coordinated timing between interacting cell populations becomes essential^8–10^. Thus, developmental defects could arise from loss of molecular programs, loss of temporal synchrony in their deployment, or simultaneous loss of both. These two modes of failure are mechanistically distinct but indistinguishable by existing approaches such as stage-matched static transcriptomic comparisons, which assume that equivalent timepoints correspond to equivalent developmental states^11^. This assumption fails when developmental timing itself is perturbed^12–14^. In such cases, a temporally shifted program appears as a distinct expression state rather than a heterochronic molecular program, making the distinction between program loss and program mistiming experimentally intractable. The concept of heterochrony, defined as changes in the timing of developmental processes, has long been recognized as a driver of morphological diversity, with compelling examples showing how altered timing of individual genes and signaling modules, including Hox codes, Shh duration, and retinoic acid gradients, reshapes limb morphology across species^15–19^. Yet heterochrony at the level of coordinated molecular programs, operating within a single genetic perturbation as a disease mechanism rather than an evolutionary adaptation, has remained inaccessible, precisely because the tools to measure program-level temporal architecture have not existed.

To make developmental timing measurable, we developed WATER (**W**eighted Windowed **A**ssignment of **T**emporal **E**xpression of **R**NA), a framework that reconstructs temporal gene expression trajectories without assuming stage equivalence between wild-type (WT) and mutant samples. By integrating expression across overlapping temporal windows, WATER defines whether genes are transiently activated, persistently expressed, or transitioning between states. It enables comparison of temporal program architecture between conditions, allowing direct comparison of when and in what order molecular programs are deployed. WATER therefore transforms temporal program architecture into a measurable regulatory dimension distinct from gene identity, expression level, or spatial distribution. In the developing limb, inhibition of the minor spliceosome provides a compelling test case. Conditional ablation of the U11 snRNA, a key component of the minor spliceosome, produces micromelia that phenocopies the human disorder microcephalic osteodysplastic primordial dwarfism type 1 (MOPD1), and morphological analysis revealed that mutant limbs follow a structurally distinct developmental path from their WT counterparts^20–25^. Stage-matched comparisons suggested a delay in developmental progression but could not resolve whether this reflected genuine temporal redistribution of molecular programs or other sources of transcriptional divergence. This distinction motivated the development of WATER, which provides a systematic quantitative framework for analyzing how temporal perturbations of gene regulatory networks produce developmental phenotypes, an approach the field has identified as essential^16^.

Applying WATER to bulk RNA-seq of WT and U11-null (mutant) mouse forelimbs across four developmental stages revealed that minor spliceosome inhibition redistributes genes across inappropriate temporal trajectories, expanding progenitor maintenance programs while delaying cell fate commitment programs—a form of molecular heterochrony invisible to static analysis. This redistribution is anchored by early mis-splicing of chromatin-regulatory minor intron-containing genes (MIGs), most critically the PRC2 component Eed, whose defective processing reduces H3K27me3 deposition and decouples chromatin state from transcriptional timing across the developing limb. Orthogonal WATER analysis of Eed-knockout stem cells confirmed that Eed loss alone is sufficient to recapitulate core features of temporal program redistribution independently of other splicing defects. Single-cell RNA sequencing connected transcriptome-level heterochrony to cellular heterochrony, revealing persistence of progenitor states and impaired chondrogenic progression. Loss of osteochondroprogenitors was blocked by ablation of Trp53 in U11-null limbs to partially rescue distal limb structures without correcting the underlying splicing defects, establishing that checkpoint activation amplifies rather than initiates developmental failure. Together, these findings demonstrate that temporal program architecture is a fundamental organizing principle of morphogenesis, and that heterochrony of molecular programs can further explain pathogenesis of developmental diseases.

## Results

We first reconstructed the temporal topology of WT forelimb development using WATER, then determined how U11 loss redistributes molecular programs across inappropriate temporal trajectories. We traced this redistribution to early mis-splicing of chromatin-regulating MIGs, particularly the PRC2 component Eed, and confirmed through orthogonal analysis of Eed-knockout stem cells that Eed loss is sufficient to produce the trajectory redistribution of genes. Single-cell RNA sequencing revealed how transcriptomic heterochrony manifests as cellular heterochrony, and genetic ablation of Trp53 established that the resulting p53-dependent checkpoint amplifies rather than initiates developmental failure.

### Minor spliceosome inhibition produces morphological and transcriptomic divergence

We sought to understand the molecular basis of developmental failure caused by minor spliceosome inhibition^20^. We collected WT and mutant littermate forelimbs at E10.5, E11.5, E12.5, and E13.5, revealing a distinct morphological deviation in mutant limbs across time (Fig. 1A–B). Quantification of limb surface area showed that mutant and WT limbs were comparable at E10.5 but diverged sharply after E11.5, resulting in severe size reduction by E13.5 (Fig. 1C, Table S1). Bulk RNA-seq across these timepoints followed by principal component analysis (PCA) separated early and late developmental stages along PC1 while genotype separated along PC2 (Fig. 1D, S1). Inherent in harvesting limbs at the same embryonic time is an expectation that the mutant limb is at a comparable stage in its developmental timing as its WT counterpart. Consequently, static RNA-seq between stage-matched limb analysis is a first natural step that revealed a general upregulation of genes in the mutant that was highest at E13.5 (Fig. 1E, Data S1). GO term enrichment of differentially expressed genes (DEGs) at each timepoint were distinct such as anterior-posterior patterning for genes that were upregulated in the E10.5 mutant, whereas muscle development and cartilage development were enriched by genes downregulated in the E10.5 mutant limb (Fig. S1A, Data S2). The mutant E11.5 upregulated genes enriched for GO terms such as stem cell population maintenance. In contrast, downregulated genes enriched for muscle contraction. The same analysis at E12.5 showed enrichment of induction of apoptosis by the upregulated genes and cartilage development by those that were downregulated. Likewise, genes upregulated in the E13.5 mutant limb enriched for cartilage development. Surprisingly, so did the genes downregulated in the mutant limb (Fig. S1A, Data S2). This suggested a distinct profile of gene expression patterns with specific molecular programs that are disrupted in the mutant. Intersection of differentially expressed genes across all timepoints revealed largely non-overlapping gene sets at each timepoint (Fig. 1F-G), indicating dynamic and stage-specific transcriptional perturbations rather than a stable mutant signature. Thus, despite the mutant limb being harvested along with the WT at the same time, they may in fact not be stage-matched in their developmental progression. Indeed, projection of E11.5 mutant samples onto WT principal component space revealed clustering with E10.5 WT samples (Fig. 1H, S1B-H), suggesting that mutant E11.5 limbs occupy an earlier transcriptional state than their stage-matched counterparts. This finding is consistent with the distinct morphological trajectory of the mutant limb compared to the stage-matched WT limb (Fig. 1B-C). Conceptually, we were inspired to shift our approach and explore novel ways to capture the dynamic shifts in gene expression across time that break free of chronological equivalence and treat the WT and mutant progression independently.

**Figure 1.**
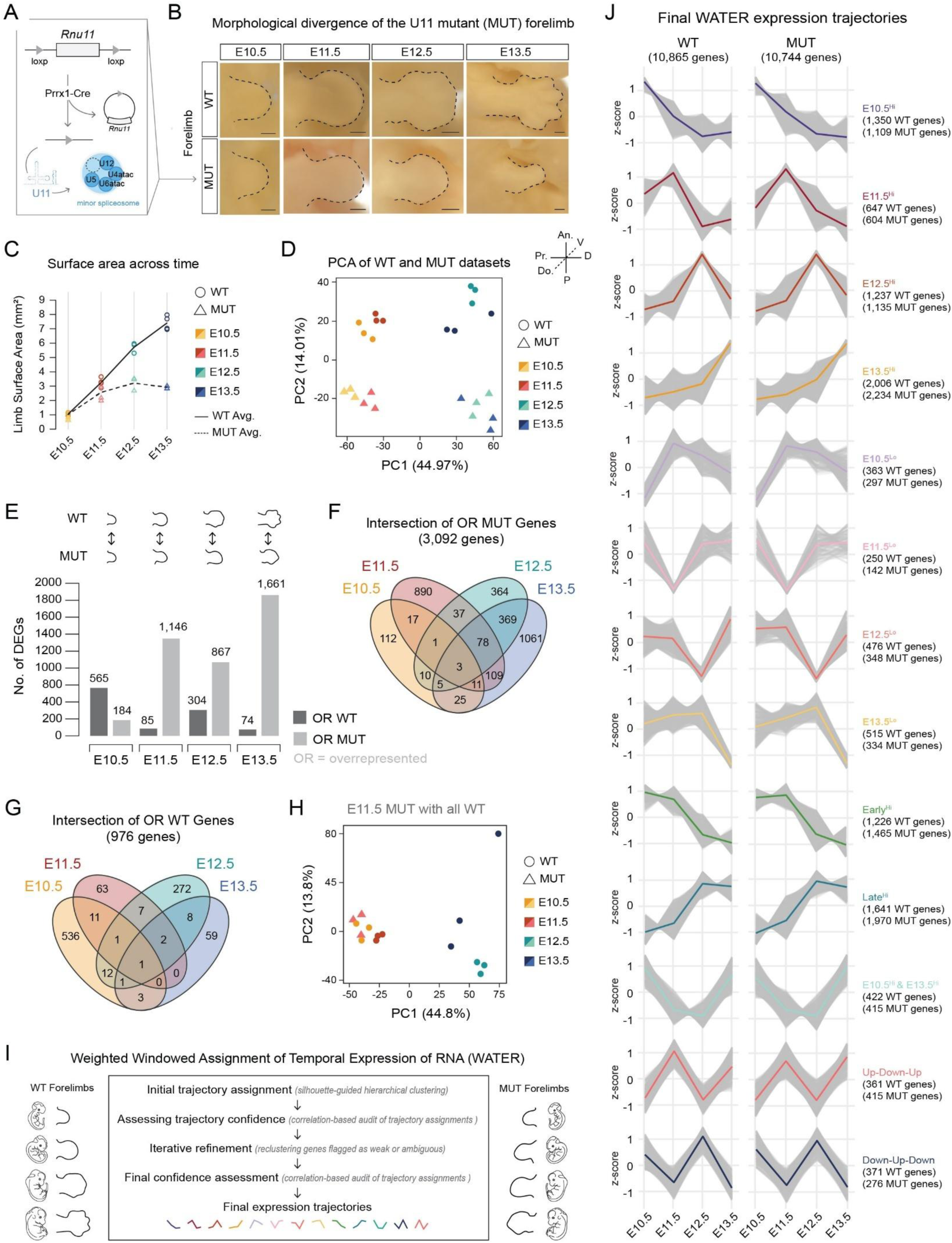
Stage-matched analyses suggest developmental delay upon U11 loss and motivate trajectory-based reconstruction of temporal gene expression programs using WATER. (A) Schematic of the conditional deletion strategy used to disrupt the minor spliceosome at the onset of limb development via *Prrx1*-Cre-mediated removal of Rnu11, resulting in loss of the U11 snRNA. Basic characterization of the U11-null mouse model and limb defects following *Prrx1*-Cre mediated recombination have been previously reported (Baumgartner et al. 2018; Drake et al. 2020). (B) Representative brightfield images of WT and mutant (MUT) forelimbs from E10.5 to E13.5, with limb outlines traced to illustrate gross morphological differences. (C) Quantification of divergence in limb surface area between WT and mutant forelimbs across time (mm^2^). Individual data points represent biological replicates; lines indicate mean surface area per genotype (WT, solid line; MUT, dashed line). Statistical significance was assessed using a one-way ANOVA followed by post hoc Tukey’s test (see also Table S1). (D) PCA of bulk RNA-seq datasets from WT and mutant forelimbs at E10.5, E11.5, E12.5, and E13.5. Samples cluster by genotype and stage. (E) Summary of pairwise differential expression (DE) analysis between WT and mutant forelimbs at matched developmental stages, referred to as static comparisons. Bar plots show the number of genes overrepresented (OR) in WT (dark grey) or mutant (light grey) at each timepoint. (F) Intersection of 3,092 OR mutant genes shows minimal overlap across static comparisons. (G) Intersection of OR WT genes (976 genes) likewise show minimal overlap across static comparisons. (H) PCA projection of E11.5 mutant samples combined with all WT timepoints show that E11.5 mutant clusters with E10.5 WT. (I) Weighted Windowed Assignment of Temporal Expression of RNA (WATER) pipeline. DE genes were initially assigned to clusters using silhouette-guided hierarchical clustering of z-scored expression profiles. Each gene’s expression profile was then compared to the mean profile of all candidate trajectories using Pearson correlation. Genes where the strongest correlation was to their originally assigned trajectory were retained, whereas genes with a stronger correlation to an alternative trajectory were reassigned. Genes with weak or ambiguous correlations were flagged for iterative refinement. Secondary silhouette analysis and hierarchical clustering were applied to weak and ambiguous genes independently. Following re-clustering, a secondary correlation-based audit was conducted to establish final high-confidence trajectory assignments. See Figures S1-S11 for a complete schematic of the WATER pipeline, detailed clustering, correlation, and reproducibility analyses of WT and mutant data. (J) Final WATER expression trajectories. Thirteen high-confidence temporal expression trajectories were defined for WT (left) and mutant (right) forelimbs following iterative refinement. Individual z-scored gene trajectories are shown in grey, with colored lines representing mean expression profiles for each trajectory. Gene counts per genotype are shown to the right of each trajectory plot. Following WATER, high-confidence trajectories were assigned to 97.5% (10,865 genes) of expressed WT genes and 97.8% (10,744 genes) of expressed mutant genes. An.; anterior, P; posterior, D; distal, Pr.; proximal, Do.; dorsal, V; ventral. See also Figures S1-S11, Table S1, Data S1-S4.

### WATER reconstructs temporal gene expression topology

To resolve this limitation, we developed **W**eighted Windowed **A**ssignment of **T**emporal **E**xpression of **R**NA (WATER), a trajectory-based analysis framework that reconstructs gene expression topology independently within WT and mutant limbs. Here topology refers to the architecture of changes in expression kinetics of genes across time, enabling identification of molecular programs that they execute (Fig. 1I). WATER infers molecular program topology by integrating overlapping temporal windows rather than relying solely on adjacent timepoint or static comparisons (Fig. 1I, S2, STAR Methods). Each timepoint is compared to progressively extended intervals, for example E10.5 versus E11.5, E10.5 versus E12.5 through E13.5, expanding the temporal window used to infer gene expression structure independently within each genotype (Fig. S3-S4, Data S3). This windowing strategy is key, because by integrating information across intervals of different lengths, WATER resolves whether a gene is transiently elevated at a single stage, persistently elevated across multiple stages, or transitioning between expression states, distinctions that adjacent-timepoint comparisons often conflate. Extended windows resolve sustained or multi-phase regulatory programs, while shorter intervals preserve sensitivity to transient shifts (Fig. S3). Critically, WATER reconstructs these topologies independently in WT and mutant, without assuming that the samples harvested at the same timepoint are developmentally equivalent, making it directly applicable to conditions where developmental timing itself is altered. This condition-independent reconstruction is the key conceptual shift, because it allows programs to be compared based on their intrinsic temporal structure rather than their position relative to a fixed chronological reference. This framework reconstructs coordinated molecular program deployment and redistribution, establishing temporal topology as a quantifiable regulatory dimension rather than a byproduct of stage-matched differential expression.

### Extraction and refinement of temporal gene expression trajectories

WATER-based hierarchical clustering of DEGs was performed separately for the WT and mutant samples. For each genotype, clusters were collapsed into coarse trajectory classes based on mean expression topology such as E10.5^Hi^, Early^Hi^, Late^Hi^, and corresponding low states (Fig. 1I, S2–S5, STAR Methods). Restricting the analysis to DEGs allowed us to focus on genes exhibiting dynamic temporal behavior, which would otherwise be overshadowed by NonDE genes. Moreover, NonDE genes would further confound GO term enrichment analysis used to identify the biological processes executed by genes with shared trajectories. Generally, the robustness of hierarchical clustering is not tested^26–29^, but given the value they hold for inferring molecular programs, we applied leave-one-out (LOO) stability analysis to discover the need for further refinement (Fig. S5). Therefore, we undertook correlation-based reassignment to optimize gene assignment (Fig. 1I, S6A, STAR Methods). In this approach, each gene expression profile is compared to the mean profiles of all candidate trajectories using Pearson correlation, and genes with a stronger correlation to an alternative trajectory are reassigned accordingly. Iterative reassignment substantially improved assignment stability and resolved ambiguous profiles (Fig. 1I, S6-S8). Ultimately, ∼97.5% of expressed genes were confidently assigned to 13 discrete temporal trajectories in both WT (97.5%; 10,865/11,141 DEGs, Fig. S8B) and mutant (97.8%; 10,744/10,975 DEGs, Fig. S8E) datasets (Fig. 1J, S8, Data S4), establishing a robust temporal architecture for comparative analysis. Final LOO analysis showed retention of ∼75–85% for the WT and ∼90% for the mutant (Fig. S9A, B, G, & H), a marked improvement from an initial LOO of ∼45% for both WT and mutant data (Fig. S5B-C).

### Validation and benchmarking of WATER trajectory inference

Biological fidelity of WATER-derived trajectories was validated against prior temporal forelimb studies and developmental regulators with established expression profiles (Fig. S9C-F). Of the 4,329 genes analyzed by Jhanwar et al., there was sufficient data to compare 2,445 genes^28^. The expression trajectories for 1,880 genes were concordant with our analysis (Fig. S9C). For the 565 discordant genes, we cross-referenced expression in the dataset published by Onimaru et al., which showed agreement for 286 genes (Fig. S9C)^29^. Canonical early patterning genes (*Shh, Fgf8, Alx4*), cell fate (*Msx1, Hand2, Hoxa11, Sall1*), and chondrogenesis (*Col2a1, Sox6, Acan*) genes mapped to expected temporal trajectories, confirming that WATER captures established developmental logic while improving resolution of complex temporal dynamics (Fig. S9D-F). To further benchmark performance, we compared WATER to TC-Seq, an unsupervised temporal clustering method (Fig. S10A, Table S2)^30^. Applying TC-Seq to WT data yielded 10 clusters corresponding to WATER trajectories, with some redundancy, and identified three WATER trajectory classes that lacked clear TC-Seq counterparts (Fig. S10B–C). Overall, 71.3% of genes were concordantly assigned between methods, indicating preservation of the global developmental architecture, but 28.7% exhibited discordant assignments (Fig. S10D-E, G). For example, *Sall3* and *Col2a1* showed overlap in expression trajectories by TC-Seq and WATER (Fig. S10F), whereas *Hoxd9* and *Actb* displayed multi-phase expression patterns that were grouped into single TC-Seq clusters but separated into distinct temporal states by WATER (Fig. S10H). This demonstrates that iterative correlation-based refinement resolves biologically meaningful expression dynamics that can be conflated by a single pass unsupervised clustering. Functional enrichment confirmed that WATER-defined trajectories correspond to coordinated biological modules rather than arbitrary expression clusters (Fig. S11, Data S5). In WT limbs, distinct temporal states were enriched for processes consistent with limb developmental progression, including early patterning and signaling modules, proliferative programs, chromatin regulation, and skeletal differentiation. These GO term enrichments mapped selectively to specific trajectories, indicating that trajectory identity reflects structured temporal developmental architecture (Fig. S11A). When applied to the mutant dataset, GO terms associated with embryonic skeletal system development and morphogenesis, which were enriched in early WT trajectories, appeared redistributed across delayed or altered temporal states, providing an initial transcriptomic indication that minor spliceosome (MiS) inhibition via U11 ablation disrupts the timing of coordinated developmental programs (Fig. S11A’–J, Data S6).

### Minor spliceosome inhibition displaces molecular programs from their normal temporal topology

Although the same 13 trajectory classes were preserved across genotypes, their relative representation was significantly rebalanced following U11 loss (Fig. 1J). For example, 43 genes that enriched for *“DNA replication*” were found in the WT-E10.5^Hi^ trajectory, but these genes were found in the Mut-E12.5^Hi^ trajectory (Fig. S11A). Similarly, 17 genes in the WT-E10.5^Hi^ trajectory underpinning “*embryonic skeletal system development*” were redistributed in the mutant to E10.5^Hi^ (6), NonDE (5), Early^Hi^ (4), E12.5^Lo^ (1) and E11.5^Hi^ (1) (Fig. S11B). In contrast, 11 of the 55 genes in the WT-E10.5^Hi^ trajectory that enriched for “mitotic cell cycle phase transition” were preserved in the Mut-E10.5^Hi^ trajectory (Fig. S11A). Thus, redistribution was selective rather than global, as a subset of trajectories displayed comparatively modest differences between WT and mutant limbs. We identified distinct shifts in gene expression topology in the mutant limb for all 13 WT trajectories that enriched for diverse molecular programs that can be mined further for pathway specific investigations (Fig. S12-15). In all, we observed preservation of gene expression trajectories between WT and mutant data for 36.9% of genes, whereas 63.1% of genes exhibited expression shifts between the two genotypes (Fig. S15C). Having established these global shifts in temporal state representation, we next examined gene-level reassignment between trajectories. Intersection analysis revealed widespread redistribution of genes between WT and mutant temporal states (Fig. 2A–A’, S12-15). Moreover, integrating temporal with static analysis showed that genes with trajectory shifts also tended to have significant magnitude changes by static comparison (Fig. S16). Thus, integrating both static and temporal analysis reveals a more complete picture of the molecular landscape of the developing limbs response to U11 loss.

**Figure 2.**
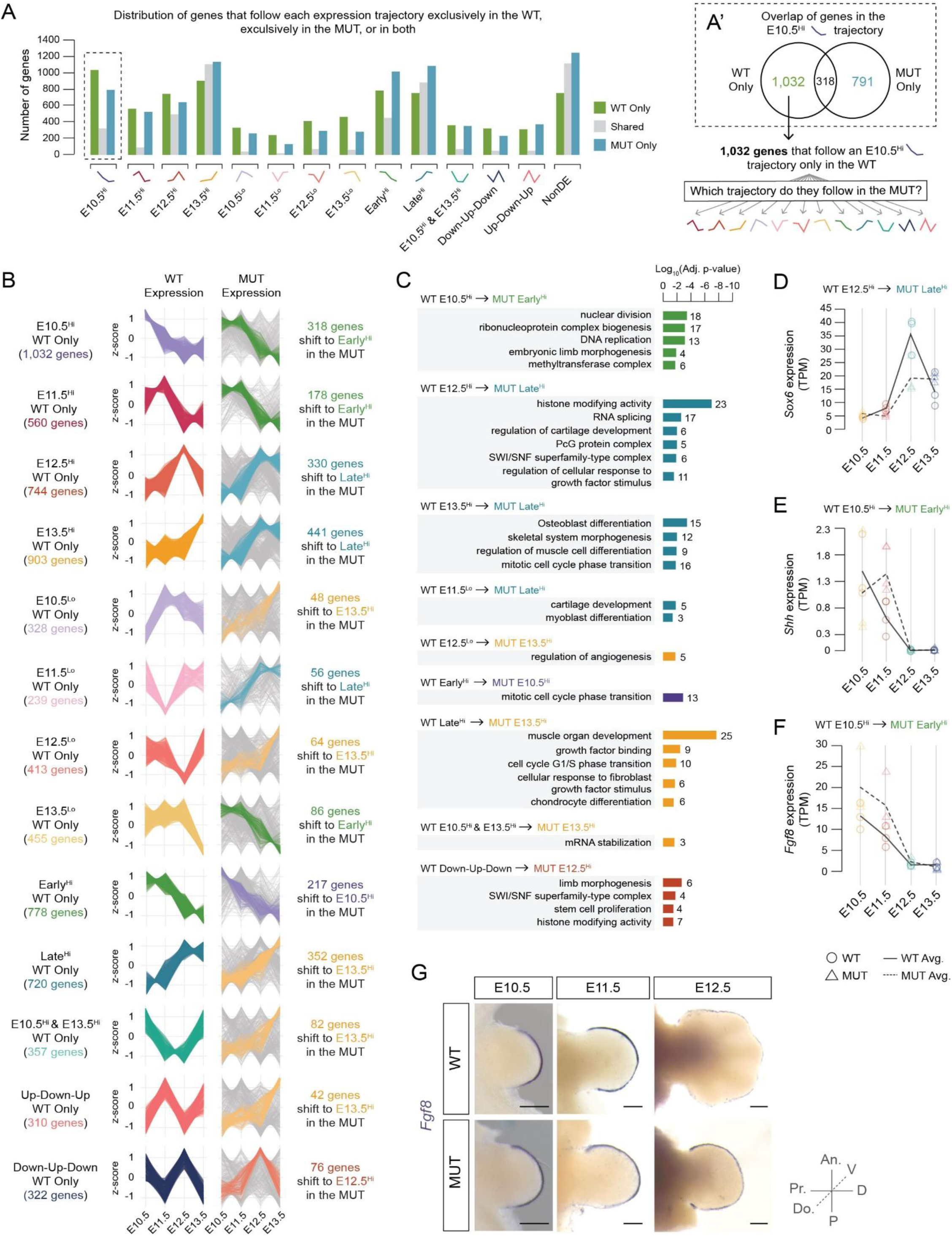
U11 loss redistributes genes across temporal expression trajectories during forelimb development. (A) Intersection of genes assigned to each expression trajectory in WT and mutant forelimbs. For each trajectory, genes were classified as WT only (green), mutant only (blue), or shared between genotypes (grey). (A’) Venn diagram highlighting overlap within the E10.5 high trajectory. While a subset of genes retain the E10.5 high trajectory in both genotypes (318 genes), a large fraction of WT-specific E10.5 high genes (1,032 genes) were reassigned to alternative trajectories in the mutant. (B) For each WT-only trajectory, gene expression trajectories (z-scored) are shown for WT (left; colored by trajectory) alongside the trajectories adopted by those same genes in the mutant (right). The most frequently reassigned mutant trajectory is highlighted in color, with the number of genes undergoing that shift indicated to the right; all other trajectories are shown in grey. (C) GO enrichment analysis for subsets of WT-only genes that shift to alternative trajectories in the mutant. Bars represent significantly enriched GO terms (BP, CC, MF; adjusted p < 0.05), with bar length indicating the log_10_(adjusted p-value) and numbers denoting the number of genes underlying enrichment of each term. (D-F) Representative examples of genes shifting from WT E12.5 high to mutant Late high (D, *Sox6*) and WT E10.5 high to mutant Early high (E, *Shh;* F, *Fgf8*). Lines show the mean expression (TPM) across time for WT (solid line, circles) and mutant (dashed line, triangles), with biological replicates overlaid. (G) Whole mount in situ hybridization (WISH) for *Fgf8* in WT (top) and mutant (bottom) forelimbs from E10.5 to E12.5, illustrating altered temporal expression dynamics of *Fgf8* upon U11 loss. See also Figure S17 for WT and mutant *Fgf8* WISH at E13.5, confirming the absence of *Fgf8* expression in either genotype. An.; anterior, P; posterior, D; distal, Pr.; proximal, Do.; dorsal, V; ventral. Scale bars, 0.5 mm. See also Figures S12–S17, Table S3, Data S7.

WT-specific trajectories, including stage-restricted early and mid-developmental programs, frequently resolved into multiple mutant trajectories, indicating loss of coordinated temporal deployment (Fig. 2A–B). For example, 318 of 1,032 genes in the WT-E10.5^Hi^ trajectory were reassigned to the Mut-Early^Hi^ trajectory (Fig. 2A’–B). GO term analysis demonstrated that these shifts occurred at the level of functional modules, including programs associated with nuclear division, morphogenesis, and epigenetic modification (Fig. 2C, Table S3, Data S7). Strikingly, this redistribution extended across all WT trajectory classes. Early, mid, late, low-state, and biphasic programs each lost unified temporal coherence and were redistributed into structured mutant states, yet reassigned subsets retained their functional enrichment signatures (Fig. S12–S15, Table S3, Data S7). Early programs redistributed into contexts enriched for cytoskeletal remodeling and genome maintenance, whereas late differentiation programs were redeployed into mutant states retaining skeletal and morphogenetic signatures (Fig. 2C, S12–S15). Thus, minor spliceosome inhibition disrupts coordinated program execution while preserving functional identity, producing temporal desynchronization rather than wholesale transcriptional failure.

### Experimental validation confirms temporal shift

*Sox6*, normally assigned to a defined differentiation trajectory in WT limbs, exhibited reduced and redistributed expression in the mutant (Fig. 2D). *Shh* and *Fgf8* similarly showed altered temporal deployment consistent with trajectory reassignment (Fig. 2E–F). Whole mount in situ hybridization confirmed persistence of *Fgf8* in the mutant E12.5 forelimb, when it is mostly lost in the WT E12.5 forelimb (Fig. 2G). Notably, trajectory reassignment was often accompanied by altered expression magnitude across developmental time (Fig. S16), suggesting that U11 loss reshapes both the timing and the amplitude of gene deployment, and demonstrating that the regulatory logic governing these genes is disrupted at multiple levels simultaneously. Quantification of RT-PCR validation of *Msx1* (WT-Late^Hi^; MUT-E12.5^Hi^), *Msx2* (WT-E13.5^Hi^; MUT-E12.5^Hi^), and *Col2a1* (WT-E13.5^Hi^; MUT-Late^Hi^) confirmed expression shifts consistent with WATER results (Fig. S17). Together, these validations confirm that U11 loss perturbs program deployment during limb morphogenesis. Developmental programs are preserved, yet their temporal coordination is uncoupled from normal progression, demonstrating that limb malformation arises from heterochrony of multiple molecular programs rather than isolated changes in gene expression at an isolated timepoint.

### MIG expression trajectories are disproportionately disrupted

Having established that MiS inhibition redistributes developmental programs globally, we next examined how MIGs, the direct targets of the MiS, are positioned within this restructured temporal framework^31–34^. In WT limbs, MIGs were distributed across all 13 temporal trajectories, ranging from 80 MIGs in the E12.5^Hi^ class to 65 MIGs in the NonDE class and 3 in the Up-Down-Up class (Fig. 3A), reflecting broad participation of MIGs in diverse developmental programs. Following MiS inhibition, although MIGs broadly mirrored the global pattern of trajectory redistribution, they exhibited a pronounced bias toward mutant trajectories characterized by delayed, flattened, or sustained expression (Fig. 3A-B, Data S8). Projection of WT-defined MIG trajectories onto mutant states revealed an overall reduction of MIGs within their corresponding trajectories and accumulation in the Late^Hi^ and NonDE categories (Fig. 3A–B). These data suggest that MiS disruption alters the timing and stability of MIG-associated regulatory programs rather than causing a uniform decrease in expression. Consistent with this interpretation, MIGs that shifted trajectory were enriched for histone modifying activity, Erk1/2 signaling, and transcriptional regulation (Fig. 3C), pointing toward disruption of upstream regulatory hubs as one of the primary consequences of MiS inhibition.

**Figure 3.**
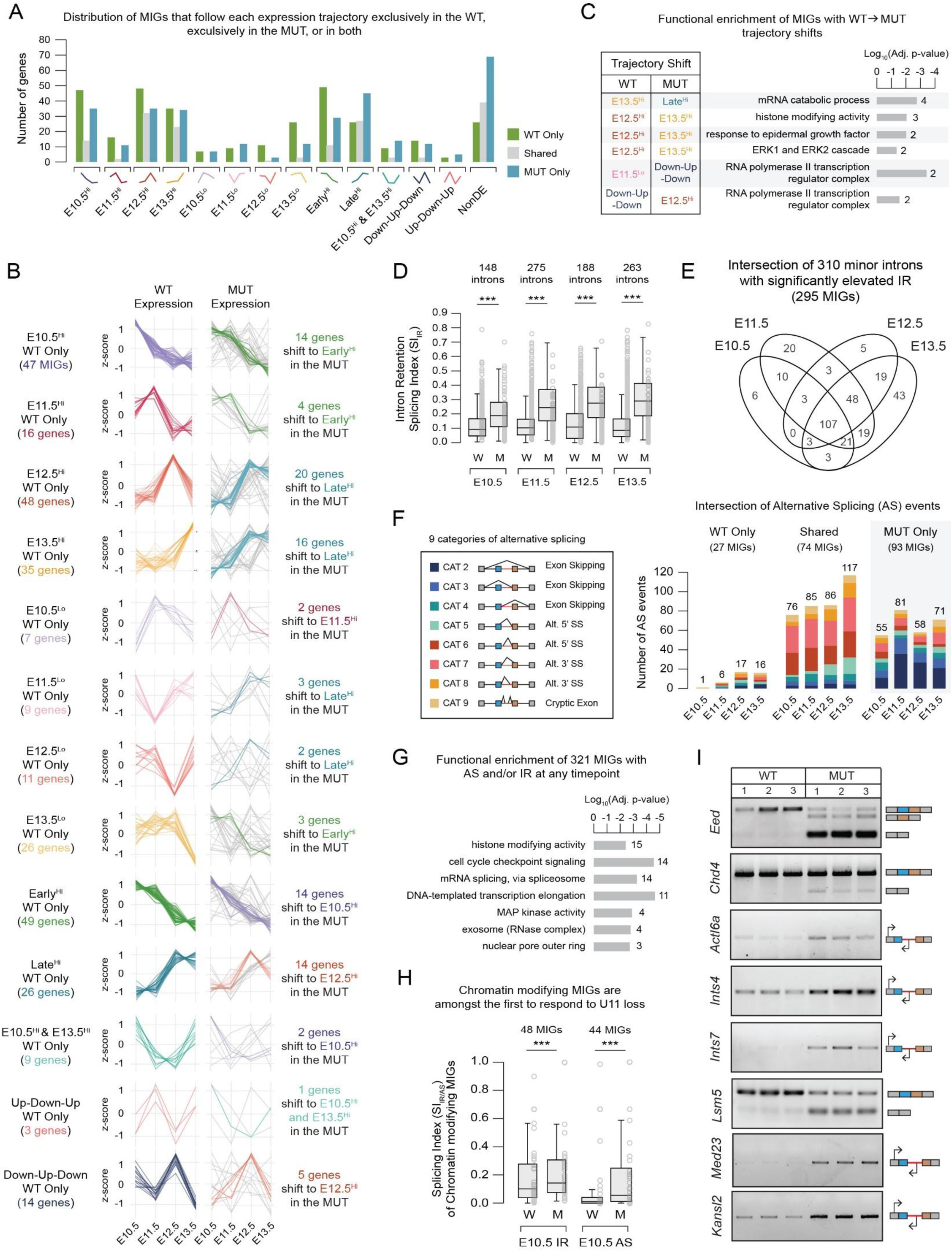
Altered expression and splicing dynamics of minor intron-containing genes across time in the U11 null forelimb. (A) Distribution of MIGs assigned to each expression trajectory in WT and MUT forelimbs. For each trajectory, MIGs are classified as WT only (green), mutant only (blue), or shared between genotypes (grey). (B) Z-scored expression trajectories for MIGs assigned exclusively to each trajectory in the WT. WT expression profiles are shown on the left (colored), and expression of the same genes in the mutant is shown on the right. MIGs reassigned to the most frequently observed trajectory shift are colored and their count is listed, while all other shifts are shown in grey. (C) Gene Ontology (GO) analysis of MIGs that shift their expression trajectory between the WT and mutant. Bars represent significantly enriched GO terms (BP, CC, MF; adjusted p-value < 0.05), grouped by WT-to-MUT trajectory shift. Bar length indicates the log_10_(adjusted p-value) and numbers denote the number of MIGs underlying each term. (D) Distribution of intron retention (IR) in WT (W) and mutant (M) limbs across developmental time, quantified as a splicing index (SI). Boxplots summarize SI values for all minor introns in expressed genes at each timepoint. Boxes represent the interquartile range (25^th^ to 75^th^ percentile), center lines indicate the median, and whiskers denote the 1.5x interquartile range. The number of introns analyzed is listed above each comparison. Wilcoxon rank-sum tests were used for statistical comparisons. (E) Intersection of minor introns that exhibit significantly elevated IR across time. The Venn diagram shows overlap of introns at E10.5, E11.5, E12.5, and E13.5, highlighting a core set of introns affected early and persistently following U11 loss. (F) Stacked bar charts depicting the frequency of alternative splicing (AS) events across nine categories for MIGs undergoing AS only in the WT (WT only), only in the mutant (MUT only), or in both genotypes (shared). AS events are colored by category. (G) Functional enrichment analysis of MIGs with elevated IR and/or AS at any timepoint. Bar length indicates the log_10_(adjusted p-value) and numbers denote the number of MIGs underlying each term. (H) Comparison of WT (W) and mutant (M) SI for IR and AS at E10.5 for chromatin modifying MIGs. Boxplots show that chromatin modifying MIGs exhibit early IR (48 introns) and AS (44 introns) following U11 loss. The number of introns analyzed is listed above each comparison. Wilcoxon rank-sum tests were used for statistical comparisons. *** p < 0.001. (I) RT-PCR validation of representative MIGs exhibiting intron retention or alternative splicing in E11.5 WT and mutant limbs. Schematics to the right indicate the type of alternative splicing or primer design used to assess intron retention. See also Figures S18–S21, Table S5, Data S8-S9.

### Early MIG mis-splicing preferentially disrupts chromatin and transcriptional regulators

We quantified minor intron retention (IR) and alternative splicing (AS) in MIGs across developmental time (Fig. 3D-F). We first defined the baseline splicing landscape of minor introns during WT limb development. Both IR and AS increase with developmental time in WT limbs, and MIGs exhibiting WT IR and/or AS are enriched for cell-cycle-associated functions, consistent with previous reports implicating MIG splicing in the regulation of cell proliferation (Fig. S18)^20,35–37^. This baseline established that some minor intron processing variability is a normal feature of WT development, making it essential to quantify mutant-specific changes relative to this background. In contrast to the gradual increase seen in WT, elevated IR in the mutant was observed as early as E10.5, coinciding with U11 ablation and preceding overt cellular or morphological abnormalities (Fig. 3D, 1B). These defects accumulated across time, producing stage-specific sets of mis-spliced MIGs and a core set of 107 MIGs consistently mis-spliced at all timepoints (Fig. 3D–E), indicating that MiS inhibition imposes a sustained burden on RNA processing rather than a transient early perturbation. Consistently, 93 MIGs exhibited mutant-specific alternative splicing (Fig. 3F). Functional enrichment of mis-spliced MIGs revealed overrepresentation of histone modifiers (15 MIGs), cell cycle regulators (14 MIGs), RNA splicing factors (14 MIGs), and transcriptional regulators (11 MIGs) (Fig. 3G). Chromatin-regulating MIGs were among the earliest affected, exhibiting elevated IR and AS at E10.5 (Fig. 3H, Data S9). Several MIGs central to chromatin and transcriptional control exhibited reproducible splicing defects, including *Eed* (PRC2), *Chd4* (NuRD), *Actl6a* (BAF), *Ints4/Ints7* (Integrator), *Med23* (Mediator), *Lsm5*, and *Kansl2* (Fig. 3I)^38–42^. IR did not uniformly correlate with reduced gene expression (Fig. S19), indicating that minor spliceosome inhibition reshapes transcript structure and turnover in a gene-specific manner rather than simply extinguishing MIG expression, consistent with previous findings^35^. This observation is important because it implies that functional disruption of MIG-encoded proteins can occur even when bulk transcript levels appear preserved, decoupling the splicing defect from any simple expression-level readout. Because IR and AS have the potential to disrupt protein production, alter functional domains, or impair complex assembly, we evaluated the predicted structural consequences of representative mis-splicing events (Fig. S20A-C’). Gene structure analysis of selected MIGs, including *Xrcc5*, *Exo1*, and *Kansl2*, revealed that their minor introns reside adjacent to exons encoding key protein domains (Fig. S20D-F). AlphaFold modeling of protein isoforms generated from validated AS events predicted truncation or distortion of domains required for protein-protein interactions and complex assembly (Fig. S20D–I), indicating that mis-splicing is likely to impair regulatory protein architecture even when overall transcript abundance is preserved. We next constructed a potential framework of disrupted MIGs and their interconnected hubs governing chromatin organization, transcriptional regulation, RNA processing, and DNA repair (Fig. S21), positioning MIGs as central coordinators of developmental program execution. Rather than broadly suppressing developmental genes, U11 loss thus perturbs a network of chromatin and genome-stability regulators whose functional disruption provides a mechanistic substrate for the temporal desynchronization revealed by WATER.

### Mis-splicing of *Eed* disrupts PRC2-dependent chromatin gating

Given that chromatin-regulating MIGs were among the earliest mis-spliced, we next investigated whether mis-splicing of epigenetic regulators translates into altered epigenomic dynamics. Among the early mis-spliced MIGs we focused on *Eed*, a core PRC2 component required for transcriptional repression during limb development (Fig. S21)^38,43^. The rationale for focusing on *Eed* was that *Prrx1-*Cre mediated ablation of *Eed* in the mouse also results in forelimb micromelia at birth^44^. Moreover, the unexpected upregulation of genes in the U11-null samples suggested either ectopic gene expression or failure to suppress gene expression (Fig. 1E). The latter possibility is consistent with the regulatory role of Eed in PRC2 activity, which is generally linked to suppression of gene expression during limb development by placing H3K27me3 marks^38^. Thus, mis-splicing of *Eed* and loss-of-function in the U11-null limb would compromise PRC2 function, impairing H3K27me3 deposition and causing failure to suppress gene expression across limb development. CUT&RUN profiling of H3K27me3 in WT and mutant forelimbs across E10.5–E12.5 revealed differential peak enrichment between genotypes at each timepoint (Fig. 4A–B, S22-23). The number of reads for the top 10,000 sites flanking the transcription start site was visibly reduced in the mutant limb at E12.5 (Fig. 4C, S24). The number of parent genes associated with altered H3K27me3 deposition increased progressively across developmental time, and representative loci including *Msx1* and *Myc* illustrate genotype-specific changes in both H3K27me3 signal and expression (Fig. 4D–F, S22, Data S10). Applying WATER to H3K27me3 profiles identified coherent chromatin trajectories, and integration with transcriptional trajectories revealed overall conservation of trajectory relationships alongside gene-level divergence (Fig. 4G–H, Data S11). A conserved class of loci, including *Ptch1*, *Bmp2*, *Wnt3*, *Lef1*, *Prrx1*, and *Runx2*, functionally enriched for limb morphogenesis and skeletal development and retained concordant H3K27me3 and expression relationships, indicating that core limb identity programs remain partially buffered despite MiS inhibition (Fig. 4I, Data S12). In contrast, three classes of regulatory divergence emerged. An expression-decoupled class showed altered transcription without concordant H3K27me3 changes, including patterning regulators *Bmp4*, *Dlx2*, *Sox9*, *Hoxb3*, *Tbx2*, *Tbx3*, and *Shox2* (Fig. 4J, Data S12). An H3K27me3-decoupled class showed altered chromatin without proportional transcriptional change, including differentiation regulators *Mef2d*, *Hdac7*, *Tbx5*, and *Sox8* (Fig. 4K, Data S12). A third rewired class exhibited inversion or breakdown of the normal chromatin-to-expression relationship, including core morphogenetic regulators *Msx1*, *Msx2*, *Myf5*, *Wnt7a*, *Pitx1*, and *Gdf5* (Fig. 4H–I, 4L, Data S12). GO term enrichment for these divergent classes highlighted pattern specification, cartilage development, BMP signaling, and mesenchyme development (Fig. 4I), indicating selective destabilization of highly leveraged developmental nodes rather than global PRC2 collapse. Representative loci further illustrate these distinct modes of chromatin-transcription uncoupling where some genes maintained elevated expression despite increased H3K27me3 deposition, whereas others failed to acquire repression despite declining transcription (Fig. S22). Together, these findings suggest that *Eed* mis-splicing disrupts PRC2-dependent chromatin gating, resulting in progressive epigenetic divergence and partial decoupling of chromatin state from transcriptional deployment. Transcriptomic topology in limb development is reinforced by PRC2-dependent chromatin gating, and minor spliceosome inhibition destabilizes this layer of regulation, providing a direct mechanistic bridge between early splicing defects (Fig. 3) and the downstream transcriptional heterochrony (Fig. 2).

**Figure 4.**
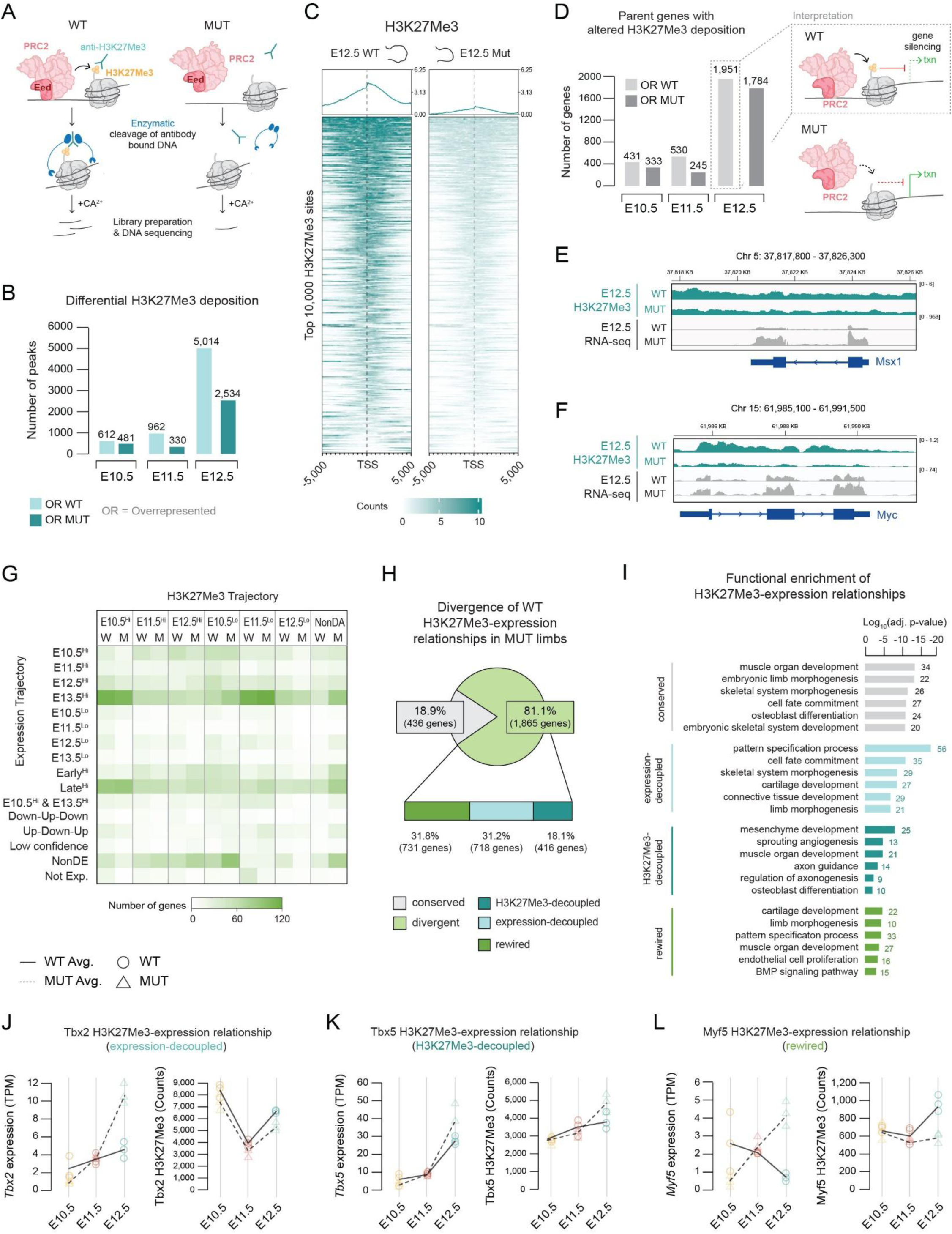
Altered coordination between H3K27Me3 deposition and gene expression during development of the U11 mutant forelimb. (A) Schematic of H3K27Me3 profiling in WT and mutant forelimbs using CUT&RUN. H3K27Me3-bound chromatin was isolated via antibody binding followed by enzymatic cleavage, enabling genome-wide assessment of H3K27Me3 in WT and mutant limbs. (B) Quantification of differentially enriched H3K27Me3 peaks between WT and mutant limbs at E10.5-E12.5. Bars indicate peaks overrepresented (OR) in each genotype relative to the other genotype at the same timepoint. (C) Heatmaps showing H3K27Me3 signal intensity centered on transcription start sites (TSS ±5 kb) in E12.5 WT and mutant limbs. The same top 10,000 WT-enriched sites are shown in both heatmaps. Average H3K27Me3 intensity profiles are shown above each heatmap. (D) The number of parent genes associated with altered H3K27Me3 deposition across time. The schematic inset illustrates the interpretation of observed H3K27Me3 deposition patterns in WT and mutant limbs and their predicted relationship to gene expression. (E-F) IGV tracks illustrating representative loci with altered H3K27Me3 deposition and expression in E12.5 mutant limbs compared to WT, including *Msx1* (E) and *Myc* (F). (G) Heatmap summarizing the relationships between H3K27Me3 and expression trajectories. Non-differentially abundant (NonDA) refers to genes with unchanging H3K27Me3 across time. (H) Classification of WT and mutant H3K27Me3-expression relationships. Pie chart indicates the proportion of genes whose H3K27Me3-expression relationship is conserved or altered in mutant relative to WT. Conserved (grey) indicates the same H3K27Me3-expression relationship was observed in both genotypes, whereas divergent (light green) indicates the mutant relationship differs from the WT baseline. Divergent genes were further subdivided into expression-decoupled (light blue), H3K27Me3-decoupled (dark blue), and rewired (dark green) categories. Expression-decoupled indicates the same H3K27Me3 trajectory was observed across genotypes while the expression trajectory differed. H3K27Me3-decoupled indicates the same expression trajectory was observed across genotypes while the H3K27Me3 trajectory differed. Rewired indicates both the H3K27Me3 and expression trajectory differed between genotypes. (I) GO enrichment analysis of conserved, expression-decoupled, H3K27Me3-decoupled, and rewired genes. (J-L) Representative examples of genes exhibiting expression-decoupled (*Tbx2*, J), H3K27Me3-decoupled (*Tbx5*, K), or rewired (*Myf5*, L) H3K27Me3-expression relationships. Left panels show gene expression (TPM) and right panels show corresponding H3K27Me3 signal in WT and mutant limbs across time. Lines indicate mean expression or H3K27Me3 signal across time for WT (solid line, circles) and mutant (dashed line, triangles), with biological replicates overlaid. See also Figures S22–S24, Data S10-S12.

### Eed loss alone is sufficient to produce temporal program redistribution

Since *Eed* is not the only chromatin modifier affected in the mutant limb, we sought to identify the core molecular defects attributable to the loss of PRC2 function. Thus, we used an orthogonal approach and applied WATER to RNA-seq data across 5 timepoints 0, 12, 24, 48, and 96 hours in WT and *Eed* CRISPR knockout (*Eed*-KO) mouse embryonic stem cells^45^. Both WT and Eed-KO cells were exposed to MEK and GSK3β inhibitors (“2i”) to block initiation of differentiation programs to study how loss of PRC2 function would alter the maintenance of stemness and onset of differentiation. In this context, minor spliceosome is normal, so the effects here are primarily owed to loss of Eed and PRC2 function. WATER applied to WT versus Eed-KO RNA-seq revealed that Eed itself undergoes dynamic expression changes across WT stem cell differentiation which is abolished in the Eed-KO (Fig. 5A, Data S13). Moreover, like the U11-null limb, many genes underlying molecular programs such as regulators of cell cycle and epigenetic regulators that were enriched in WT-0hr^Hi^ shifted to the Eed-KO-12hr^Hi^ or were absent (Fig. 5B-B’). Likewise, the GO term “*bone development*” enriched in the WT-48hr^Hi^ trajectory was absent in the mutant, whereas genes enriching for “*pattern specification*” were observed in the mutant-0hr^Hi^ instead of the WT-96hr^Hi^ (Fig. 5B-B’, Data S14). Specific genes such as *Bmp7* and *Msx2* that were affected in the U11 null limb were similarly affected in the Eed-KO cells (Fig. 5C-D, S14D’, S17E-G). Projecting genes in the WT trajectories onto the mutant showed distinct shifts in molecular programs regulating chromatin modification, RNA processing and DNA damage response (Fig. 5E-F). The most prominent shift in the absence of Eed was 498 genes in the WT-0hr^Hi^ that were NonDE in the Eed-KO (Fig. 5G) whereas others such as *Msx1* and *Dlx3* are ectopically expressed in the Eed-KO (Fig. 5H-I).

**Figure 5.**
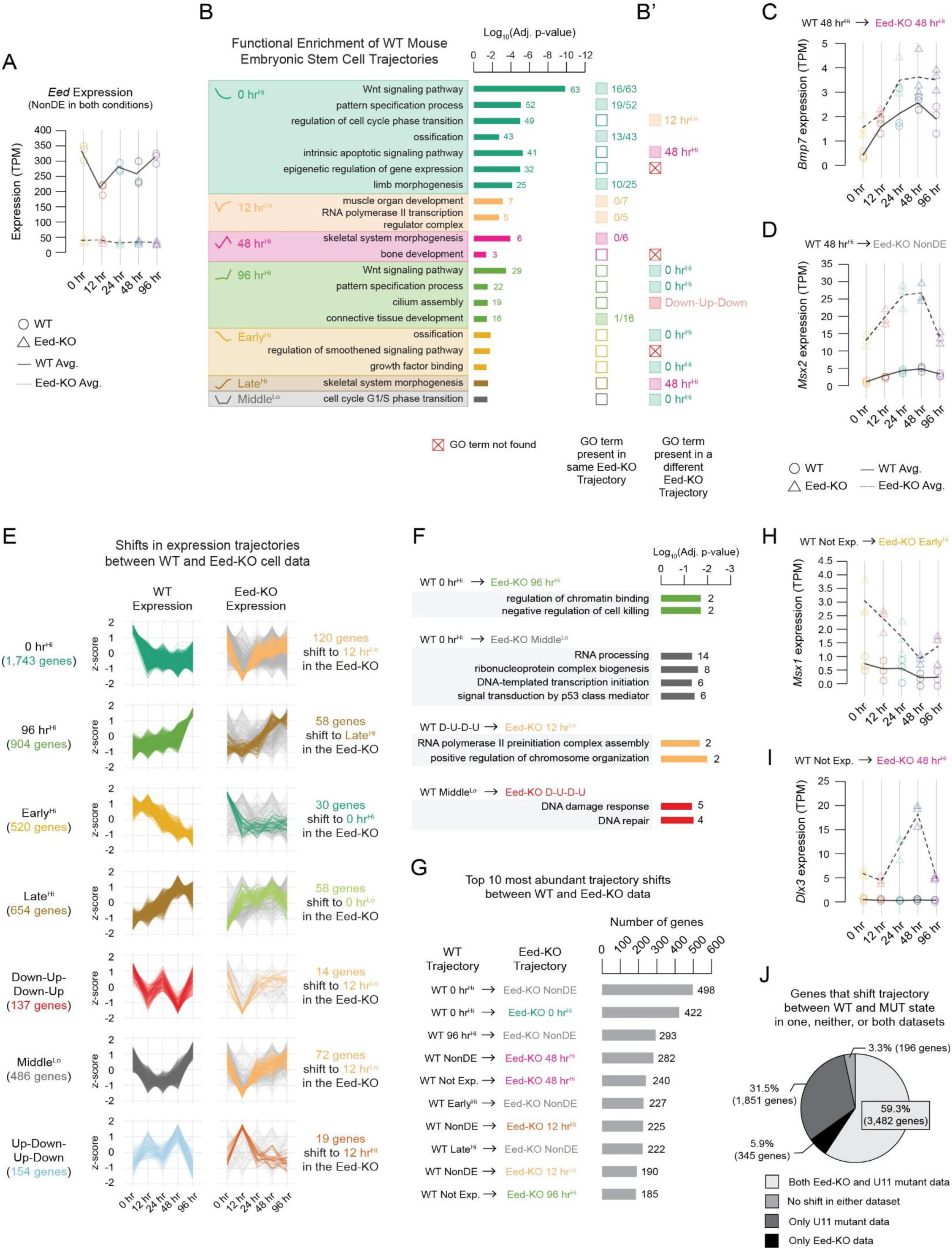
Loss of Eed in mouse embryonic stem cells results in shifts in transcriptional trajectories and functional programs consistent with patterns observed following U11 loss, related to. Figure 4. (A) Expression of *Eed* across time (0, 12, 24, 48, 96 hr) in WT and Eed-KO mouse embryonic stem cells. Points represent individual replicates; lines indicate mean expression (TPM). *Eed* expression is dramatically reduced in KO cells across all timepoints. (B) GO enrichment analysis of WT trajectories. Terms are grouped by trajectory. Bar length indicates significance (-log10 adjusted p-value), and numbers indicate gene count per term. (B’) Colored boxes indicate whether the same GO term is present in the corresponding Eed-KO trajectories or shifted to different temporal programs. (C-D) Representative expression trajectories illustrate altered temporal dynamics in Eed-KO cells. (C) *Bmp7* and (D) *Msx2* show shifted expression patterns in the Eed-KO relative to WT cells that are consistent with changes observed in U11 mutant forelimbs relative to WT. (E) For each WT trajectory (left), z-scored expression trajectories are shown in the Eed-KO (right) for the same gene set. The most frequently reassigned Eed-KO trajectory is highlighted in color, with the number of genes undergoing that shift indicated to the right; all other trajectories are shown in grey. Not all trajectories are shown. (F) GO enrichment analysis for subsets of WT genes that shift to alternative trajectories in the Eed-KO. Bars represent significantly enriched GO terms (BP, CC, MF; adjusted p < 0.05), with bar length indicating the log_10_(adjusted p-value) and numbers denoting the number of genes underlying enrichment of each term. (G) Top trajectory shifts ranked by the number of genes changing between WT and Eed-KO data. (H-I) Examples of genes not expressed in the WT but aberrantly activated in Eed-KO cells at specific timepoints. (H) *Msx1* and (I) *Dlx3* illustrate activation in KO cells consistent with changes observed in U11 mutant forelimbs relative to WT. (J) Proportion of genes exhibiting trajectory shifts between WT and mutant (U11-cKO or Eed-KO) datasets. Pie chart summarizes genes that shift in both datasets, only one dataset, or neither. See also Data S13-S14.

The 0hr^Lo^ trajectory in Eed-KO was strongly enriched for embryonic skeletal system development (adj. p-value = 8.5 × 10⁻^3^, n = 9 genes) and pattern specification (adj. p-value = 5.2 × 10⁻^3^, n = 18 genes), indicating that PRC2 normally suppresses Hox and skeletal identity loci via H3K27me3 at the earliest timepoints (Data S14). Canonical PRC2 targets including *Hoxb4*, *Hoxc4*, *Hoxc6*, *Alx3*, *Rdh10*, and *Wnt9a* populated this gate-failure trajectory, consistent with previous reports that H3K27me3-dependent Hox cluster suppression is abolished by Eed loss^44^. Despite differences in the precise trajectory redistribution between the U11-null limb and Eed-KO stem cells, 59.3% of genes showed a trajectory shift in both systems (Fig. 5J, Data S13), revealing the molecular programs regulated in part through the Eed–PRC2 axis in the U11-null limb.

### Minor spliceosome inhibition delays progenitor maturation and impairs chondrogenesis

The chromatin-transcription uncoupling observed in Figure 5 predicted that progenitor cells in the mutant limb would fail to execute timely lineage transitions. During normal limb development, sequential chromatin gating progressively restricts progenitor cell competence and enables orderly progression from proliferative progenitors to fate-committed skeletal lineages^46–48^. To determine whether epigenetic destabilization manifests at the cellular level, we performed single-cell RNA-seq of WT and U11-null forelimbs at E10.5, E11.5, and E12.5 (Fig. 6A, S25A). Integration of 48,632 WT and 63,742 mutant cells revealed 22 transcriptionally distinct clusters (Fig. 6A). Cluster identities were validated using canonical lineage markers (Fig. S25B, Data S15), and timepoint-specific UMAP projections demonstrated preserved cluster identities but altered occupancy across developmental stages (Fig. S25A, C). Quantification of major cell-type proportions revealed sustained abundance of limb progenitor cells (LPCs) and reduced representation of chondroblast populations in mutant limbs (Fig. 6B–D). Timepoint-resolved composition analysis confirmed delayed depletion of progenitors and impaired expansion of skeletal lineages in mutant samples (Fig. S25C). To determine whether spatial patterning programs were preserved, we examined domain-specific marker expression (Fig. S26A-B, Data S16). Feature projections demonstrated appropriate expression of anterior-posterior and proximal-distal markers (Fig. S26C), indicating that MiS inhibition perturbs the temporal deployment of cell states rather than spatial patterning programs. This was consistent with our previous report that expression of patterning genes was conserved in the mutant forelimb by whole mount in situ^20^. Interrogation of *Col2a1* expression, a chondroblast marker, across single-cell clusters revealed a marked reduction in distal chondroblasts corresponding to cluster 19 (Fig. 6E, S26A). Section in situ hybridization revealed *Col2a1* expression in both WT and mutant stylopods, whereas distal autopodial signal appeared nearly absent in mutant limbs at E12.5 (Fig. 6F). Spatial annotation identified cluster 17 as proximal chondroblasts and cluster 19 as distal chondroblasts (Fig. 6G, S26A–A’), highlighting selective depletion of distal skeletal progenitors. Expression of Sox9, a marker of osteochondroprogenitor cells, revealed defective condensation dynamics in mutant limbs (Fig. 6H–I). Whereas WT limbs exhibited bifurcation of the radius and ulna by E11.5 and clear digit condensations by E12.5, mutant limbs failed to undergo zeugopodial bifurcation and lacked distal digit condensations, with expansion of the distal Sox9-negative territory (Fig. 6I). Quantification confirmed reduced Sox9-positive chondrogenic area at early stages and increased distal ectoderm-to-Sox9 gap distance, in agreement with delayed initiation of distal condensation (Fig. S27A–B, Table S4). Three-dimensional reconstruction confirmed reduced distal condensations and zeugopodial defects (Fig. S27C–D). Additional in situ analyses for *Col1a1* and *Col10a1,* which mark tendon/osteoblast progenitors and maturing chondroblasts, respectively; demonstrated impaired maturation of the osteogenic lineage in addition to defective chondrogenesis (Fig. S27E–F). Conversely, *Msx1*, a marker of naive limb progenitor cells and a locus previously identified as chromatin-expression rewired gene (Fig. 4), remained abnormally elevated and spatially expanded in mutant limbs (Fig. 6J–K), indicating persistence of early progenitor programs^49^. Consistent with progenitor maintenance and impaired chondrogenesis, pseudotime analysis across integrated progenitor populations revealed enrichment of mutant cells in transcriptionally earlier states relative to WT (Fig. 6L), supporting delayed lineage progression following MiS inhibition.

**Figure 6.**
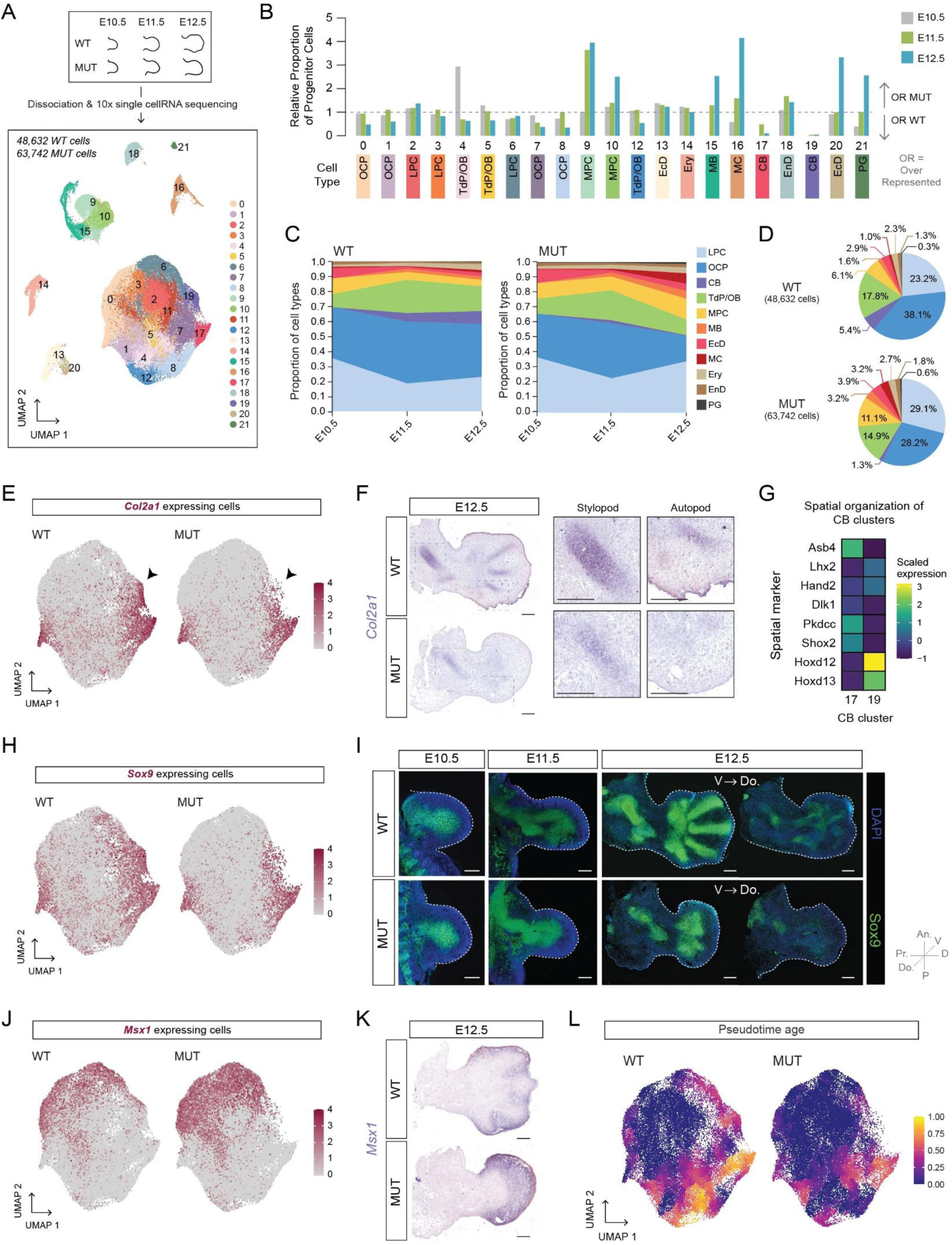
Minor spliceosome inhibition results in accumulation of naïve limb progenitor cells and delayed chondrogenesis. (A) Experimental design and integration of single-cell RNA sequencing data from WT and U11-mutant (MUT) forelimbs at E10.5, E11.5, and E12.5. UMAP visualization shows integrated clustering of 48,632 WT cells and 63,742 MUT cells, colored by cluster identity. (B) Relative proportion of progenitor cells across clusters at each developmental timepoint. Bars show enrichment of cluster in WT (< 1) or mutant (> 1) limbs, with the dashed line indicating equal representation. (C) Proportion of major limb cell types across developmental time in WT and mutant forelimbs. Stacked area plots reveal sustained abundance of limb progenitor cell (LPC, light blue) populations and lack of chondroblasts (CB, purple) in mutant limbs relative to WT. (D) Overall distribution of major cell types in WT and mutant datasets, shown as pie charts. (E) Feature plots showing *Col2a1* expression in WT and mutant forelimbs. Arrows highlight reduced *Col2a1* expression in a subset of chondrocytes that correspond to cluster 19. (F) In situ hybridization for *Col2a1* at E12.5 in WT and mutant forelimbs. Insets highlight stylopod and autopod regions, revealing a pronounced reduction in autopodial chondrocytes in mutant limbs. (G) Heatmap depicting scaled expression of spatial marker genes across chondroblast clusters, identifying cluster 17 as proximal chondroblasts (*Pkdcc* and *Shox2* expressing) and cluster 19 as distal chondroblasts (*Hoxd12* and *Hoxd13* expressing). (H) Feature plots showing *Sox9* expression in WT and mutant limbs. (I) Immunofluorescence for Sox9 at E10.5, E11.5, and E12.5 in WT and mutant forelimbs. Dotted lines denote limb boundaries. Two representative sections are shown for E12.5 limbs to highlight skeletal differences along the dorsal (Do.) to ventral (V) axis. (J) Feature plots showing *Msx1* expression in WT and mutant limbs. (K) In situ hybridization for *Msx1* at E12.5 in WT and mutant forelimbs. (L) Pseudotime analysis of WT and mutant progenitor cells projected onto the integrated UMAP, revealing the mutant limb is composed of younger cells relative to the WT. Scale bars, 200 μm. An.; anterior, P; posterior, D; distal, Pr.; proximal, Do.; dorsal, V; ventral, OCP; osteochondroprogenitor cell, Tdp/OB; tendon progenitor cell/osteoblast progenitor, MPC; muscle progenitor cell, EcD; ectoderm, MB; myoblast, MC; myeloid cells, Ery; erythroid cells, EnD; endothelial cells, PG; peripheral glia. See also Figures S25–S28, Data S15-S17.

### Trajectory shifts reflect both cellular composition and intrinsic transcriptional rewiring

We directly tested whether WATER-defined expression trajectory shifts reflect altered cellular abundance or intrinsic transcriptional reprogramming. Applying WATER to gene-specific cellular proportions (Fig. S28A–B, Data S17) revealed that some genes, such as *Acan*, exhibit concordance between bulk expression and cellular abundance (Fig. S28C). However, key redistributed genes such as *Msx1* displayed preserved abundance kinetics but increased per-cell expression in the mutant (Fig. S28D–E), demonstrating that the observed bulk expression shift cannot be attributed to changes in cell composition alone. Systematic intersection analysis demonstrated that approximately 72.8% of mutant trajectory shifts cannot be explained by cellular composition changes alone (Fig. S28F–I). Together, these data demonstrate that minor spliceosome inhibition drives both altered intrinsic transcriptional temporal shifts and cell-state composition. Persistence of early transcriptional states, failure to extinguish progenitor identity, and impaired activation of chondrogenic differentiation collectively provide a cellular basis for the chromatin and transcriptomic redistribution observed at the tissue level. Thus, U11 loss disrupts developmental chronology, converting coordinated lineage progression into temporally desynchronized maturation that is a form of cellular heterochrony in which cell states are displaced in time relative to the organism’s developmental clock. In essence, the crosstalk between various developing systems in the limb becomes asynchronous thereby compromising limb development.

### p53 ablation alleviates checkpoint-mediated apoptosis and partially rescues micromelia

Through single-cell analysis, we identified cluster 11 as a distinct population present almost exclusively in mutant limbs (Fig. 7A–A’). Marker analysis indicated that these cells correspond to distal LPCs exhibiting a stress-associated transcriptional profile (Fig. 7B, Data S15). Cluster 11 cells accumulated in mutant limbs across developmental time, with absolute cell numbers elevated relative to WT at each stage (Fig. 7A–A’), consistent with our previous reports of apoptosis upon U11 loss^20^. These cluster 11 cells therefore represent a mutant-specific progenitor state defined by both spatial identity (distal LPC markers) and cellular stress, providing a cellular readout of the consequences of splicing-driven temporal miscoordination at single-cell resolution. Violin plots revealed enrichment of canonical p53-responsive cell death transcripts including *Pmaip1*, *Cdkn1a*, *Phlda3*, and *Ccng1* (Fig. 7B), consistent with a checkpoint-mediated apoptotic program. To assess the lineage fate of these stress-associated LPCs, we reconstructed developmental trajectories using Monocle3 (Fig. 7C). In WT limbs, LPCs transitioned through osteochondroprogenitor cell intermediates toward chondroblast lineages along continuous trajectories. In contrast, mutant limbs exhibited fragmented trajectories with reduced representation of osteochondroprogenitor cells and distal chondroblast states. Focused analysis revealed disruption of the trajectory connecting cluster 11 progenitors through cluster 7 osteochondroprogenitor cells to cluster 19 distal chondroblasts (Fig. 7C–D). These observations indicate that LPCs undergoing apoptosis in mutant limbs would normally contribute to distal skeletal progenitors and autopodial chondroblasts, directly linking the progenitor loss to the selective absence of distal skeletal elements observed morphologically. To test whether checkpoint activation contributes to the phenotype, we ablated *Trp53* in U11-null mice (Fig. S29A–C). TUNEL staining at E12.5 showed high levels of apoptotic cells in U11 mutants, whereas U11-p53 double mutants displayed minimal cell death, confirmed by quantification (Fig. 7E–F). Skeletal preparations at postnatal day 0 revealed that while U11 single mutants exhibited severe hypoplasia and reduced skeletal complexity, U11-p53 double mutants displayed zeugopod bifurcation and elaboration of the autopod (Fig. 7G, S29D). Quantification of total limb length and long bone width demonstrated significant increases in stylopod and zeugopod measurements in double mutants relative to U11 single mutants, and measurement of ossified limb length showed partial recovery of mineralized regions (Fig. 7H–I, S29D–E), although values remained below WT levels, consistent with partial rather than complete rescue. RT-PCR analysis of representative MIGs including *Eed*, *Lsm5*, *Med23*, *Kansl2*, *Ints7*, and *Xrcc5* demonstrated persistent splicing defects in U11-p53 mutant limbs (Fig. 7J). Thus, p53 loss alleviates apoptosis and partially restores skeletal growth without correcting minor spliceosome-dependent molecular defects, demonstrating that checkpoint activation amplifies rather than initiates developmental failure.

**Figure 7.**
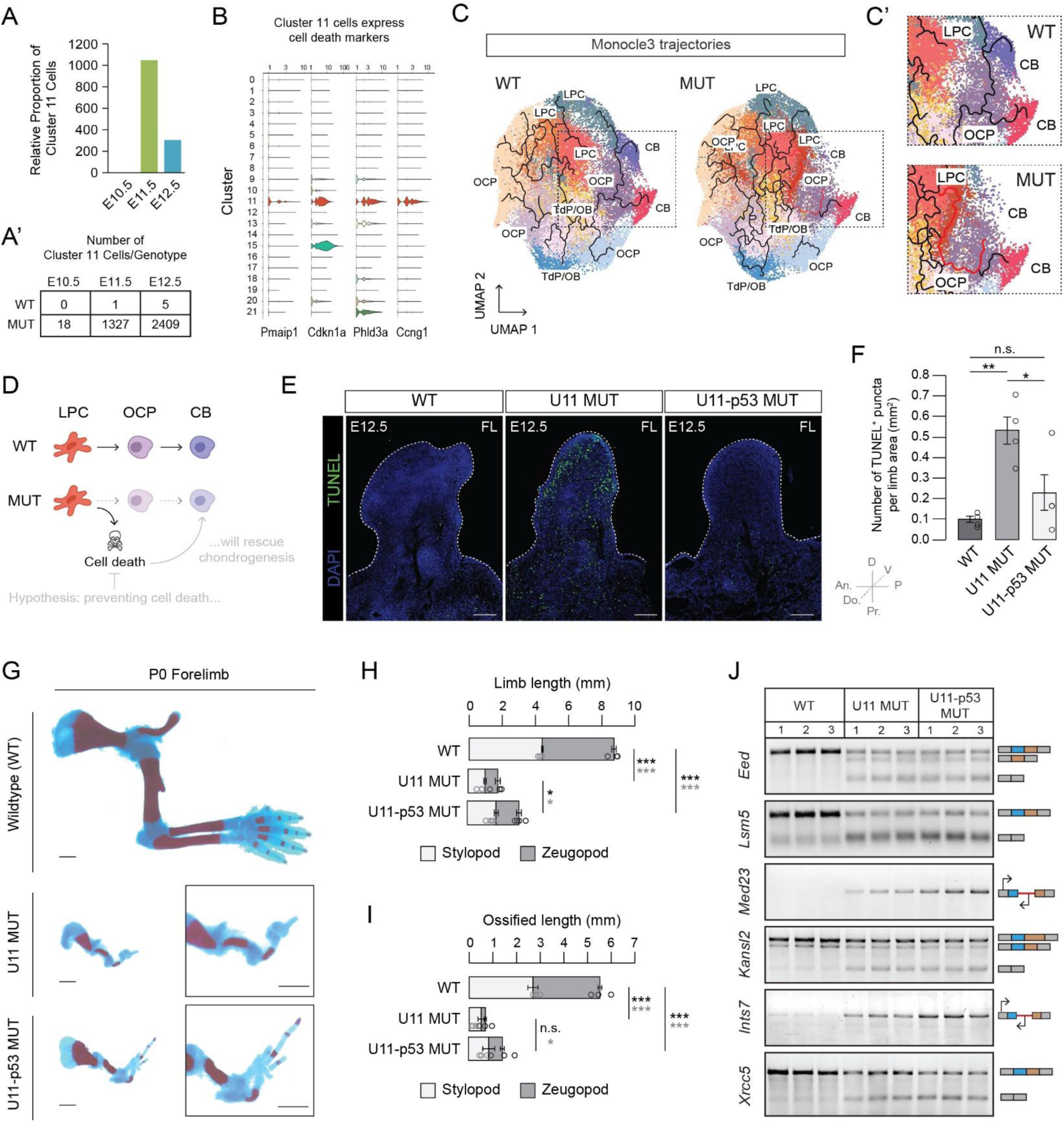
p53-dependent cell death contributes to impaired chondrogenesis and morphological defects in the developing forelimb following U11 loss. (A) Relative proportion of cells assigned to cluster 11 across time (E10.5-E12.5). (A’) Number of cluster 11 cells per genotype at each timepoint. (B) Violin plots showing expression of cell death-associated genes (*Pmaip1, Cdkn1a, Phlda3,* and *Ccng1*) across clusters, highlighting enrichment in cluster 11. (C) Monocle3 trajectories of single-cell transcriptomes from WT and mutant forelimbs, annotated by cell type including limb progenitor cells (LPC), osteochondroprogenitor cells (OCP), tendon progenitor cells/osteoblast progenitors (Tdp/OB), and chondroblasts (CB). (C’) Zoomed views of clusters 7, 11, 17, and 19. The red line indicates a Monocle3 trajectory connecting cluster 11 (dying LPCs) to cluster 19 (distal CBs) through cluster 7 (OCPs), which are reduced in mutant limbs. (D) Model illustrating the proposed relationship between cell differentiation and death in the WT and mutant limbs. In MUT limbs, increased cell death is hypothesized to reduce progenitor populations that would normally contribute to distal cartilage formation, thereby delaying cartilage formation. (E) Representative images of TUNEL staining of WT, U11 mutant, and U11-p53 mutant E12.5 forelimb sections. DAPI (blue) marks nuclei while TUNEL-positive puncta indicate apoptotic nuclei (green). Dashed lines outline the limb boundaries. An.; anterior, P; posterior, D; distal, Pr.; proximal, Do.; dorsal, V; ventral. (F) Quantification of TUNEL-positive puncta per limb area across genotypes. Each point represents a biological replicate; bars indicate mean + SEM. Statistical significance was determined by a one-way ANOVA followed by post-hoc Tukey’s test. (G) Skeletal preparations of postnatal day 0 (P0) forelimbs from WT, U11 mutant, and U11-p53 mutant mice, illustrating differences in overall limb size and skeletal complexity. (H) Quantification of total limb length at P0, separated into stylopod and zeugopod measurements. (I) Quantification of ossified limb length at P0, separated into stylopod and zeugopod measurements. For both H and I, each point represents a biological replicate, and bars indicate mean + SEM. P values in H and I indicate the results of a one-way ANOVA followed by post-hoc Tukey’s test. * p < 0.05, ** p < 0.01, *** p < 0.001, n.s., not significant. (J) RT-PCR analysis of representative minor intron-containing genes (*Eed*, *Lsm5*, *Med23*, *Kansl2*, *Ints7*, and *Xrcc5*) across WT, U11 mutant, and U11-p53 mutant forelimbs. Schematics to the right indicate the type of alternative splicing or primer design used to assess intron retention. RT-PCR analysis demonstrates that p53 loss does not rescue minor spliceosome-dependent splicing defects. See also Figure S29, Table S5.

## Discussion

A central insight from this study is that tissue development is governed not only by which genes are expressed, but by when their expression kinetics constitute programs that are deployed, transitioned, and terminated in a defined sequence. By resolving temporal program topology, WATER reveals that developmental progression depends on the precise order of program execution rather than the steady accumulation of transcriptional changes at any given timepoint. Disruption of this temporal architecture, even in the absence of wholesale transcriptional loss, is sufficient to derail morphogenesis. This framework helps resolve a longstanding paradox in developmental genetics: many mutants retain expression of key developmental regulators yet exhibit profound morphological defects. Preservation of gene expression does not imply preservation of developmental logic. Failure arises when programs are deployed outside their appropriate temporal context, and the mutant limb proceeds along an irreversible alternative developmental path that is only revealed through time-resolved analysis. These observations argue that temporal topology should be treated as a primary dimension of developmental regulation alongside expression level and spatial domain. Classical heterochrony studies have documented how changes in the timing of individual developmental events such as the duration of Shh expression, the onset of Hox codes, the kinetics of retinoic acid gradients drive morphological evolution across species^15–17,29^. The present work demonstrates that equivalent timing disruptions at the level of coordinated molecular programs can arise within a single genetic perturbation and constitute a primary disease mechanism, extending the concept of heterochrony from evolutionary morphology into developmental pathology. Minor spliceosome inhibition disrupts this architecture through MIGs that are disproportionately enriched for the regulatory nodes such as chromatin modifiers, transcription factors, cell cycle regulators, and RNA processing factors, which are required to terminate early programs and initiate later ones. Aberrant splicing of these genes creates a developmental bottleneck in which early biosynthetic and proliferative programs fail to terminate while differentiation programs are displaced into later windows.

The selective enrichment of chromatin regulators among the earliest mis-spliced MIGs is mechanistically significant: it places epigenetic misregulation at the apex of the cascade, upstream of transcriptional redeployment. Mis-splicing of *Eed* compromises PRC2 function and reduces H3K27me3 deposition across the developing limb, decoupling chromatin state from transcriptional timing. That *Eed* loss alone is sufficient to recapitulate core features of this redistribution in stem cells with intact minor spliceosome function establishes PRC2 as one primary effector arm of the phenotype. Moreover, in the Eed-KO the p53-checkpoint activation happens earlier than in the U11-null limb as the MIGs involved in DNA repair axis are intact whereas in the U11-null limb these MIGs are mis-spliced and p53 checkpoint activation is delayed. Crucially, the partial rescue of distal limb structures by Trp53 ablation confirms that blocking cell death allows patterning programs to operate longer but cannot restore the temporal architecture that splicing defects have already disrupted. This does not exclude independent contributions from *Chd4*, *Actl6a*, or other chromatin MIGs; those factors likely contribute additional dimensions of the phenotype that are now accessible to investigation using WATER.

At the cellular level, disruption of temporal program architecture manifests as cellular heterochrony. Single-cell analysis reveals that progenitor populations persist at stages when they should have transitioned, osteochondroprogenitor and chondroblast lineages are selectively reduced, and approximately 72.8% of trajectory shifts reflect intrinsic transcriptional rewiring rather than changes in cell composition alone. Spatial patterning programs remain largely intact despite this widespread temporal desynchronization, suggesting that developmental systems buffer spatial identity while temporal sequence is disrupted. It is this buffering capacity that collapses as progressive progenitor attrition through the p53 checkpoint removes the cellular substrate required to elaborate distal skeletal elements. This mechanistic pathway, from splicing defect through epigenetic gating failure through temporal program redistribution through cellular heterochrony to morphological failure, represents a disease mechanism distinct from classical loss-of-function models and invisible to stage-matched transcriptomic analysis.

More broadly, these results position temporal program architecture as a primary regulatory dimension of development that is susceptible to disruption as a pathogenic mechanism independently of gene identity or expression level. Many developmental diseases present with dynamic, stage-specific transcriptional signatures that resist simple categorical interpretation; WATER provides a framework to ask whether such phenotypes reflect program mistiming rather than program loss. The minor spliceosome, long treated as a constitutive processing system, emerges here as an active regulator of epigenetic gating and developmental timing through its control of a specific set of chromatin-regulatory targets. We propose that trajectory-based approaches that treat time as a primary analytical dimension will be essential tools for understanding how developmental timing is encoded and how its disruption produces morphological disease. The many open questions in limb heterochrony: 1) the molecular basis of segment-specific differences in chondrogenesis timing, 2) the cell-intrinsic and extrinsic regulators of Shh duration, and 3) the anterior-posterior developmental delay in marsupials^16,18,49,50^. These issues are now approachable using WATER, which can quantify program-level temporal shifts in any system where bulk temporal RNA-seq and other omics data are available.

## Limitations

This study focuses on mis-splicing of *Eed* and the disruption of PRC2 to illustrate chromatin gating of molecular programs in limb development. However, other chromatin modifiers that are MIGs are also disrupted. Further analysis of these factors is appropriate for future studies. Regarding the partial rescue with p53 ablation, we propose that the cells rescued correspond to the loss of osteochondroprogenitors identified in scRNA-seq that show activation of the p53 checkpoint. The identity of the rescued cell type does not change the overall interpretation that we rescue cell death of a specific cell type without rescuing the splicing defect, which underpins the partial rescue of micromelia. Conditional deletion via *Prrx1-*Cre achieves broad but not complete recombination within the limb mesenchyme, and mosaic retention of Rnu11 in a subset of cells may attenuate the severity of the observed phenotype. However, the consistent and reproducible nature of the molecular and morphological phenotypes across biological replicates argues that mosaicism does not fundamentally alter our conclusions. WATER is further constrained by its reliance on bulk temporal RNA-seq data, and performance may be reduced in systems with very few timepoints or where temporal and spatial variation are strongly confounded. Future integration of WATER with spatial transcriptomics may address these limitations and expand its applicability to systems where cell-type-specific spatiotemporal dynamics are of primary interest.

## Acknowledgement

The authors would like to thank Dr. Clifford Tabin, Dr. Vanessa Aguiar-Pulido, Dr. Anouk Olthof, Dr. Marybeth Baumgartner, and Dr. Leighton Core for their insightful comments that allowed us to improve the manuscript; Dr. Bo Reese from the University of Connecticut’s Center for Genome Innovation for assistance with RNA-seq, CUT&RUN, and single cell RNA-seq; and the Computational Biology Core within the Institute for Systems Genomics at the University of Connecticut for access to the high-performance computing cluster resources. This project has received funding from the NINDS R01NS102538, Prostate Cancer Foundation Challenge Award 2022, and UConn CLAS Academic Themes Award.

## Author Contributions

Conceptualization – R.N.K.; Methodology – S.M.S. and R.N.K.; Investigation – S.M.S., A.R.B., K.D.D., K.O.A., K.N.G., T.K., I.S., N.C., T.L., and R.N.K.; Software – S.M.S. and R.N.K.; Resources – R.N.K.; Writing – original draft – S.M.S. and R.N.K.; Writing – review and editing – S.M.S., A.R.B., K.D.D., and R.N.K.; Supervision – R.N.K.; Funding acquisition – R.N.K.

## Star Methods

Detailed methods are provided in the online version of this paper and include the following:

- Resource Availability
  - Lead Contact
  - Materials Availability
  - Data and code availability
- Experimental Model and Subject Details
  - Mouse strains
  - Animal husbandry
- Method Details
  - RNA isolation and cDNA preparation
  - Single and duplex RT-PCR
  - RNA probe preparation
  - Section in situ hybridization
  - Whole mount in situ hybridization
  - Immunofluorescence
  - TUNEL assay
  - Skeletal preparation
  - Total RNA sequencing
  - Alignment, quantification, and DE analysis
  - Static DE analysis and PCA
  - WATER trajectory analysis
    - Preparing data for WATER
    - Initial trajectory analysis
    - Initial correlation-based trajectory refinement
    - Iterative trajectory refinement
    - Final trajectory assignment
    - Leave-One-Out (LOO) stability analysis
    - Comparative analysis of WT and MUT data
  - Alternative splicing analysis
  - ORF prediction and AlphaFold modeling
  - CUT&RUN profiling and data processing
  - CUT&RUN signal visualization
  - Single-cell RNA sequencing
  - scRNA-seq data filtering and processing
  - scRNA-seq data integration and analysis
  - Pseudotime and Monocle3 analysis
- Quantification and Statistical Analysis
  - Filtering alternative splicing data
  - Gene ontology enrichment analysis
  - Densitometric analysis of RT-PCR
  - 3D reconstruction from serial sections
  - Section staining quantification
  - Quantification of limb surface area
  - Quantification of skeletal elements

## Resource availability

### Lead Contact

Further information and requests for resources and reagents should be directed to and will be fulfilled by the Lead Contact, Rahul Kanadia.

### Materials Availability

Primer sequences used to generate custom riboprobes are provided in Table S5.

### Data and code availability

- Previously published E10.5 and E11.5 total RNA-seq data are available under the GEO accession number GSE146424.
- Bulk RNA-Seq, CUT&RUN-Seq, and single-cell RNA-Seq data generated in this study have been deposited in NCBI’s Gene Expression Omnibus (GEO) and will be made publicly available upon publication. The dataset includes total RNA-Seq at E12.5 and E13.5, CUT&RUN profiling of IgG and H3K27Me3 at E10.5-E12.5, and single-cell RNA-Seq data at E10.5-E12.5. All datasets include both WT and mutant forelimb samples.
- Bulk RNA-seq from WT and Eed-KO mouse embryonic stem cells was previously published and are available under the GEO accession number GSE237656.
- Raw data and unprocessed images have been deposited in Mendeley Data and will be made publicly available upon publication.
- All custom scripts and code used in this study have been deposited to GitHub and will be made publicly accessible upon publication.
- Any additional information required to reanalyze the data reported in this paper is available from the Lead Contact upon request.

### Experimental Model and Subject Details

#### Mouse Strains

The generation of *Rnu11^Flx/Flx^*, *Trp53^Flx/Flx^, Ai9-Rosa26-tdTomato, and Prrx1-Cre^+^* mice has been described previously. *Rnu11^WT/Flx^*::*Prrx1-*Cre^+^ males were crossed with *Rnu11^Flx/Flx^* females to generate U11 conditional mutant mice (MUT; *Rnu11^Flx/Flx^*::*Prrx1-*Cre^+^) and WT littermate controls (WT; *Prrx1-*Cre^-^). Initial characterization of the *Rnu11^Flx/Flx^::Prrx1-Cre^+^* line was previously reported^20^. To generate U11-p53 double mutant mice, *Rnu11^WT/Flx^*::*Trp53^WT/Flx^::Prrx1-*Cre^+^ males were crossed with *Rnu11^Flx/Flx^::Trp53^Flx/Flx^* females. All mice were maintained on a C57BL/6 background.

#### Animal Husbandry

All animal procedures were conducted in compliance with protocols approved by the University of Connecticut Institutional Animal Care and Use Committee (IACUC), in accordance with the U.S. Public Health Service Policy on laboratory animal care. Researchers, animal care staff, and veterinary personnel monitored all mice daily for general health and activity. Experimental animals included both sexes. Littermate controls were used for all sequencing experiments and when possible were also used for all molecular validation. For embryonic harvests, tail snips from embryos were used for genotyping. For embryonic experiments, embryonic day (E) 0.5 was designated as noon on the day a vaginal plug was observed. For postnatal animals, genomic DNA was collected via ear punches at postnatal day 10. Primers used for genotyping of each mouse strain can be found in Table S5. Generation of the *Rnu11^Flx/Flx^*line and validation was previously reported^35^. Downregulation of the U11 snRNA following *Prrx1*-Cre-mediated recombination was previously reported^35^.

### Method Details

#### RNA isolation and cDNA preparation

Forelimbs from E10.5, E11.5, E12.5, and E13.5 WT and mutant embryos were collected for RNA isolation (n=3 biological replicates per timepoint and genotype). Left and right forelimbs from a single embryo were pooled into 150 microliters of TRIzol reagent (Thermo Fisher Scientific) and homogenized prior to storage at -80°C. Total RNA was extracted using the Direct-zol RNA Miniprep Kit (Zymo Research) according to manufacturer instructions. RNA concentration and purity were assessed using a NanoDrop One Spectrophotometer (Thermo Fisher Scientific) and visualized on a 1% agarose gel prior to cDNA synthesis.

To assess RNA integrity before sequencing, 100-300 ng total RNA from each sample was used for cDNA synthesis using a customized protocol described previously^72^. RNA was incubated with a 1:1 mix of oligo-dT (Thermo Fisher Scientific) and random hexamers (Thermo Fisher Scientific) at 65°C for 15 minutes, followed by 25°C for 15 minutes. A master mix containing 5x First Strand Buffer, dTT, RNase inhibitor (Roche), dNTPs (Promega), and M-MLV Reverse Transcriptase (Thermo Fisher Scientific) was added, and reactions were incubated at 37°C for 50 minutes, followed by 70°C for 15 minutes before storage at -20°C. cDNA quality was assessed by RT-PCR amplification of *Gapdh* to confirm sample integrity (Table S5). cDNA was used for subsequent RT-PCR validation.

#### Single and duplex RT-PCR

RT-PCR analyses were performed using the GO Taq Flexi DNA Polymerase system (Promega) according to the manufacturer’s instructions. Products were amplified using standard PCR conditions with an annealing temperature of 58°C and 28-32 amplification cycles, depending on the target transcript. Primers used for the detection of gene expression, alternative splicing events, and intron retention are listed in Table S5. For duplex RT-PCR reactions, *Gapdh* primers were included as an internal control and were diluted at a 1:4 ratio relative to the target primer sets to prevent preferential amplification. PCR products were resolved on a 2% agarose gel and visualized using ethidium bromide under UV illumination.

#### RNA probe preparation

DIG-labeled RNA probes were generated using a PCR-based in vitro transcription method adapted from previously described protocols^73,74^.Gene-specific primers were designed to amplify 100-800 bp fragments. A T7 promoter sequence with a linker (5’ – CAGTGAATTGTAATACGACTCACTATAGG – 3’) was appended to the 5’ end of the reverse primer to enable in vitro transcription. Probe templates were amplified from cDNA using Platinum High-Fidelity Taq polymerase (Thermo Fisher Scientific). PCR products were resolved on a 2% agarose gel, excised, and purified using the QIAquick Gel Extraction Kit (Qiagen). Purified products were then utilized as input to a second PCR using the T7 promoter-containing reverse primer to generate transcription templates. DIG-labeled antisense RNA probes were synthesized using T7 RNA polymerase (Roche) and DIG RNA labeling mix (Roche) in an in vitro transcription reaction. Following transcription, DNA templates were degraded using DNase I (Invitrogen) digestion. Purified RNA probes were precipitated using LiCl and glycogen (Invitrogen) followed by resuspension in DI formamide (Millipore-Sigma) and storage at -20°C until use.

#### Section in situ hybridization

ISH was performed as previously described with minor modifications^72^. Embryonic forelimbs were fixed overnight in 4% paraformaldehyde in PBS at 4°C, washed with PBS, and incubated overnight in 30% sucrose followed by a final overnight incubation in a sucrose/OCT mixture. Samples were embedded in OCT and sectioned at 10 μm using a cryostat. Sections were stored at -80°C until use. Prior to staining, sections were brought to room temperature (RT), rehydrated in PBS/DEPC, and permeabilized with PBT (PBS containing Tween-20). Sections were treated with 25 μg/mL Proteinase K (Roche) and post-fixed in 4% paraformaldehyde. Following several PBT washes slides were incubated in an acetylation mix and further washed with PBT prior to the addition of DIG-labeled RNA probes in hybridization buffer. Sections were allowed to hybridize with the probe overnight at 65°C and were subsequently washed in a formamide/SSC buffer at 65°C after which sections were incubated with RNase A (Roche) at 37°C. Following several SSC washes at 65° and MABT washes at RT, slides were incubated in blocking solution with serum for an hour. Blocking solution was replaced with fresh that also contained anti-DIG alkaline phosphatase antibody (1:2500, Roche) and incubated overnight at 4°C. After extensive MABT washes, signal was detected using standard alkaline phosphatase colorimetric detection using NBT and BCIP. After developing, slides were fixed with 4% paraformaldehyde, washed with PBS, and mounted in gelvatol. White-light images were acquired using a Keyence BZ-X710 fluorescence microscope.

#### Whole mount in situ hybridization

WISH was performed as previously described with minor modifications^3^. Embryos at E10.5-E13.5 were fixed overnight in 4% paraformaldehyde in PBS at 4°C, dehydrated through a methanol/PBS series, and stored in 100% methanol at -20°C until processing. Prior to hybridization, embryos were rehydrated through a methanol/PBT series and treated with 10 μg/mL Proteinase K (Roche) for 5-17 minutes, depending on the probe. For Fgf8 staining, Proteinase K treatment was omitted. Embryos were post-fixed in 4% paraformaldehyde with 0.1% glutaraldehyde, washed in PBT, and incubated with hybridization buffer for 1 hour at 70°C. DIG-labeled RNA probes were denatured at 70°C for 30 minutes and hybridized overnight (12-16 hours) at 70°C. Following hybridization, embryos were washed in formamide/SSC buffers at 65-70°C and then equilibrated in TBST. Embryos were blocked in TBST containing blocking reagent (Roche) and heat-inactivated sheep serum before incubation with anti-DIG alkaline phosphatase antibody (1:2500) overnight at 4°C. After extensive TBST washes, signal was detected using standard alkaline phosphatase colorimetric detection using NBT and BCIP. Staining was performed on a minimum of two biological replicates per timepoint and genotype. Samples were imaged using an Olympus SZX16 UV dissecting microscope equipped with a top-mounted camera that allows for both white light and fluorescent imaging.

#### Immunofluorescence

IF for Sox9 was performed as previously described with minor modifications^35^. Embryonic forelimb sections were prepared as previously described for section in situ hybridization and stored at -80°C until use. Prior to staining, sections were brought to RT, rehydrated in PBS, and permeabilized with PBT. Samples were then blocked in blocking solution containing serum and blocking reagent before incubation with Sox9 primary antibody (Millipore-Sigma) overnight at 4°C. Sections were washed in PBT and incubated with fluorescent secondary antibodies for 3 hours at RT. Both primary and secondary antibodies were diluted to 1:500 prior to use. After washing in PBT, samples were mounted using ProLong Gold antifade reagent (Thermo Fisher Scientific), stained with DAPI to visualize nuclei, and imaged using a Keyence BZ-X710 fluorescence microscope at 10x and 20x with automatic settings for signal intensity and image stitching. For 3D reconstruction, serial cryosections of 25 μm thickness were collected for n=1 WT and mutant limb bud per slide set (total n=3) and reconstructed as described in the Quantification and Statistical Analysis section.

#### TUNEL assay

Detection of apoptotic nuclei was performed using the In Situ Cell Death Detection Kit, TMR red (Roche) according to the manufacturer’s instructions. Embryonic forelimb sections were prepared as previously described for section in situ hybridization and stored at -80°C until use. Prior to staining, sections were brought to RT, rehydrated in PBS, and permeabilized with PBT prior to incubation with the TUNEL reaction mixture for 1 hour at RT. Following labeling, sections were washed in PBS and stained with DAPI to visualize nuclei. Slides were mounted using ProLong Gold antifade reagent and imaged using a Keyence BZ-X710 fluorescence microscope. All TUNEL images have been deposited in the Mendeley Data repository associated with this study.

#### Skeletal preparation

Skeletal staining was performed as previously described with minor modifications^75^. Postnatal day 0 (P0) forelimbs were fixed in 4% paraformaldehyde in PBS at 4°C, after which excess soft tissue was removed. Samples were dehydrated in 95% ethanol overnight at RT and stained with Alcian blue (Millipore-Sigma) solution for a minimum of two nights at RT. Following staining, forelimbs were washed in 95% ethanol for 5 hours and cleared with 2% KOH for a minimum of three nights at RT. After clearing, additional soft tissue was removed and samples were transferred to Alizarin red (Millipore-Sigma) solution (0.015% in 1% KOH) for a minimum of 2 nights at RT. Additional clearing in 1% KOH was performed as needed. Stained forelimbs were imaged using an Olympus SZX16 UV dissecting microscope equipped with a top-mounted camera.

#### Total RNA sequencing

Sequencing of total RNA from E10.5 and E11.5 WT and mutant forelimbs was previously reported^20^. This study extended the previous analysis by performing total RNA sequencing on E12.5 and E13.5 WT and mutant forelimbs (n=3 biological replicates per timepoint and genotype). Sample quality control, library preparation, and sequencing were performed by the University of Connecticut’s Center for Genome Innovation. RNA quality was assessed using an Agilent 4200 TapeStation and Qubit 3.0. Library preparation was completed using TruSeq Stranded Total RNA Library Sample Prep Kit with RiboZero (Illumina) for depletion of ribosomal RNA. Sequencing was performed on an Illumina NovaSeq 6000, generating approximately 50 million 150 bp paired-end reads per sample.

#### Alignment, quantification, and DE analysis

Read quality was assessed with FastQC and compiled using MultiQC^51^. Reads were aligned to the mm10 reference genome using HISAT2 with the following parameters: --end-to-end --no-discordant --no-mixed --sensitive -p16 ^52^. Resulting SAM files were used as input for IsoEM2 with the following parameter -C 95 to estimate transcript abundances and assign transcripts per million (TPM) values for each gene within a 99% confidence interval based on 200 bootstrapping iterations^53^. The bootstrap.tar.gz file produced from IsoEM2 was subsequently used as input for IsoDE2-based differential expression (DE) analysis. DE analysis was performed using an all-to-all pairwise comparison strategy between all timepoints and genotypes. This included both static comparisons (stage-matched WT and mutant samples) and temporal comparisons (samples from the same genotype across developmental time). TPM values from each biological replicate and log_2_ fold-change values temporal comparisons were used to generate gene expression matrices for WT and mutant datasets.

#### Static DE analysis and PCA

Gene-level TPM values and corresponding Log2 fold changes between WT and mutant samples generated by the HISAT2-IsoEM2-IsoDE2 pipeline were merged across all pairwise comparisons to generate a unified expression matrix. Genes were filtered to retain only protein-coding, autosomal genes based on Ensembl annotation (release 99). To be considered expressed, a gene had to have a TPM of ≥ 0.5 in at least one sample. This threshold was empirically determined to retain biologically relevant lowly expressed genes while minimizing noise, as confirmed by expression of key limb patterning genes such as *Shh* at early developmental stages. Static differential expression was assessed independently at each developmental timepoint by comparing WT and mutant forelimbs. To be considered differentially expressed a gene needed to meet the following criteria: (i) the confident Log2 fold change produced by IsoDE2 based bootstrapping was ≥ |1| and (ii) the gene was considered expressed. All other expressed genes with a log2 fold change < |1| were considered non-differentially expressed (NonDE). All differentially expressed genes were used as input for WATER trajectory analysis.

PCA was performed on gene-level TPM values generated by the HISAT2-IsoEM2-IsoDE2 pipeline. To account for differences in sequencing depth and distribution across samples, quantile normalization was performed separately on WT and mutant datasets, as previously described. The normalized datasets were then merged to generate the final matrix used for PCA. For dimensionality reduction, genes were ranked by variance across samples and PCA was performed using the top 10,000, 5,000, or 2,500 most variable genes. PCA plots presented in figures were generated using the top 5,000 most variable genes, which provided an optimal balance between signal resolution and noise reduction. Expression matrices were transposed to sample-by-gene format and analyzed using the *prcomp* function in R with scaling enabled. Multiple PCA configurations were performed to assess global transcriptomic relationships under different biological contexts, including: (i) all WT and mutant samples, (ii) WT-only samples, (iii) mutant-only samples, (iv) individual WT timepoints analyzed alongside all mutant samples, and (v) individual mutant timepoints analyzed alongside all WT samples. Principal components 1 (PC1) and 2 (PC2) were visualized to evaluate clustering by developmental stage and genotype.

#### WATER trajectory analysis

All R package dependencies required to run WATER can be found in Data S18.

##### Preparing data for WATER

Gene expression and differential expression data were preprocessed and formatted for WATER trajectory analysis using a series of custom R scripts representing the first three steps of the WATER pipeline. Replicate-level gene expression values obtained from IsoEM2 were first merged into a single matrix by extracting gene-level TPM values for each sample and merged across timepoints and replicates based on gene symbols. In parallel, differential expression results from IsoDE2 were used to extract gene-level average TPM values and log2 fold-changes for each comparison and reformatted into a single table. All expression data, including replicate and average TPM values, were then merged with differential expression results to create a unified matrix for subsequent analysis. The resulting dataset contains replicate TPM values, averaged TPM values, and log2 fold-change values across all pairwise timepoint comparisons for each gene. Data from different conditions (WT and mutant) were processed independently. Genes were then filtered to retain those with detectable expression (TPM ≥ 0.5 in at least one sample). Differentially expressed genes were defined based on an absolute log2 fold change threshold (|log2FC| ≥ 1) across any comparison, and genes were classified as differentially expressed or non-differentially expressed. Finally, expression values were quantile normalized across samples to ensure comparability, and genes with zero variance were removed. The resulting quantile-normalized matrix was exported and used as input for WATER trajectory analysis.

##### Initial trajectory analysis

Quantile-normalized expression data were subset to include only genes classified as differentially expressed in the preprocessing step. Expression values were scaled on a per-gene basis and filtered to remove genes with non-finite values or zero variance across samples. To assess global temporal expression structure, hierarchical clustering was performed using Ward’s minimum variance method on Euclidean distances computed from scaled expression values. Heatmaps were generated to visualize gene expression patterns across samples without clustering of columns to preserve temporal ordering. To reduce replicate-level noise, averaged expression values were used as input for downstream clustering analysis. The optimal number of clusters was estimated using silhouette analysis across a range of cluster numbers (k), with the maximum k determined dynamically based on the number of timepoints. Average silhouette widths were computed for each k, and candidate cluster numbers were selected based on high silhouette scores. The final number of clusters carried forward was determined based on both silhouette results and empirical inspection of cluster structure. Cluster assignments were extracted from the hierarchical clustering tree and used to define initial gene expression trajectories. Scaled expression values were plotted across timepoints for each cluster, and average cluster trajectories were computed to represent the mean temporal expression pattern of genes within each cluster. Clusters exhibiting similar trajectory shapes (e.g., maximal expression at the same timepoint) were collapsed into a single group. These initial trajectories served as the basis for subsequent correlation-based refinement and trajectory classification.

##### Initial correlation-based trajectory refinement

Following initial trajectory assignment, correlation-based refinement was performed to evaluate the consistency of individual gene expression profiles with their assigned trajectories. For each trajectory, a representative template was generated by calculating the mean scaled expression across all genes assigned to that trajectory at each timepoint. Each gene’s expression profile was then correlated against all trajectory templates using Pearson correlation. For every gene, the correlation to its assigned trajectory (r_original) and the maximum correlation to any trajectory (r_new) were computed, along with the identity of the best-matching trajectory. Genes were classified based on these correlation values using a decision framework. Genes with strong correlation to their assigned trajectory (r_ original ≥ 0.8 and r_new < 0.8) and no higher correlation to another trajectory retained their original assignment. Genes with weak correlations to all trajectories (r_original < 0.8 and r_new < 0.8) were classified as weak. Genes with a strong correlation to multiple trajectories (r_original ≥ 0.8 and r_new ≥ 0.8) were classified as ambiguous. Genes with a weak correlation to their assigned trajectory but a strong correlation to an alternative trajectory (r_original < 0.8 and r_new ≥ 0.8) were reassigned to the best-matching trajectory. Summary statistics were generated to quantify the proportion of genes retained, reassigned, or flagged for re-clustering. Confusion matrices were constructed to assess transitions between original and updated trajectory assignments. Genes classified as weak or ambiguous were extracted and carried forward for subsequent re-clustering steps, while retained and reassigned genes were used to refine trajectory definitions.

##### Iterative trajectory refinement

Genes classified as weak or ambiguous during the initial correlation analysis were subjected to additional refinement to improve trajectory assignment. For each category, genes were subset from the quantile-normalized expression matrix and analyzed independently. Hierarchical clustering was performed as previously described and the number of clusters carried forward per category was determined based on both silhouette results and empirical inspection of cluster structure. Following re-clustering, trajectory assignments for ambiguous and weak genes were manually curated based on trajectory categories (e.g., early high, late high) according to the timepoint at which maximal or minimal expression occurred. All manual assignments were applied in a single step to generate an updated trajectory label for each gene. The updated trajectory assignments were integrated with automatically assigned and retained trajectories from earlier steps to produce a unified set of gene trajectory labels. These updated labels were used as input for subsequent correlation-based validation and final trajectory assignment.

##### Final trajectory assignment

To refine and validate trajectory assignments, a secondary correlation analysis was performed using the updated trajectory labels. Mean expression profiles were recomputed for each trajectory by averaging quantile-normalized, per-gene expression values across all genes assigned to a given trajectory. For each gene, Pearson correlation coefficients were calculated between its expression profile and the mean profiles of all trajectories. The correlation to the assigned trajectory (r_updated) and the maximum correlation across all trajectories (r_best) were used to assess assignment confidence. Genes were classified into three categories based on a correlation threshold (r ≥ 0.8). Genes with r_updated ≥ 0.8 and equal to r_best were classified as unchanged, indicating strong agreement with their assigned trajectory. Genes with r_best ≥ 0.8 and greater than r_updated were classified as updated and reassigned to the trajectory with the highest correlation. Genes for which no trajectory achieved r ≥ 0.8 were classified as low confidence. Final trajectory labels were generated by integrating these results, such that genes classified as updated were reassigned to their best-correlated trajectory, while all other genes retained their current assignments. Genes classified as low confidence were flagged but retained in the dataset for downstream analyses. The resulting finalized trajectory assignments were used for all subsequent analyses, including stability assessment and downstream biological interpretation.

##### Leave-One-Out (LOO) stability analysis

To assess the robustness of trajectory assignments and clustering structure, a leave-one-out (LOO) stability analysis was performed. For each iteration, a single replicate was removed from the dataset, and gene expression values were recomputed by averaging the remaining replicates at each timepoint. Genes were then re-clustered using the same hierarchical clustering approach applied to the full dataset. The number of clusters (k) was fixed to the value selected during initial clustering. To enable direct comparison between LOO-derived clusters and the original clustering, cluster labels were aligned using a centroid-based matching approach. Specifically, mean expression profiles (centroids) were computed for each cluster in both the full and LOO datasets, and clusters were matched by maximizing Pearson correlation between centroids using the Hungarian algorithm. Clustering concordance between the original and LOO results was quantified using the adjusted Rand index (ARI), calculated before and after cluster label assignment. In addition, cluster retention was defined as the proportion of genes assigned to the same cluster following LOO analysis while trajectory retention was defined based on agreement in simplified trajectory classifications. Retention rates were computed to assess the stability of individual patterns and summary statistics across all LOO iterations were used to evaluate overall robustness. This LOO analysis was applied both after initial clustering and following final trajectory refinement, enabling assessment of stability at multiple stages of the WATER pipeline.

##### Comparative analysis of WT and MUT data

To compare temporal gene expression dynamics between WT and mutant conditions, quantile normalized data were processed to generate matched trajectory profiles across developmental timepoints. Trajectory assignments for WT and mutant data were independently produced by WATER and integrated at the gene level, allowing genes to be grouped based on their transition between WT and mutant states. All possible WT to mutant trajectory combinations were interrogated and genes were subset accordingly. For each WT to mutant trajectory combination, gene-level expression profiles were visualized across timepoints. Individual gene trajectories were plotted with low opacity to represent within-group variability, and mean expression trajectories were overlaid to capture the overall temporal pattern for each condition and cluster. WT and mutant trajectories were displayed simultaneously to facilitate direct comparison of trajectory shifts. Functional enrichment analysis was performed independently for each WT to mutant gene set as described in the *Gene ontology enrichment analysis* section of the methods. In addition to the full gene set, a subset analysis was performed on a predefined list of genes of interest (MIGs). The same trajectory comparison and enrichment workflow was applied to this subset to identify condition-specific functional changes within this targeted gene group. All enrichment results were compiled across trajectory combinations to generate a comprehensive dataset linking trajectory transitions with functional annotations. This analysis enabled identification of trajectory-specific biological processes and provided insight into how gene expression dynamics are altered between WT and MUT conditions.

#### Alternative splicing analysis

Analysis of minor intron retention was performed as previously described (Olthof et al. 2019) with some modifications. Briefly, uniquely mapped reads (NH:i:1) were subset using awk and converted to BAM files using SAMtools. These reads were further subset into spliced and unspliced reads using awk-based partitioning of CIGAR strings. Minor intron coordinates were obtained from previously published annotations. Additional genomic features, including 5’ splice sites, 3’ splice sites, and flanking exons, were derived from these intron coordinates using custom scripts. For intron retention analysis, unspliced reads mapping to minor introns were identified using BEDtools and further filtered to isolate reads spanning exon-intron junctions. A splicing index (SI_IR_) was calculated for each intron as the proportion of unspliced reads relative to the total number of reads (spliced and unspliced) overlapping that intron. For alternative splicing analysis, spliced reads were binned into nine categories (CAT) based on splice junction usage as determined by BEDtools. CAT1 represents canonical splicing of the minor intron, while CAT2-4 represent exon skipping, CAT5-8 represent alternative splice site usage, and CAT9 represents cryptic exon usage. For each alternative splicing category, a splicing index (SI_AS_) was calculated as the proportion of reads assigned to that category relative to the total number of reads supporting either canonical splicing or alternative splicing for the given intron. All intron retention and alternative splicing data were then filtered to produce high-confidence events based on read support and replicate reproducibility (see Quantification and Statistical Analysis for thresholds). Custom scripts for intron retention analysis are publicly available at https://github.com/amolthof/minor-intron-retention.git ^32,33^

#### ORF prediction and AlphaFold modeling

Alternative splicing events were visualized using sashimi plots generated in Integrative Genomics Viewer (IGV) from E11.5 WT and mutant limb samples ^76^. Representative tracks from individual biological replicates were used for visualization. For select genes exhibiting validated alternative splicing events, exon structures corresponding to both canonical and alternative splice isoforms were manually reconstructed using SnapGene^77^. Nucleotide sequences for each isoform were generated by concatenating exons sequences based on observed splice junction usage. Protein domain architecture was annotated using domain information from UniProt^78^. Predicted open reading frames (ORFs) for each reconstructed transcript were identified using the NCBI ORFfinder tool, and the longest ORF was selected for downstream analysis. Protein structures were predicted from the resulting amino acid sequences AlphaFold^79^. Predicted models were visualized to assess the structural consequences of alternative splicing events, including potential disruption of annotated protein domains and overall folding. All SnapGene files, nucleotide sequences, ORF predictions, AlphaFold output files, and videos of protein structures have been deposited in the Mendeley Data repository associated with this study.

#### CUT&RUN profiling and data processing

Cleavage Under Targets and Release Using Nuclease (CUT&RUN) profiling was performed on forelimb buds from E10.5-E12.5 dissected from WT and mutant mouse embryos (n=3 per timepoint and condition). Forelimb buds were dissociated into single-cell suspensions using TrypLE Express (Gibco) or Collagenase (Sigma-Aldrich). CUT&RUN was performed using the EpiCypher CUTANA kit (version 2) according to the manufacturer’s instructions. Cells were immobilized on Concanavalin A-conjugated beads and incubated with an antibody targeting H3K27Me3, with IgG used as a control. Following antibody binding, pAG-MNase digestion was performed to cleave DNA at antibody-bound regions, and released DNA fragments were collected for library preparation and sequencing. *E. coli* spike-in DNA was added to facilitate normalization during data analysis. DNA fragments were used for library preparation with the NEBNext Ultra II DNA library Prep Kit for Illumina and NEBNext Multiplex Oligos for Illumina (Dual Index Primers Set 1) following the manufacturer’s protocol. Libraries were pooled and sequenced on an Illumina NovaSeq 6000 platform at a depth of ∼100 million 100 bp paired-end reads per sample.

Raw sequencing reads were assessed for quality using FastQC and aligned to the mouse reference genome (mm10) using Bowtie2 with the parameters --end-to-end --very-sensitive --no-mixed --no-discordant -p 4 ^54^. Reads were also aligned in parallel to the *E. coli* reference genome (K-12 substr. MG1655) to enable spike-in normalization. Peaks were called using MACS3 with parameters appropriate for broad histone marks (-g mm -f BAMPE --broad) ^56^. Consensus peak sets were generated by merging peaks across all timepoints for each target. Peaks were annotated to genes based on the nearest transcription start site using *ChIPseeker* (v1.44.0) ^56,57^. To quantify signal across samples, consensus peak sets were converted to SAF format and used as input for featureCounts ^59^. Read counts overlapping each peak were quantified for each sample using featureCounts with paired-end mode and a minimum overlap threshold of 50%. Differential binding analysis was performed using *DESeq2* (v1.48.1) ^60^. For each developmental timepoint, samples were subset and differential enrichment between WT and mutant conditions was assessed with a design formula of ∼condition. Log2 fold changes and p-values were calculated for each peak, and results were ordered based on statistical significance. Peaks were considered differentially bound if they met the following criteria: (i) statistical significance as determined by *DESeq2* (p-value < 0.05), (ii) a mean count of ≥ 1 in at least one timepoint, and (iii) an absolute log2 fold change |log2FC| ≥ 0.5. These thresholds were applied to ensure detected differences reflected biologically meaningful changes in signal.

#### CUT&RUN signal visualization

Genome-wide signal tracks (bigWig files) were generated from aligned reads and normalized for sequencing depth prior to visualization of H3K27Me3 enrichment around transcription start sites (TSSs). A reference set of genomic regions was defined by selecting the top 10,000 TSSs ranked by H3K27Me3 signal in E12.5 WT samples. For each region, signal was quantified within a ± 5 kb window centered on the TSS. Signal matrices were generated from bigWig files using the *EnrichedHeatmap* package in R, with a bin size of 50 bp ^62^. For each timepoint, replicate matrices were computed independently and averaged to obtain WT and mutant signal profiles. Heatmaps were generated without row clustering to preserve the reference ordering of regions. Color scales were defined based on the 99^th^ percentile of signal intensity to ensure comparability across conditions. Enrichment profiles were displayed as aggregated signal across all regions.

#### Single-cell RNA sequencing

Forelimb buds were dissected from WT and mutant mouse embryos at E10.5-E12.5 and dissociated into single-cell suspensions using TrypLE Express (Gibco) or Collagenase (Sigma-Aldrich). For each developmental stage and genotype, left and right forelimb buds from multiple embryos were pooled following dissociation (E10.5: n=2; E11.5: n=6, E12.5: n=3 embryos per condition). Single-cell RNA-seq libraries were generated using the 10x Genomics Chromium platform (Single Cell 3’ v3.1 chemistry) according to the manufacturer’s instructions. Cells were diluted to the appropriate concentration to ensure optimal loading. Gel bead-in-emulsion (GEM) reverse transcription, cDNA amplification, and library construction were performed following the manufacturer’s protocol. Final libraries were assessed for fragment size and adapter contamination using the Agilent TapeStation 4200 D1000 High Sensitivity assay and quantified using the dsDNA High Sensitivity Assay on a Qubit 3.0 fluorometer. For each genotype and developmental stage, pooled samples were split across multiple libraries to increase cell capture efficiency. These libraries represent independent technical captures from the same biological pool rather than independent biological replicates. Sequencing was performed on an Illumina NovaSeq 6000 using version 1.5 chemistry to generate approximately 50,000 reads per cell. Raw sequencing reads were processed using the 10x Genomics Cell Ranger pipeline (v7.0.1) for demultiplexing, alignment, and generation of gene-barcode count matrices. BCL files were converted to fastq format using cellranger mkfastq. Reads were aligned to the mouse reference genome (mm10) using cellranger count with the parameter --nosecondary. Cell Ranger aggregation was used to combine libraries within each genotype and developmental stage. Aggregation was performed with normalization to mapped reads to account for differences in sequencing depth across libraries. Processed data were output as gene-barcode expression matrices which were then imported into R (v4.5.1) and analyzed using *Seurat* (v5.4.0) ^63^.

#### scRNA-seq data filtering and processing

Datasets corresponding to each developmental stage (E10.5, E11.5, and E12.5) and genotype (WT and mutant) were initially processed independently. Seurat objects were created from filtered feature-barcode matrices using a minimum threshold of 200 detected genes per cell and requiring genes to be expressed in at least 5 cells. Quality control filtering was performed by excluding cells with fewer than 200 or more than 6,000 detected genes and cells with greater than 10% mitochondrial transcript content. Following quality control, data were normalized using log normalization with a scale factor of 10,000. Highly variable genes were identified using the variance stabilizing transformation (VST) method, selecting the top 1,000 variable features per dataset. Data were then scaled across all genes prior to performing dimensional reduction. Cell cycle effects were regressed during initial data scaling prior to dimensional reduction. No additional regression was performed after dataset integration to avoid overcorrection and preserve biologically meaningful differences between conditions. PCA was performed using the identified variable features, and the top principal components were used to construct a shared nearest neighbor (SNN) graph. Cells were clustered using a graph-based approach with a resolution parameter of 0.5 and visualized using Uniform Manifold Approximation and Projection (UMAP).

#### scRNA-seq data integration and analysis

To enable comparison between WT and mutant datasets within each developmental stage, datasets were integrated using Seurat’s anchor-based integration workflow. Briefly, datasets were independently normalized, and variable features were identified prior to selecting integration features. Integration anchors were identified using FindIntegrationAnchors, and datasets were combined using IntegrateData. The integrated data were scaled and subjected to PCA, UMAP embedding, and clustering as previously described.

To enable joint analysis across developmental stages and genotypes, processed Seurat data from all conditions (E10.5, E11.5, and E12.5; WT and U11 mutant) were integrated as previously described except that reciprocal PCA (RPCA) reduction was used for integration. RPCA-based integration was selected to minimize overcorrection and preserve biologically meaningful differences between genotypes. The resulting integrated dataset was subjected to PCA and UMAP visualization using the top principal components prior to downstream analysis. Integrated clustering and visualization enabled comparison of cell populations across developmental time and genotype within a shared low-dimensional space. Cluster-specific marker genes were identified using the FindAllMarkers function with parameters min.pct = 0.25 and logfc.threshold = 0.25, and top 20 markers were used in identifying cell types along with canonical marker gene expression from previous studies. Gene expression was visualized using violin plots and feature plots. Cell counts per cluster were quantified for each developmental stage and genotype and used to calculate the relative abundance of each cell population. For each cluster, proportions were determined by dividing the number of cells assigned to that cluster by the total number of cells captured within the corresponding genotype and timepoint. To assess differences in cell population representation between genotypes, the ratio of the mutant to WT proportions was calculated for each cluster and timepoint. Values less than 1 indicate relative enrichment in WT samples, values equal to 1 indicate equal representation, and values greater than 1 indicate relative enrichment in mutant samples.

#### Pseudotime and Monocle3 analysis

Single-cell RNA-seq data were analyzed using *Monocle3* to infer developmental trajectories and pseudotime progression ^64^. Cells were first subset by genotype (WT and U11 mutant) prior to trajectory inference to avoid introducing genotype-driven structure into the learned manifold. For each genotype, a *Monocle3* cell data set (cds) object was constructed and processed following the standard *Monocle3* workflow, including preprocessing, dimensional reduction, graph learning, and cell ordering. Pseudotime values were calculated for each cell using the pseudotime function and exported for downstream analysis. Because pseudotime was computed independently for each genotype, values were not directly comparable across datasets. To enable cross-genotype visualization, a global scaling approach was applied. Pseudotime values from WT and mutant limbs were combined, and a shared minimum and maximum value were used to rescale pseudotime to a 0-1 range according to the following formula:

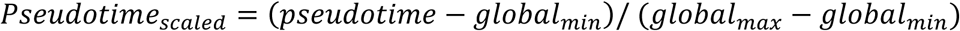

Scaled pseudotime values were added to the *Monocle3* object metadata and used for visualization of developmental progression between WT and mutant datasets.

### Quantification and Statistical Analysis

#### Filtering alternative splicing data

To identify high-confidence alternative splicing events, SI_AS_, TPM, and raw read counts derived from gene expression and alternative splicing analysis were used. For each intron, values were averaged across biological replicates before filtering. An AS event was kept if it satisfied all the following criteria: (i) at least one alternatively spliced read per three million uniquely mapped reads for the category of interest, (ii) SI_AS_ ≥ 0.10, and (iii) the parent gene was classified as expressed based on IsoEM2-derived TPM values. Introns with SI_AS_ values of 0 or 1 in both conditions were excluded, as these represent exclusively canonical or exclusively alternative splicing events and therefore do not permit quantitative comparisons of splicing between conditions. To identify high-confidence intron retention events, SI_IR_, TPM, read coverage, and raw read counts were considered. An intron was considered retained if at least one biological replicate satisfied all the following criteria: (i) ≥ 95% read coverage across the intron body, (ii) >4 reads spanning exon-intron junctions (5’ and/or 3’), (iii) ≥ 1 read spanning the 5’ exon-intron boundary, and (iv) ≥ 1 read spanning the 3’. To evaluate systematic differences in SI_IR_ across timepoints, median SI_IR_ values were compared across stages using multiple baseline references (E10.5, E11.5, E12.5, and E13.5). This analysis revealed a global increase in SI_IR_ at E12.5 relative to other timepoints, independent of baseline selection, indicative of a timepoint-specific global shift. To enable direct comparison across timepoints, SI_IR_ values at E12.5 were centered by subtracting the median shift (0.17) from both WT and mutant samples. This normalization preserves relative differences between genotypes while correcting for timepoint-specific global offsets in intron retention. To assess global shifts in intron retention across shared introns between genotypes per timepoint, paired, two-sided Wilcoxon signed-rank tests were performed on SI_IR_ values. Statistical significance was defined as p < 0.05.

#### Gene ontology enrichment analysis

Functional enrichment analysis was performed using Gene Ontology (GO) annotations from the R package *clusterProfiler* ^66–68^. Gene symbols were converted to Entrez Gene IDs using the bitr function with the *org.Mm.eg.db* annotation database, and gene sets that failed to map were excluded from downstream analysis. GO enrichment was performed separately for the Biological Process (BP), Molecular Function (MF), and Cellular Component (CC) ontologies using the enrichGO function. Adjusted p-values were calculated using the Benjamini-Hochberg method, and terms passing default clusterProfiler thresholds (p < 0.05, q < 0.2) were retained. Redundant GO terms were reduced using the simplify function (cutoff = 0.7) to collapse semantically similar categories and retain only the most significant term based on adjusted p-values. For all analyses, the background gene set consisted of all mouse genes annotated in *org.Mm.eg.db* with valid Entrez Gene IDs. Enrichment results were exported separately for each ontology and combined into a single output for downstream interpretation and visualization.

#### Densitometric analysis of RT-PCR

Densitometric quantification of RT-PCR bands was performed using ImageJ. For each lane, a fixed-size rectangular region of interest (ROI) was drawn around bands corresponding to the target transcript and *Gapdh*. Background signal was measured using an identically sized ROI placed in a region of the same lane lacking visible bands. Integrated density values were obtained for each band, and background-subtracted signal intensities were calculated by subtracting the background measurement from each band. Relative expression was quantified as a ratio of background-subtracted band intensity to the corresponding *Gapdh* band intensity within the same lane. Mean values were plotted for visualization with individual biological replicates overlaid. Raw RT-PCR images used in quantification have been deposited in the Mendeley Data repository associated with this study.

#### 3D reconstruction from serial sections

Three-dimensional (3D) reconstructions were generated from serial histological sections using Reconstruct software ^80^. Serial images were imported into separate series for WT and mutant samples. Images were calibrated using a 50 μm scale bar and manually aligned across sections to ensure spatial continuity. Following alignment, Sox9 signal and the section boundaries were manually traced for each section. Traces were then used to create 3D objects using the Boissonnat surface reconstruction algorithm implemented in Reconstruct with default parameters. This process was performed for three WT and three mutant limb buds.

#### Section staining quantification

Histological section images were imported into QuPath (v0.6.0) for quantitative analysis ^81^. Images from WT and mutant samples were analyzed together and image type was set to fluorescence. Section size calibration was performed using the scale bar present in each image, assuming equal pixel dimensions in the x and y axes. For each section, the limb outline and Sox9-positive regions were manually annotated by tracing regions of interest (ROIs). Limb area and Sox9-positive area were measured in square microns (μm^2^) based on these annotations. Linear measurements, including total limb length, Sox9 domain length, and the distance between the distal edge of the Sox9 domain and the distal limb boundary (referred to as the “gap distance”), were measured in microns (μm). All area and length measurements were converted to mm^2^ and mm, respectively. Quantified values were used to calculate normalized metrics, including the ratio of Sox9 area to total limb area and gap distance normalized to Sox9 length or total limb length. Differences between genotypes were assessed at each developmental stage for each measurement using two-sided Welch’s t-tests to account for potential unequal variance between groups.

Quantification of TUNEL-positive puncta was performed in QuPath using a workflow similar to that described for Sox9 staining. TUNEL-positive puncta were detected using QuPath’s automatic cell detection function, with parameters for pixel size, intensity thresholds, and channel detection held constant across WT and mutant samples. Limb area was manually annotated and measured in square microns (μm^2^) and converted to mm^2^ as described above. For each section, the number of TUNEL-positive puncta was normalized to total limb area to generate a density metric (puncta per mm^2^). Quantification was performed across n=4 biological replicates per genotype (WT, U11 mutant, and U11-p53 mutant) at E12.5. Differences between genotypes were assessed using a one-way analysis of variance (ANOVA) followed by post hoc multiple comparison testing with Tukey’s test. Adjusted p-values < 0.05 were considered statistically significant. Raw section images used in the quantification of section staining have been deposited in the Mendeley Data repository associated with this study.

#### Quantification of limb surface area

White light images of WT and mutant limbs at E10.5-E13.5 were acquired using an Olympus SZX16 UV dissecting microscope equipped with a top-mounted camera. Images were used to quantify changes in limb surface area across developmental time. Images were imported into ImageJ and calibrated using a 1 mm scale bar. Limb surface area was measured in mm^2^ by manually tracing the limb perimeter to create an ROI. A minimum of 3 biological replicates were used per genotype and timepoint. Differences in limb surface area across developmental timepoints and between genotypes were assessed using a two-way ANOVA with timepoint and genotype as factors, including an interaction term. Post hoc multiple comparisons were performed using Tukey’s honestly significant difference (HSD) test. Raw images used in quantification have been deposited in the Mendeley Data repository associated with this study.

#### Quantification of skeletal elements

Skeletal preparations from WT, U11 mutant, and U11p53 mutant P0 forelimbs were dissected to isolate individual long bones and imaged using an Olympus SZX16 UV dissecting microscope equipped with a top-mounted camera. Images were imported into ImageJ and spatial calibration was performed using a 1 mm scale bar. Long bone length was measured by drawing a line segment from one epiphysis to the other. Ossified length was measured as the length of the Alizarin red stained region, and bone width was measured at the midpoint of each long bone (Figure S29D). Measurements were collected for stylopod (humerus) and zeugopod (radius/ulna) elements from both left and right forelimbs. Left and right measurements were averaged for each biological replicate prior to downstream analysis. A minimum of n = 3 biological replicates per genotype were included in the analysis. Differences in long bone length, width, and ossified length between genotypes and skeletal elements were assessed using a two-way ANOVA with genotype and skeletal element as factors, including an interaction term. Post hoc multiple comparisons were performed using Tukey’s HSD test. Raw images used in quantification have been deposited in the Mendeley Data repository associated with this study.

