## Supplemental Information for "WATER reveals heterochrony of molecular programs underlies developmental failure caused by minor spliceosome inhibition"

**Contents**

**Supplementary Figures**

Figure S1. Functional enrichment from stage-matched differential expression analysis and reciprocal PCA projections, related to Figure 1.

Figure S2. Application of WATER to temporal RNA-seq data from WT mouse forelimbs, related to Figure 1.

Figure S3. Initial differential expression analysis and hierarchical clustering in WT mouse forelimbs, related to Figure 1.

Figure S4. Initial temporal differential expression and hierarchical clustering analysis of mutant forelimbs, related to Figure 1.

Figure S5. Reproducibility assessment of initial WT and mutant gene expression trajectories, related to Figure 1.

Figure S6. Correlation-based reassessment of WT trajectory assignments identifies weak and ambiguous gene expression profiles, related to Figure 1.

Figure S7. Correlation-based reassessment of mutant trajectory assignments identifies weak and ambiguous expression profiles, related to Figure 1.

Figure S8. Final correlation-based refinement confirms high-confidence gene expression trajectory assignments in WT and mutant forelimbs, related to Figure 1.

Figure S9. Validation of final expression trajectory assignments in WT and mutant forelimbs, related to Figure 1.

Figure S10. Benchmarking WATER against TC-seq for high confidence temporal trajectory inference, related to Figure 1.

Figure S11. Functional enrichment of expression trajectories reveals differences between WT and mutant forelimbs, related to Figure 1.

Figure S12. Redistribution of High WT-only expression trajectories following U11 loss, related to Figure 2.

Figure S13. Redistribution of Low WT-only expression trajectories following U11 loss, related to Figure 2.

Figure S14. Redistribution of Early High, Late High, and biphasic WT-only expression trajectories following U11 loss, related to Figure 2.

Figure S15. Redistribution of E10.5 and E13.5 High WT-only trajectories following U11 loss, related to Figure 2.

Figure S16. Superimposing static differential gene expression and expression trajectories reveals widespread changes in magnitude independent of trajectory shifts, related to Figure 2.

Figure S17. Validation of trajectory shifts by whole-mount in situ hybridization and RT-PCR analysis, related to Figure 2.

Figure S18. MIG splicing is dynamically regulated across developmental time in the WT forelimb, related to Figure 3.

Figure S19. Altered MIG splicing upon U11 loss does not globally correlate with shifts in MIG expression, related to Figure 3.

Figure S20. Predicted consequences of minor intron mis-splicing in DNA repair and chromatin-associated MIGs, related to Figure 3.

Figure S21. Minor intron mis-splicing converges on critical MIG nodes within diverse regulatory networks, including epigenetic modifiers such as Eed, related to Figure 3.

Figure S22. Extended analysis of H3K27Me3 dynamics across developmental time in U11 mutant forelimbs, related to Figure 4.

Figure S23. IgG control profiles for H3K27Me3 CUT&RUN experiments, related to Figure 4.

Figure S24. Replicate concordance and H3K27Me3 signal profiles across developmental timepoints, related to Figure 4.

Figure S25. Single-cell cluster annotation across developmental timepoints, related to Figure 6.

Figure S26. Spatial marker-based organization of single-cell clusters, related to Figure 6.

Figure S27. Histological validation of chondrogenesis defects in mutant forelimbs, related to Figure 6.

Figure S28. Dissecting the relationship between gene expression trajectories and shifts in cellular abundance in WT and mutant forelimbs, related to Figure 6.

Figure S29. Phenotypic analysis of U11-p53 mutant limbs, related to Figure 7.

**Supplementary Tables**

Table S1. Summary statistics from limb surface area quantifications, related to Figure 1.

Table S2. Comparison of WATER with other analysis methods for temporal RNA-seq data, related to Figure S10.

Table S3. Redistribution of functional modules across temporal trajectories following minor spliceosome inhibition, related to Figure 2. See also Data S7.

Table S4. Summary statistics from quantification of distal ectoderm-Sox9 gap distance and Sox9-positive area quantifications, related to Figure S27.

Table S5. RT-PCR primer sequences, related to Figure 2, 3, 6, S17, S20, S27 and S29.

**Supplementary Datasets**

Data S1. Genes overrepresented in each genotype and timepoint, related to Figures 1E, S1A, and S23.

Data S2. Functional enrichment of genes overrepresented in each genotype and timepoint, related to Figure S1A.

Data S3. Replicate expression and log2 fold-change data from WT and U11 mutant forelimbs across time, related to Figures S3 and S9.

Data S4. WATER trajectories for WT and mutant datasets, related to Figure 1J.

Data S5. Functional enrichment results of all WT gene expression trajectories, related to Figure S11A.

Data S6. Functional enrichment results of all U11 mutant gene expression trajectories, related to Figure S11A'.

Data S7. Functional enrichment results for all trajectory shift categories between WT and U11 mutant datasets, related to Figures 2B-C and S12-S15.

Data S8. Shifts in minor intron-containing gene trajectories between WT and mutant datasets, related to Figure 3.

Data S9. List of minor intron-containing genes associated with chromatin-modifying activity, related to Figure 3H.

Data S10. Parent genes of peaks with overrepresented H3K27Me3 per genotype and timepoint, related to Figure 4.

Data S11. WATER trajectories of H3K27Me3 in WT and U11 mutant forelimbs, related to Figure 4G.

Data S12. Functional enrichment of H3K27Me3 relationships, related to Figure 4I.

Data S13. WATER trajectories for WT and Eed-KO datasets, related to Figure 5.

Data S14. Functional enrichment results for WT, Eed-KO, and shifts between the two datasets, related to Figure 5.

Data S15. Top cluster marker genes identified from single cell data, related to Figures 6B and S25B.

Data S16. References used for spatial annotation of single-cell RNA-Seq data, Figure S26.

Data S17. WATER trajectories from WT and U11 mutant single-cell RNA-seq data, related to Figure S28.

Data S18. R package dependencies required for WATER analysis.

**Supplementary Figures**

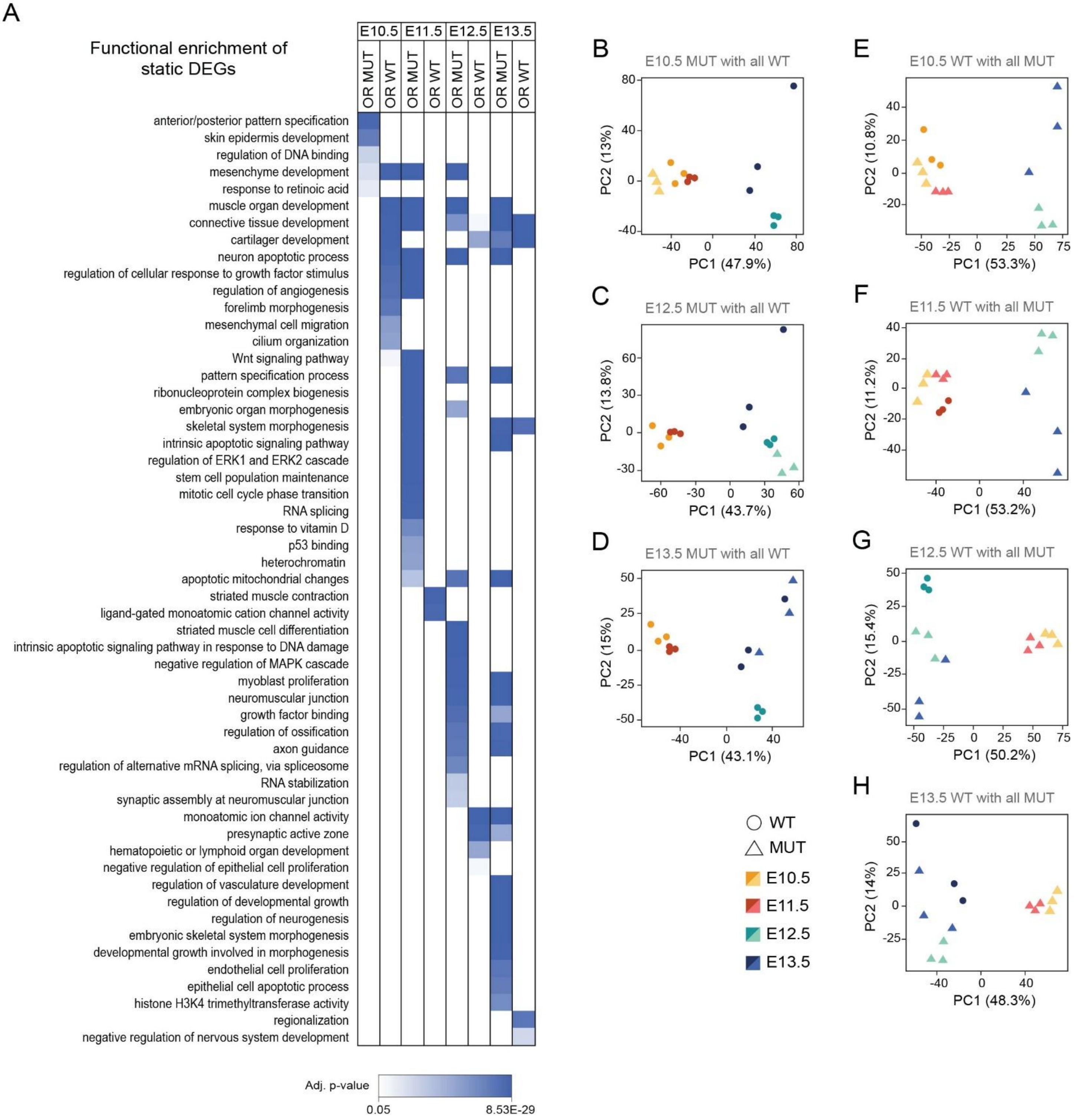


**Figure S1. Functional enrichment from stage-matched differential expression analysis and reciprocal PCA projections, related to Figure 1.** (A) Functional enrichment analysis of static differentially expressed genes across developmental stages. Heatmap shows significantly enriched GO terms (adjusted p-value < 0.05) for overrepresented (OR) WT and OR mutant gene sets at each timepoint. Color intensity reflects magnitude of significance. (B-D) Principal component analysis (PCA) projections of stage specific mutant samples combined with all WT timepoints. In each case, the mutant sample clusters with the stage-matched WT samples. (E-H) PCA projections of stage specific WT samples combined with all mutant timepoints. In each case, the WT sample clusters with the stage-matched mutant samples. See also Figure 1E and 1H.

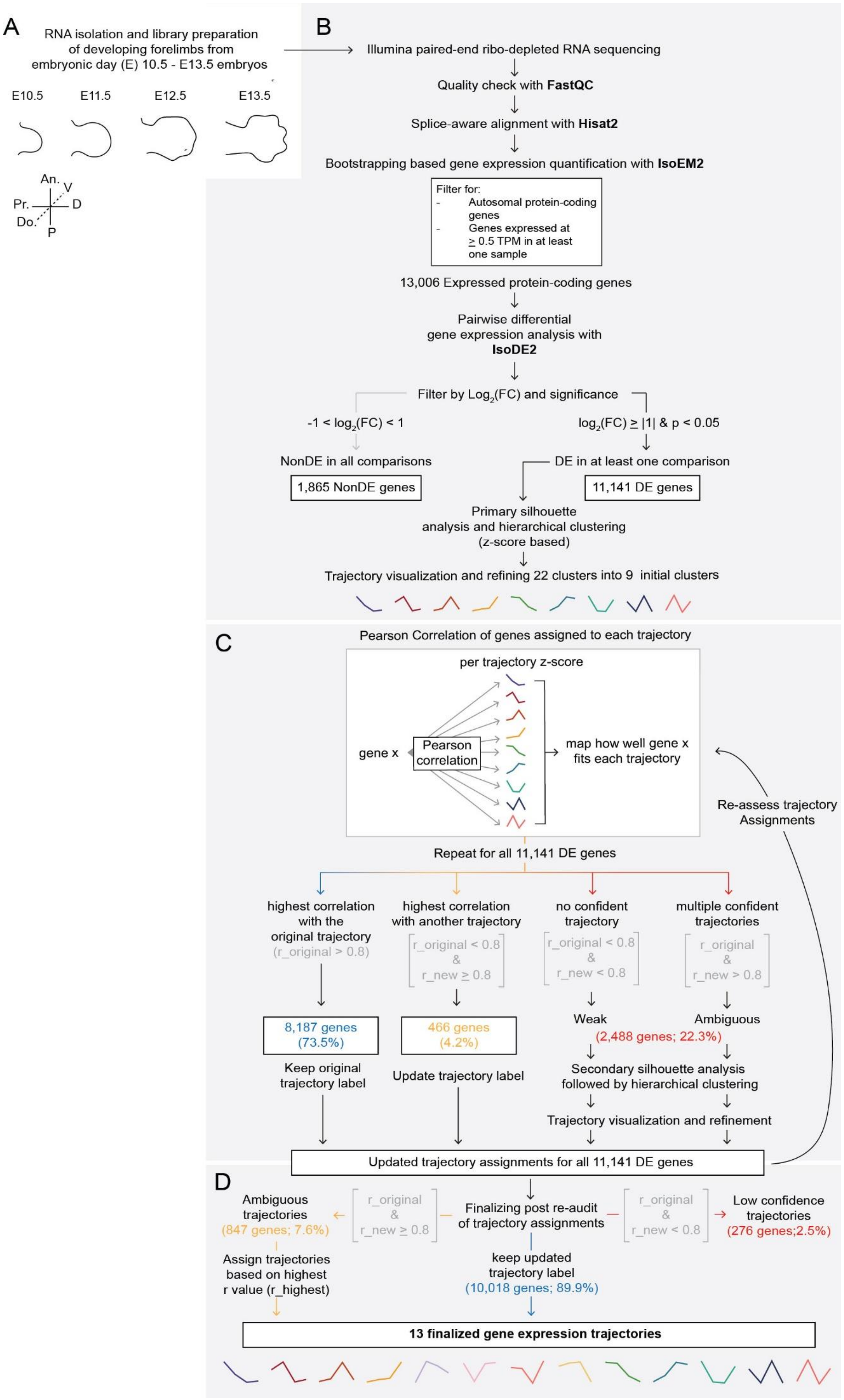
A
RNA isolation and library preparation of developing forelimbs from embryonic day (E) 10.5 - E13.5 embryos
E10.5
E11.5
E12.5
E13.5
An.
V
Pr.
D
Do.
P
B
Illumina paired-end ribo-depleted RNA sequencing
Quality check with FastQC
Splice-aware alignment with Hisat2
Bootstrapping based gene expression quantification with IsoEM2
Filter for:
- Autosomal protein-coding genes
- Genes expressed at ≥ 0.5 TPM in at least one sample
13,006 Expressed protein-coding genes
Pairwise differential gene expression analysis with IsoDE2
Filter by Log2(FC) and significance
-1 < log2(FC) < 1
log2(FC) ≥ |1| & p < 0.05
NonDE in all comparisons
DE in at least one comparison
1,865 NonDE genes
11,141 DE genes
Primary silhouette analysis and hierarchical clustering (z-score based)
Trajectory visualization and refining 22 clusters into 9 initial clusters
C
Pearson Correlation of genes assigned to each trajectory
per trajectory z-score
gene x
Pearson correlation
map how well gene x fits each trajectory
Re-assess trajectory Assignments
Repeat for all 11,141 DE genes
highest correlation with the original trajectory
(r_original > 0.8)
highest correlation with another trajectory
r_original < 0.8 & r_new ≥ 0.8
no confident trajectory
r_original < 0.8 & r_new < 0.8
multiple confident trajectories
r_original & r_new > 0.8
8,187 genes (73.5%)
466 genes (4.2%)
Weak
Ambiguous
(2,488 genes; 22.3%)
Keep original trajectory label
Update trajectory label
Secondary silhouette analysis followed by hierarchical clustering
Trajectory visualization and refinement
Updated trajectory assignments for all 11,141 DE genes
D
Ambiguous trajectories
(847 genes; 7.6%)
r_original & r_new ≥ 0.8
Finalizing post re-audit of trajectory assignments
r_original & r_new < 0.8
Low confidence trajectories
(276 genes;2.5%)
Assign trajectories based on highest r value (r_highest)
keep updated trajectory label
(10,018 genes; 89.9%)
13 finalized gene expression trajectories

**Figure S2. Application of WATER to temporal RNA-seq data from WT mouse forelimbs, related to Figure 1.** (A) Bulk RNA-seq datasets were generated from whole WT mouse forelimbs collected at E10.5-E13.5. Forelimb schematics depict each developmental stage, and a key is provided to denote the developmental axes. (B-E) Schematization of the WATER pipeline used to assess temporal gene expression patterns. (B) Following quality control (FastQC), alignment (HISAT2), and gene expression quantification (IsoEM2); 13,006 autosomally encoded protein-coding genes with an average expression $\geq$ 0.5 TPM in at least one sample were retained for subsequent analysis. Pairwise differential expression analysis (IsoDE2) identified 11,141 differentially expressed (DE) genes and 1,865 non-differentially expressed (NonDE) genes. To characterize temporal expression patterns, z-score-transformed DE genes were subjected to silhouette analysis and hierarchical clustering. Silhouette analysis supported a broad range of robust clustering windows (k = 2-22); selection of k = 22 followed by trajectory consolidation based on shared temporal patterns yielded nine initial expression trajectories ($E10.5^{Hi}$, $E11.5^{Hi}$, $E12.5^{Hi}$, $E13.5^{Hi}$, $Early^{Hi}$, $Late^{Hi}$, $E10.5^{Hi}$ and $E13.5^{Hi}$, Up-Down-Up, and Down-Up-Down). (C) To validate and refine trajectory assignments, each gene's expression profile was compared to the mean expression profile of all trajectories using Pearson correlation. Most genes (8,187; 73.5%) showed the strongest correlation with their originally assigned trajectory (Pearson correlation coefficient (r) for the original trajectory ≥ 0.8; panel C, blue). Genes whose expression patterns more strongly matched a different trajectory (r_original < 0.8 & r_new ≥ 0.8; panel C, yellow) were reassigned, whereas genes with weak (r_original < 0.8 and r_new < 0.8) or ambiguous (r_original ≥ 0.8 and r_new ≥ 0.8) correlations were flagged for further evaluation (panel C, red). Genes with weak or ambiguous assignments were reanalyzed using an additional round of silhouette analysis and hierarchical clustering. This revealed four additional trajectories characterized by low expression at a single timepoint ($E10.5^{Lo}$, $E11.5^{Lo}$, $E12.5^{Lo}$, and $E13.5^{Lo}$). This additional round of clustering produced updated trajectory assignments for all 11,141 DE genes which were subjected to a second correlation-based audit to assess improvement in trajectory fit. (D) After the second audit, 13 high-confidence trajectories were defined (median r = 0.94), encompassing 97.5% (10,865 out of 11,141 genes) of all DE genes. The remaining 276 genes (2.5%) exhibited low similarity to any trajectory and were retained as a low-confidence category for transparency. An.; anterior, P; posterior, D; distal, Pr.; proximal, Do.; dorsal, V; ventral. See also Figures S3-S10.

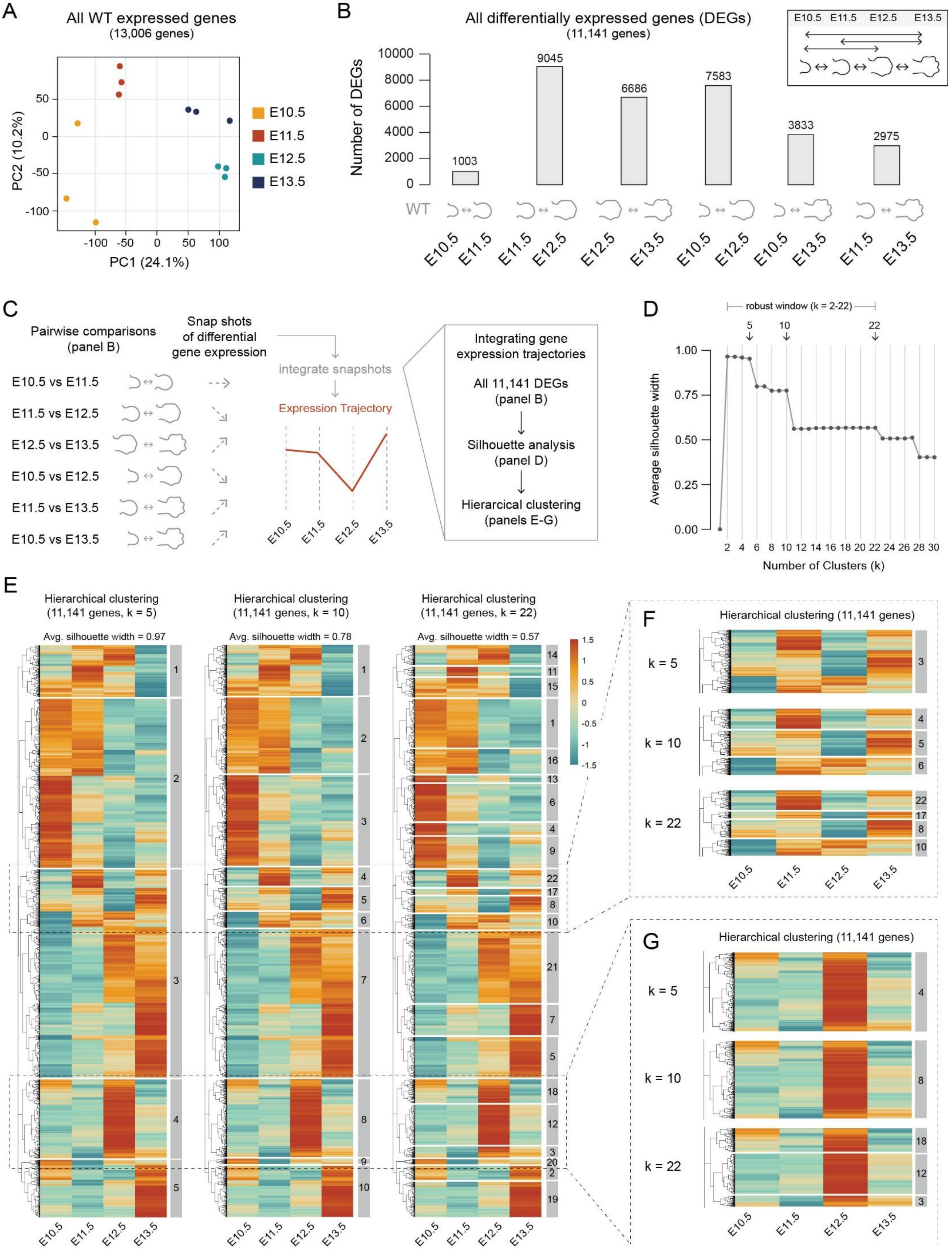

A
All WT expressed genes
(13,006 genes)
PC1 (24.1%)
PC2 (10.2%)
E10.5
E11.5
E12.5
E13.5
B
All differentially expressed genes (DEGs)
(11,141 genes)
Number of DEGs
1003
9045
6686
7583
3833
2975
WT
C
Pairwise comparisons
(panel B)
Snap shots
of differential
gene expression
integrate snapshots
Expression Trajectory
E10.5 vs E11.5
E11.5 vs E12.5
E12.5 vs E13.5
E10.5 vs E12.5
E11.5 vs E13.5
E10.5 vs E13.5
Integrating gene
expression trajectories
All 11,141 DEGs
(panel B)
Silhouette analysis
(panel D)
Hierarcical clustering
(panels E-G)
D
robust window (k = 2-22)
Average silhouette width
Number of Clusters (k)
E
Hierarchical clustering
(11,141 genes, k = 5)
Avg. silhouette width = 0.97
Hierarchical clustering
(11,141 genes, k = 10)
Avg. silhouette width = 0.78
Hierarchical clustering
(11,141 genes, k = 22)
Avg. silhouette width = 0.57
F
Hierarchical clustering (11,141 genes)
k = 5
k = 10
k = 22
G
Hierarchical clustering (11,141 genes)
k = 5
k = 10
k = 22

**Figure S3. Initial differential expression analysis and hierarchical clustering in WT mouse forelimbs, related to Figure 1.** (A) PCA of all genes expressed in WT forelimbs from E10.5 to E13.5, demonstrating temporal separation of samples prior to trajectory analysis with WATER. (B) Number of differentially expressed genes (DEGs) identified in each pairwise temporal comparison, including comparisons between non-adjacent timepoints as indicated by schematic icons below each bar. (C) Conceptual overview illustrating how pairwise differential expression snapshots are integrated across timepoints to reconstruct gene expression trajectories. All genes identified in panel B were retained for subsequent trajectory analysis. (D) Silhouette analysis across increasing numbers of clusters (k) for all DEGs identified a robust clustering window (k = 2-22) where the average silhouette width was greater than 0.5. These values were used to guide subsequent hierarchical clustering. (E) Hierarchical clustering heatmaps of all WT DEGs (z-scored expression) at representative k values (5, 10, and 22). Heatmaps illustrate increasing resolution of temporal expression patterns with higher cluster numbers. Average silhouette width is indicated for each k value. (F-G) Zoomed views of representative clusters from k = 5, 10, and 22, highlighting the emergence and refinement of distinct clusters as resolution increases.

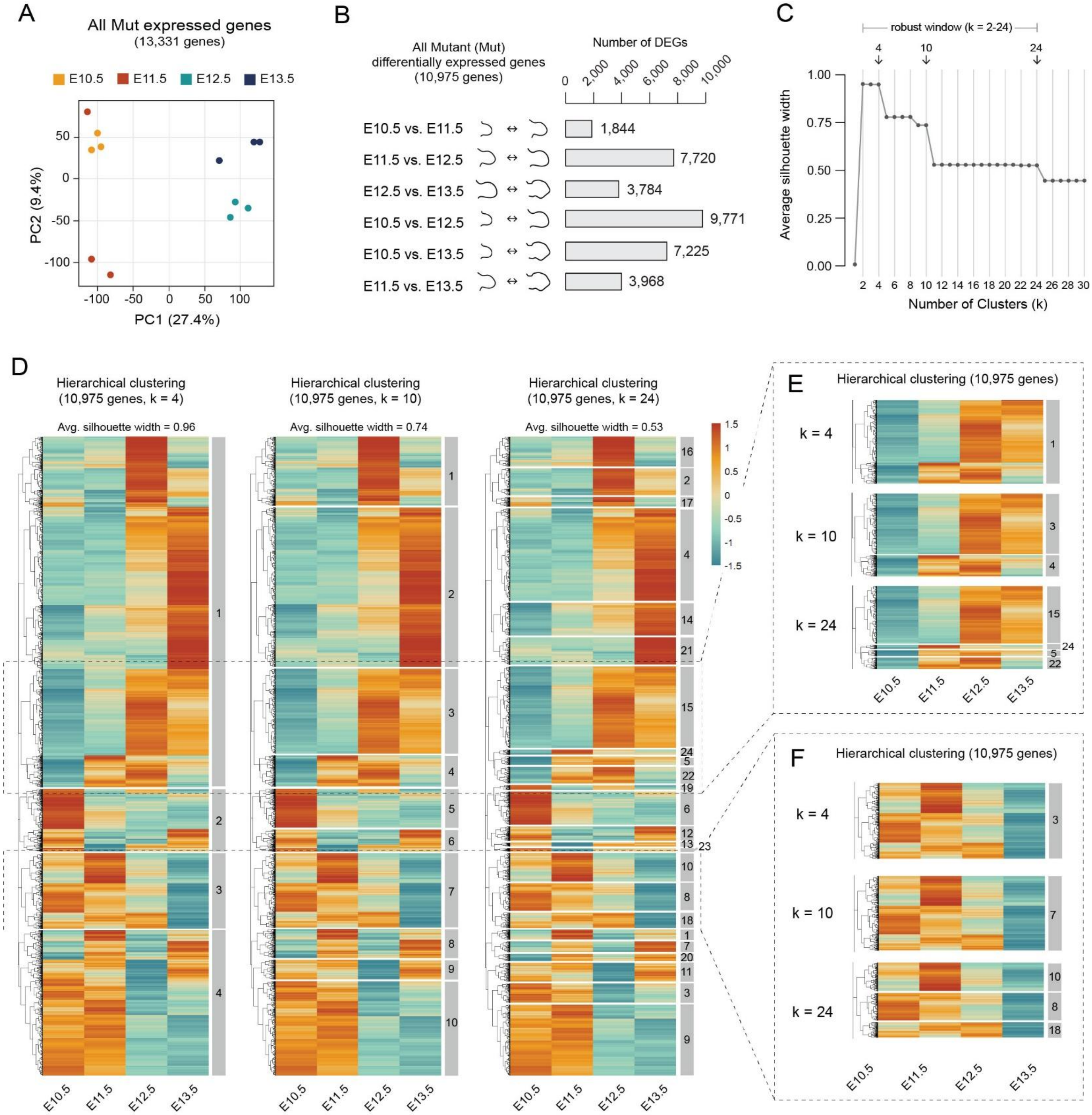


**Figure S4. Initial temporal differential expression and hierarchical clustering analysis of mutant forelimbs, related to Figure 1.** (A) PCA of all genes expressed in mutant forelimbs (13,331 genes), demonstrating temporal separation of samples prior to trajectory analysis. (B) Number of DEGs identified across all pairwise temporal comparisons in mutant forelimbs, including both adjacent and non-adjacent developmental timepoints. All DEGs were carried forward into trajectory analysis. (C) Silhouette analysis across increasing number of clusters (k) for all mutant DEGs identified a robust window of clustering (k = 2-24) where the average silhouette width was greater than 0.5. These values were used to guide subsequent hierarchical clustering. (D) Hierarchical clustering heatmaps of all mutant DEGs (10,975 genes; z-scored expression) at representative k values (4, 10, and 24). Heatmaps illustrate increasing resolution of temporal expression patterns with higher cluster numbers. (E-F) Zoomed views of representative clusters from k = 4, 10, and 24, highlighting the emergence and refinement of distinct clusters as resolution increases.

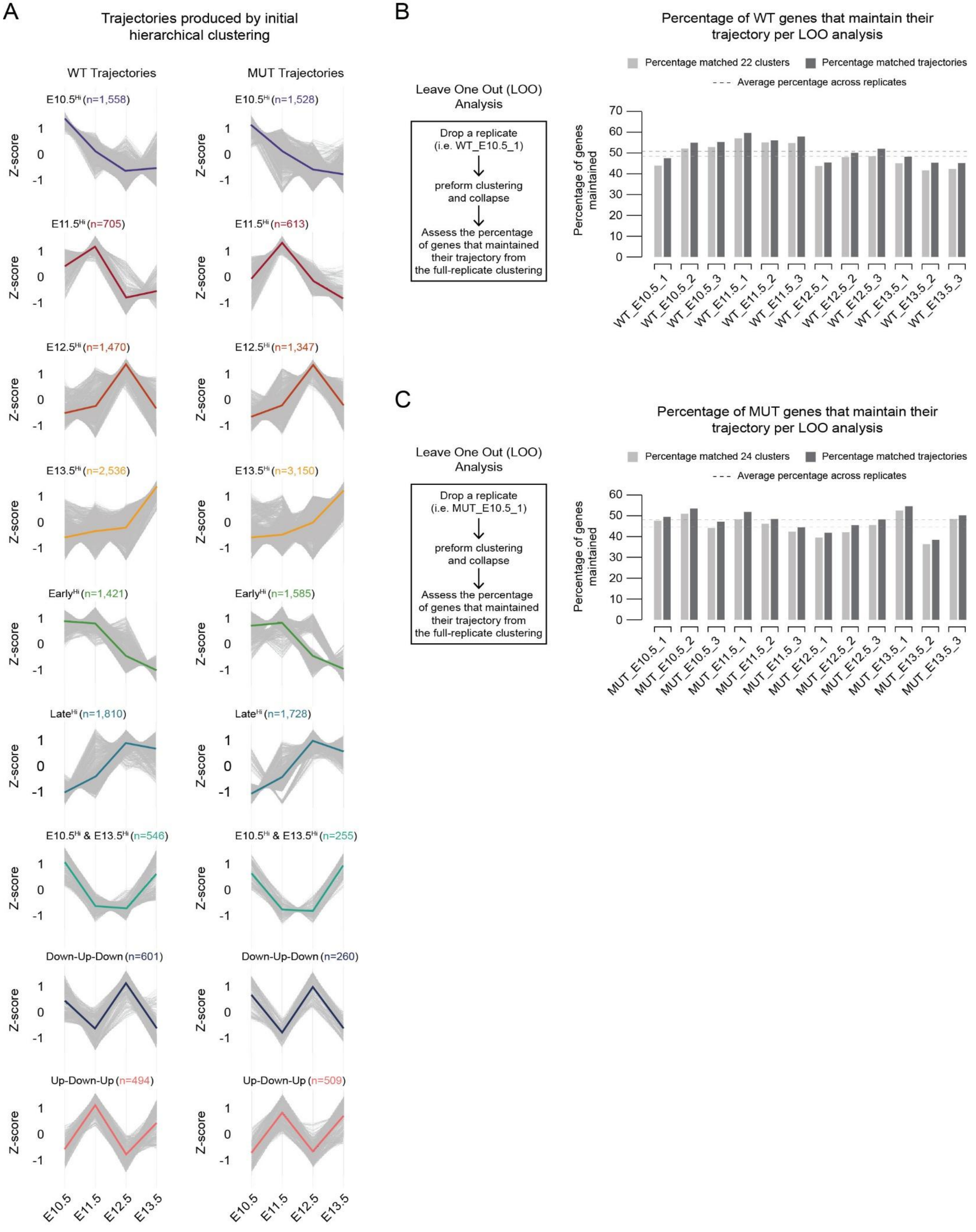
A
Trajectories produced by initial hierarchical clustering
WT Trajectories
MUT Trajectories
E10.5Hi (n=1,558)
E10.5Hi (n=1,528)
E11.5Hi (n=705)
E11.5Hi (n=613)
E12.5Hi (n=1,470)
E12.5Hi (n=1,347)
E13.5Hi (n=2,536)
E13.5Hi (n=3,150)
EarlyHi (n=1,421)
EarlyHi (n=1,585)
LateHi (n=1,810)
LateHi (n=1,728)
E10.5Hi & E13.5Hi (n=546)
E10.5Hi & E13.5Hi (n=255)
Down-Up-Down (n=601)
Down-Up-Down (n=260)
Up-Down-Up (n=494)
Up-Down-Up (n=509)
Z-score
E10.5
E11.5
E12.5
E13.5
B
Percentage of WT genes that maintain their trajectory per LOO analysis
Leave One Out (LOO) Analysis
Drop a replicate (i.e. WT_E10.5_1)
preform clustering and collapse
Assess the percentage of genes that maintained their trajectory from the full-replicate clustering
Percentage matched 22 clusters
Percentage matched trajectories
Average percentage across replicates
Percentage of genes maintained
WT_E10.5_1
WT_E10.5_2
WT_E10.5_3
WT_E11.5_1
WT_E11.5_2
WT_E11.5_3
WT_E12.5_1
WT_E12.5_2
WT_E12.5_3
WT_E13.5_1
WT_E13.5_2
WT_E13.5_3
C
Percentage of MUT genes that maintain their trajectory per LOO analysis
Leave One Out (LOO) Analysis
Drop a replicate (i.e. MUT_E10.5_1)
preform clustering and collapse
Assess the percentage of genes that maintained their trajectory from the full-replicate clustering
Percentage matched 24 clusters
Percentage matched trajectories
Average percentage across replicates
Percentage of genes maintained
MUT_E10.5_1
MUT_E10.5_2
MUT_E10.5_3
MUT_E11.5_1
MUT_E11.5_2
MUT_E11.5_3
MUT_E12.5_1
MUT_E12.5_2
MUT_E12.5_3
MUT_E13.5_1
MUT_E13.5_2
MUT_E13.5_3

**Figure S5. Reproducibility assessment of initial WT and mutant gene expression trajectories, related to Figure 1.** (A) Averaged z-score expression profiles (colored lines) for each collapsed WT and mutant trajectory produced by initial hierarchical clustering. Individual gene expression profiles are shown in grey. Gene counts per trajectory are indicated above each plot. (B) Assessing internal reproducibility of WT clusters and trajectories using a Leave-One-Out (LOO) analysis where a single replicate is removed and clustering is repeated. Cluster assignments generated during each LOO iteration were compared to those obtained using the full dataset. The percentage of WT genes that maintain their original trajectory assignment is reported for both the k = 22 cluster level and the collapsed trajectory level. Mean retention values across all LOO iterations are indicated by dashed lines (light grey, mean cluster retention = 44.3%; dark grey, mean trajectory retention = 49.9%). (C) Assessing internal reproducibility of mutant clusters (k = 24) and trajectories using the same LOO strategy produced comparable mean retention values (light grey, mean cluster retention = 45.6%; dark grey, mean trajectory retention = 48.1%).

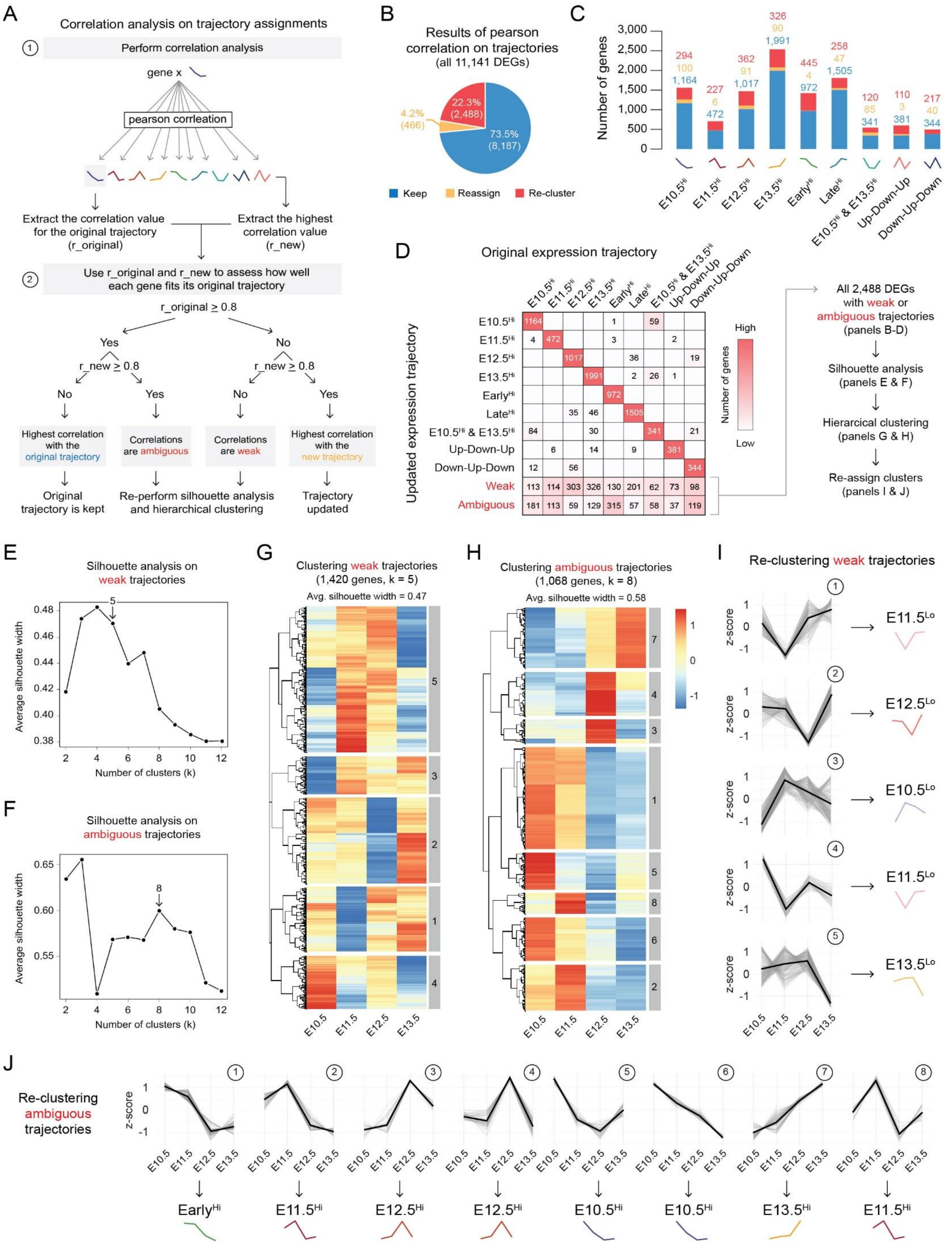

A
Correlation analysis on trajectory assignments
1 Perform correlation analysis
gene x
pearson corrleation
Extract the correlation value for the original trajectory (r_original)
Extract the highest correlation value (r_new)
2 Use r_original and r_new to assess how well each gene fits its original trajectory
r_original ≥ 0.8
Yes
No
r_new ≥ 0.8
No
Yes
Highest correlation with the original trajectory
Correlations are ambiguous
Correlations are weak
Highest correlation with the new trajectory
Original trajectory is kept
Re-perform silhouette analysis and hierarchical clustering
Trajectory updated
B
Results of pearson correlation on trajectories (all 11,141 DEGs)
22.3% (2,488)
4.2% (466)
73.5% (8,187)
Keep
Reassign
Re-cluster
C
Number of genes
E10.5Hi
E11.5Hi
E12.5Hi
E13.5Hi
EarlyHi
LateHi
E10.5Hi & E13.5Hi
Up-Down-Up
Down-Up-Down
D
Original expression trajectory
Updated expression trajectory
Number of genes
High
Low
Weak
Ambiguous
All 2,488 DEGs with weak or ambiguous trajectories (panels B-D)
Silhouette analysis (panels E & F)
Hierarcical clustering (panels G & H)
Re-assign clusters (panels I & J)
E
Silhouette analysis on weak trajectories
Average silhouette width
Number of clusters (k)
F
Silhouette analysis on ambiguous trajectories
G
Clustering weak trajectories (1,420 genes, k = 5)
Avg. silhouette width = 0.47
H
Clustering ambiguous trajectories (1,068 genes, k = 8)
Avg. silhouette width = 0.58
I
Re-clustering weak trajectories
z-score
E11.5Lo
E12.5Lo
E10.5Lo
E11.5Lo
E13.5Lo
J
Re-clustering ambiguous trajectories
EarlyHi
E11.5Hi
E12.5Hi
E12.5Hi
E10.5Hi
E10.5Hi
E13.5Hi
E11.5Hi

**Figure S6. Correlation-based reassessment of WT trajectory assignments identifies weak and ambiguous gene expression profiles, related to Figure 1.** (A) Schematic overview of the correlation-based reassignment strategy used to evaluate confidence in trajectory assignments. For each gene, Pearson correlation coefficients (r) were calculated between its z-scored expression profile and each reference trajectory. The correlation with the originally assigned trajectory (r_original) and the highest correlation observed across all trajectories (r_new) were extracted. These values were then evaluated using a decision tree to determine whether a gene best fit its originally assigned trajectory (r_original $\geq$ 0.8 and r_new < 0.8; original trajectory kept), correlated strongly with multiple trajectories (r_original $\geq$ 0.8 and r_new $\geq$ 0.8; ambiguous), lacked strong correlation to any trajectory (r_original < 0.8 and r_new < 0.8; weak), or correlated more strongly with a different trajectory than the one originally assigned (r_original < 0.8 and r_new $\geq$ 0.8**;** trajectory updated). (B) Summary of correlation outcomes for all WT DEGs (11,141 genes), showing the proportion of genes that kept their original trajectory (Keep), were reassigned to a different trajectory (Reassign), or were flagged for re-clustering due to weak or ambiguous correlations (Re-cluster). (C) Distribution of genes per trajectory, colored by reassignment outcome from panel B, demonstrating that weak and ambiguous assignments are distributed across multiple trajectories rather than concentrated within a single trajectory. (D) Confusion matrix comparing original and updated trajectory assignments following correlation analysis. Numbers indicate the number of genes per comparison; empty boxes denote no genes in that category. Heatmap coloring is scaled per column (red, many genes; white, no genes). Genes classified as weak or ambiguous are highlighted and carried forward for secondary clustering. (E-F) Silhouette analysis performed separately on weak (E) and ambiguous (F) gene sets to determine an appropriate cluster resolution for re-analysis. (G-H) Hierarchical clustering heatmaps of weak (G; 1,420 genes, k = 5) and ambiguous (H; 1,068 genes, k = 8) gene sets, shown as z-scored expression across time. (I-J) Trajectory plots of re-clustered weak (I) and ambiguous (J) gene sets, revealing four additional low-expression trajectories: E10.5 Low, E11.5 Low, E12.5 Low, and E13.5 Low.

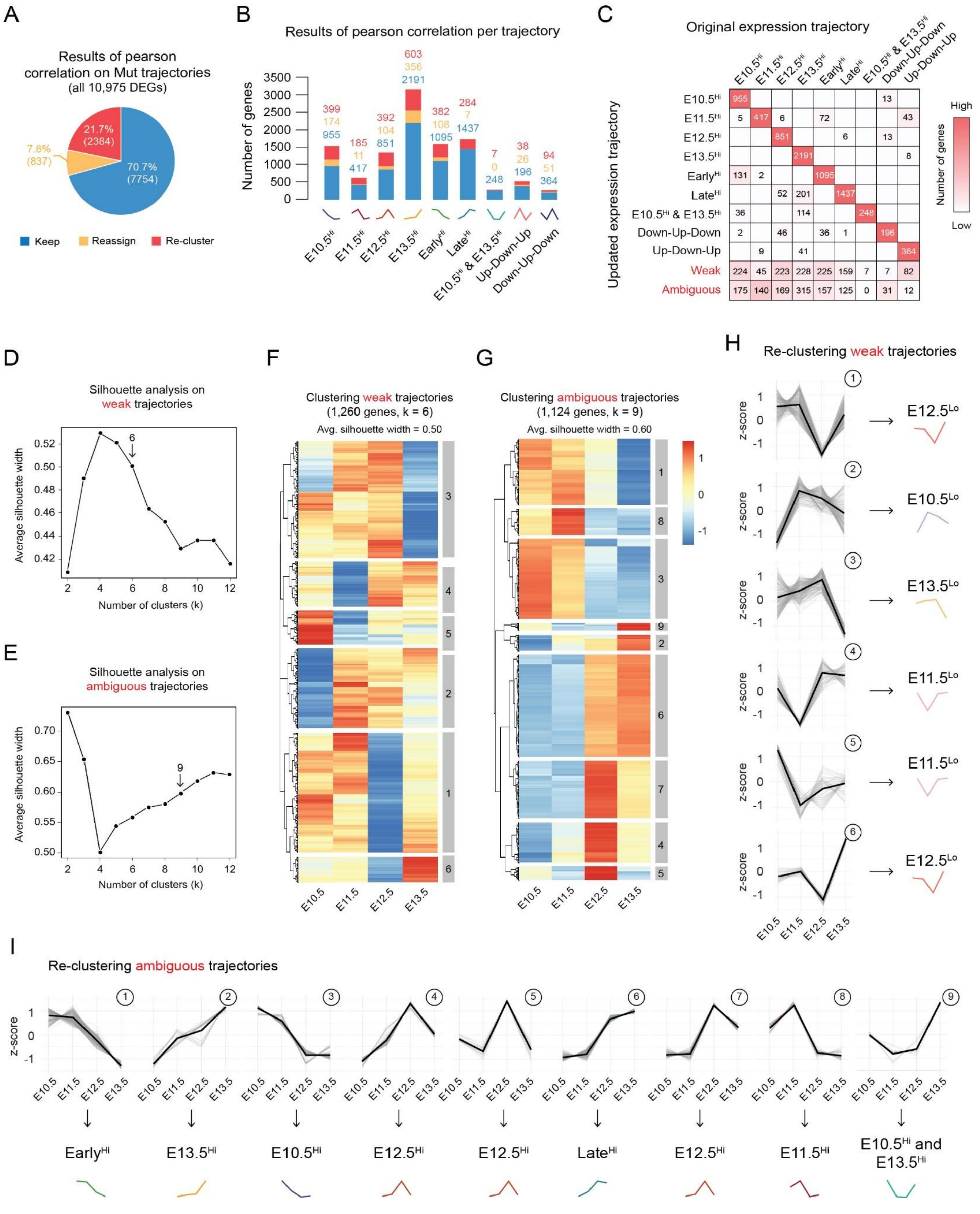
A
Results of pearson correlation on Mut trajectories
(all 10,975 DEGs)
21.7% (2384)
7.6% (837)
70.7% (7754)
Keep
Reassign
Re-cluster
B
Results of pearson correlation per trajectory
Number of genes
E10.5Hi
E11.5Hi
E12.5Hi
E13.5Hi
EarlyHi
LateHi
E10.5Hi & E13.5Hi
Up-Down-Up
Down-Up-Down
C
Original expression trajectory
Updated expression trajectory
Weak
Ambiguous
Number of genes
High
Low
D
Silhouette analysis on weak trajectories
Average silhouette width
Number of clusters (k)
E
Silhouette analysis on ambiguous trajectories
F
Clustering weak trajectories
(1,260 genes, k = 6)
Avg. silhouette width = 0.50
G
Clustering ambiguous trajectories
(1,124 genes, k = 9)
Avg. silhouette width = 0.60
H
Re-clustering weak trajectories
z-score
E12.5Lo
E10.5Lo
E13.5Lo
E11.5Lo
E11.5Lo
E12.5Lo
I
Re-clustering ambiguous trajectories
EarlyHi
E13.5Hi
E10.5Hi
E12.5Hi
E12.5Hi
LateHi
E12.5Hi
E11.5Hi
E10.5Hi and E13.5Hi

**Figure S7. Correlation-based reassessment of mutant trajectory assignments identifies weak and ambiguous expression profiles, related to Figure 1.** (A) Results of the correlation-based trajectory reassignment strategy as previously described in Figure S6A. Pie chart reporting correlation outcomes for all mutant DEGs (10,975 genes), showing the proportion of genes that kept their original trajectory (Keep), were reassigned to a different trajectory (Reassign), or were flagged for re-clustering due to weak or ambiguous correlations (Re-cluster). (B) Distribution of genes per trajectory, colored by reassignment outcome from panel A, demonstrating that weak and ambiguous correlations occur across multiple trajectories rather than originating from a single trajectory. (C) Confusion matrix comparing original and updated trajectory assignments following correlation analysis. Numbers indicate the number of genes per comparison; empty boxes denote no genes in that category. Heatmap coloring was scaled per column (red, many genes; white, no genes). Genes classified as weak or ambiguous are highlighted and carried forward for secondary clustering. (D-E) Silhouette analysis performed separately on weak (D) and ambiguous (E) gene sets to determine an appropriate cluster resolution for re-analysis. (F-G) Hierarchical clustering heatmaps of weak (F; 1,260 genes, k = 6) and ambiguous (G; 1,124 genes, k = 9) gene sets, shown as z-scored expression across time. (H-I) Trajectory plots of re-clustered weak (H) and ambiguous (I) gene sets.

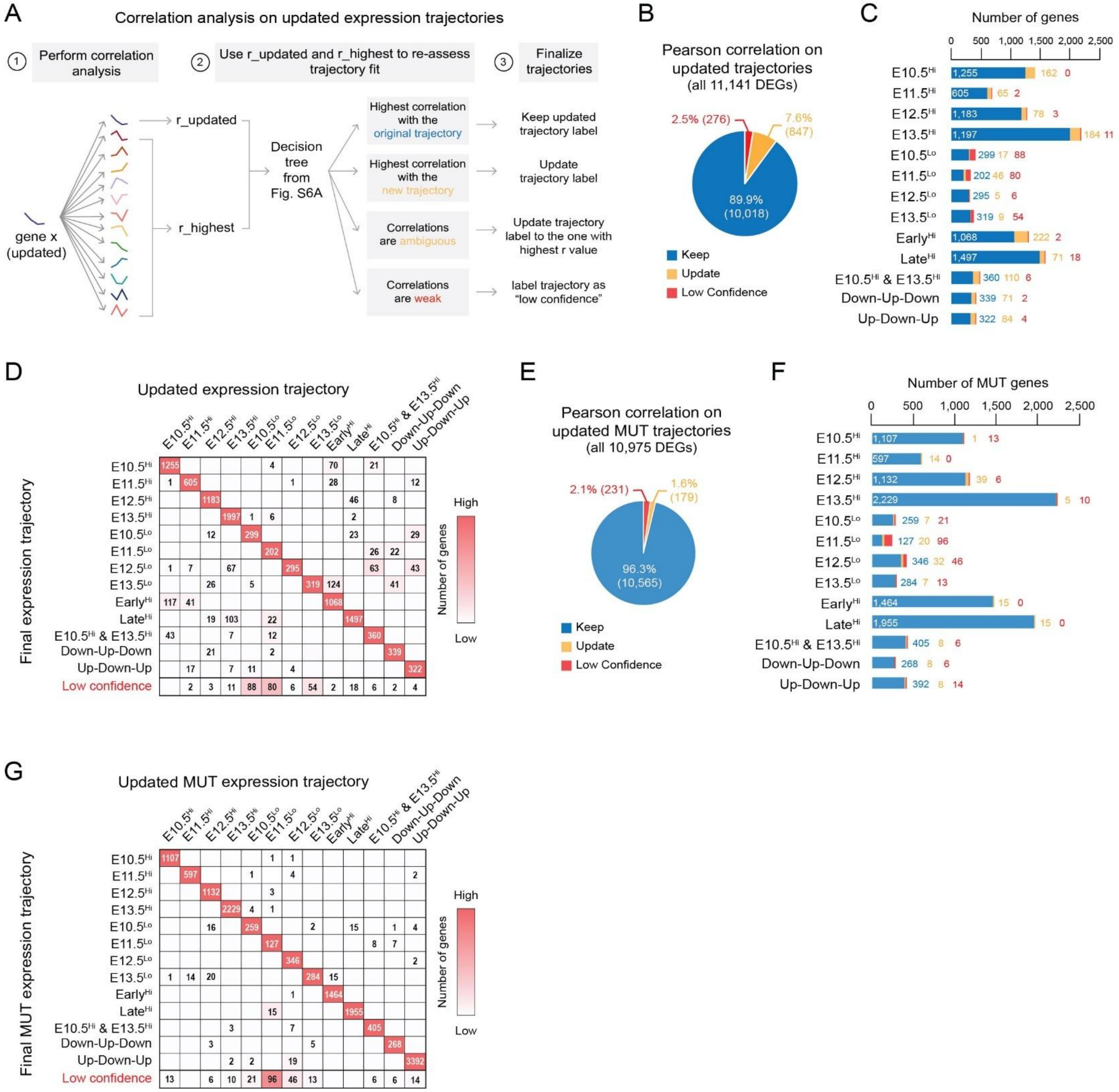

A
Correlation analysis on updated expression trajectories
1 Perform correlation analysis
2 Use r_updated and r_highest to re-assess trajectory fit
3 Finalize trajectories
gene x (updated)
r_updated
r_highest
Decision tree from Fig. S6A
Highest correlation with the original trajectory
Keep updated trajectory label
Highest correlation with the new trajectory
Update trajectory label
Correlations are ambiguous
Update trajectory label to the one with highest r value
Correlations are weak
label trajectory as "low confidence"
B
Pearson correlation on updated trajectories (all 11,141 DEGs)
2.5% (276)
7.6% (847)
89.9% (10,018)
Keep
Update
Low Confidence
C
Number of genes
0 500 1,000 1,500 2,000 2,500
E10.5Hi 1,255 162 0
E11.5Hi 605 65 2
E12.5Hi 1,183 78 3
E13.5Hi 1,197 184 11
E10.5Lo 299 17 88
E11.5Lo 202 46 80
E12.5Lo 295 5 6
E13.5Lo 319 9 54
EarlyHi 1,068 222 2
LateHi 1,497 71 18
E10.5Hi & E13.5Hi 360 110 6
Down-Up-Down 339 71 2
Up-Down-Up 322 84 4
D
Updated expression trajectory
Final expression trajectory
Low confidence
Number of genes
High
Low
E
Pearson correlation on updated MUT trajectories (all 10,975 DEGs)
2.1% (231)
1.6% (179)
96.3% (10,565)
Keep
Update
Low Confidence
F
Number of MUT genes
0 500 1,000 1,500 2,000 2,500
E10.5Hi 1,107 1 13
E11.5Hi 597 14 0
E12.5Hi 1,132 39 6
E13.5Hi 2,229 5 10
E10.5Lo 259 7 21
E11.5Lo 127 20 96
E12.5Lo 346 32 46
E13.5Lo 284 7 13
EarlyHi 1,464 15 0
LateHi 1,955 15 0
E10.5Hi & E13.5Hi 405 8 6
Down-Up-Down 268 8 6
Up-Down-Up 392 8 14
G
Updated MUT expression trajectory
Final MUT expression trajectory
Low confidence
Number of genes
High
Low

**Figure S8. Final correlation-based refinement confirms high-confidence gene expression trajectory assignments in WT and mutant forelimbs, related to Figure 1.** (A) Schematic overview of the final correlation-based strategy applied to updated expression trajectories. For each gene, Pearson correlation coefficients were calculated between its z-scored expression profile and all updated reference trajectories. The correlation with the currently assigned trajectory (r_updated) and the highest correlation observed across all trajectories (r_highest) were extracted and evaluated using the decision framework established in Figure S6A. Genes either retained their updated trajectory, were reassigned to a different trajectory, or were labeled as low confidence if correlations remained weak. Ambiguous trajectories were reassigned to the trajectory with the highest correlation value. (B) Summary of correlation outcomes for all WT DEGs (11,141 genes) following final trajectory assignment, showing the percentage of genes that kept their trajectory assignment (Keep), were reassigned (Update), or were designated as low confidence. (C) Distribution of WT genes per trajectory, colored by reassignment outcome, demonstrating that most genes retained their trajectory assignment following iterative refinement. (D) Confusion matrix comparing the updated and final WT expression trajectory assignments. Numbers indicate the number of genes per comparison; empty boxes denote no genes in that category. Heatmap coloring was scaled per column (red, many genes; white, no genes). Genes classified as low confidence are highlighted and indicate a small subset (276 genes) that does not robustly fit any defined trajectory. (E) Pie chart reporting correlation outcomes for all mutant DEGs (10,975 genes) following final trajectory assignment, showing the percentage of genes that kept their trajectory assignment (Keep), were reassigned (Update), or were designated as low confidence. (F) Distribution of genes per trajectory, colored by reassignment outcome, demonstrating that most mutant genes retained their trajectory assignment following iterative refinement. (G) Confusion matrix comparing the updated and final mutant expression trajectory assignments. Numbers indicate the number of genes per comparison; empty boxes denote no genes in that category. Heatmap coloring was scaled per column (red, many genes; white, no genes). Genes classified as low confidence are highlighted and represent a small subset (231 genes) that do not robustly fit any defined trajectory.

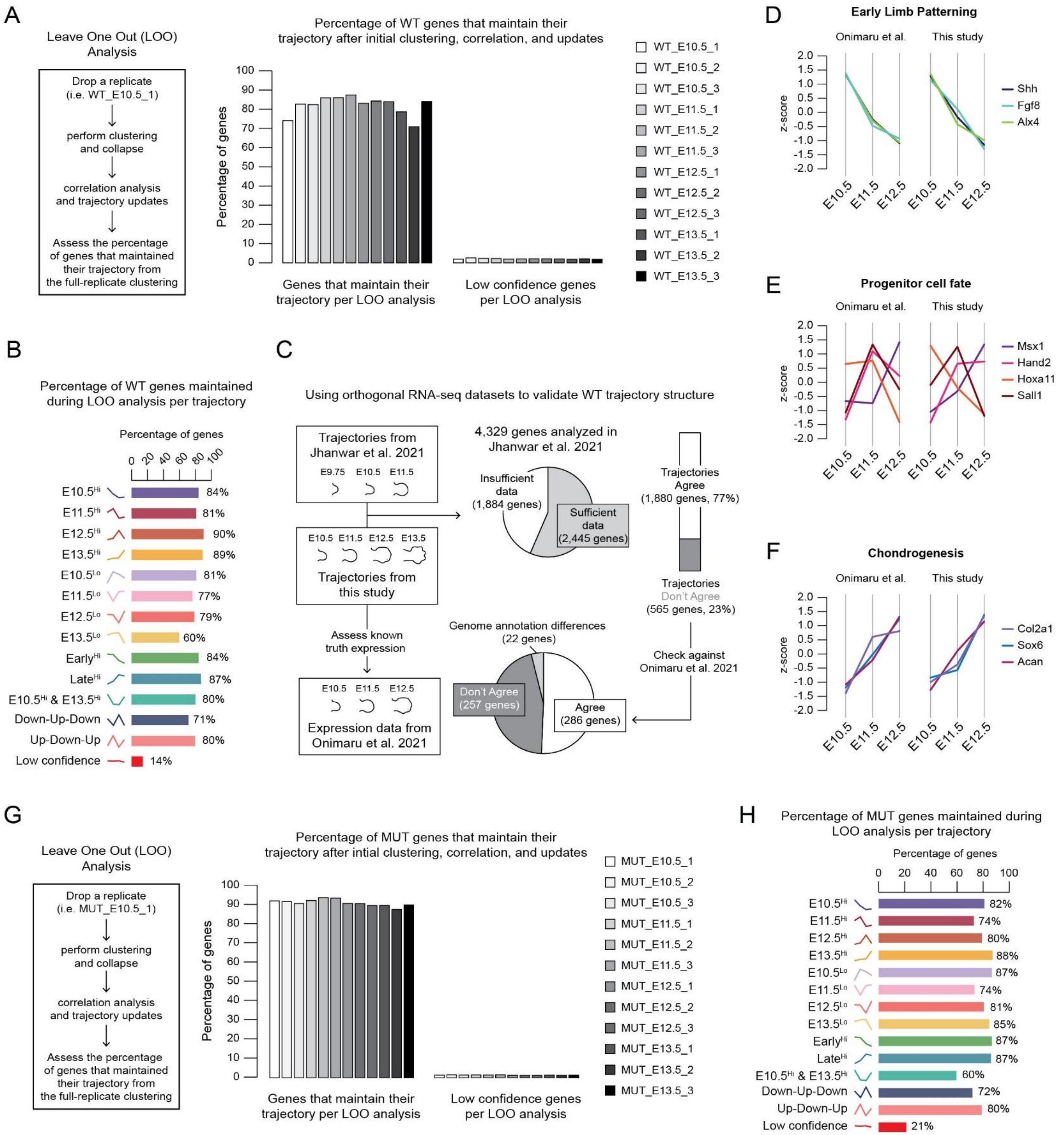
A
Leave One Out (LOO) Analysis
Drop a replicate (i.e. WT_E10.5_1)
perform clustering and collapse
correlation analysis and trajectory updates
Assess the percentage of genes that maintained their trajectory from the full-replicate clustering
Percentage of WT genes that maintain their trajectory after initial clustering, correlation, and updates
Percentage of genes
Genes that maintain their trajectory per LOO analysis
Low confidence genes per LOO analysis
WT_E10.5_1
WT_E10.5_2
WT_E10.5_3
WT_E11.5_1
WT_E11.5_2
WT_E11.5_3
WT_E12.5_1
WT_E12.5_2
WT_E12.5_3
WT_E13.5_1
WT_E13.5_2
WT_E13.5_3
D
Early Limb Patterning
Onimaru et al.
This study
z-score
Shh
Fgf8
Alx4
E10.5
E11.5
E12.5
B
Percentage of WT genes maintained during LOO analysis per trajectory
Percentage of genes
E10.5Hi 84%
E11.5Hi 81%
E12.5Hi 90%
E13.5Hi 89%
E10.5Lo 81%
E11.5Lo 77%
E12.5Lo 79%
E13.5Lo 60%
EarlyHi 84%
LateHi 87%
E10.5Hi & E13.5Hi 80%
Down-Up-Down 71%
Up-Down-Up 80%
Low confidence 14%
C
Using orthogonal RNA-seq datasets to validate WT trajectory structure
Trajectories from Jhanwar et al. 2021
E9.75 E10.5 E11.5
4,329 genes analyzed in Jhanwar et al. 2021
Insufficient data (1,884 genes)
Sufficient data (2,445 genes)
Trajectories Agree (1,880 genes, 77%)
Trajectories Don't Agree (565 genes, 23%)
E10.5 E11.5 E12.5 E13.5
Trajectories from this study
Assess known truth expression
Genome annotation differences (22 genes)
Check against Onimaru et al. 2021
Don't Agree (257 genes)
Agree (286 genes)
E10.5 E11.5 E12.5
Expression data from Onimaru et al. 2021
E
Progenitor cell fate
Msx1
Hand2
Hoxa11
Sall1
F
Chondrogenesis
Col2a1
Sox6
Acan
G
Leave One Out (LOO) Analysis
Drop a replicate (i.e. MUT_E10.5_1)
perform clustering and collapse
correlation analysis and trajectory updates
Assess the percentage of genes that maintained their trajectory from the full-replicate clustering
Percentage of MUT genes that maintain their trajectory after intial clustering, correlation, and updates
Percentage of genes
Genes that maintain their trajectory per LOO analysis
Low confidence genes per LOO analysis
MUT_E10.5_1
MUT_E10.5_2
MUT_E10.5_3
MUT_E11.5_1
MUT_E11.5_2
MUT_E11.5_3
MUT_E12.5_1
MUT_E12.5_2
MUT_E12.5_3
MUT_E13.5_1
MUT_E13.5_2
MUT_E13.5_3
H
Percentage of MUT genes maintained during LOO analysis per trajectory
Percentage of genes
E10.5Hi 82%
E11.5Hi 74%
E12.5Hi 80%
E13.5Hi 88%
E10.5Lo 87%
E11.5Lo 74%
E12.5Lo 81%
E13.5Lo 85%
EarlyHi 87%
LateHi 87%
E10.5Hi & E13.5Hi 60%
Down-Up-Down 72%
Up-Down-Up 80%
Low confidence 21%

**Figure S9. Validation of final expression trajectory assignments in WT and mutant forelimbs, related to Figure 1.** (A) Schematic overview of LOO analysis used to assess reproducibility of final WT trajectory assignments. For each LOO iteration, a single replicate was removed and clustering was repeated. Cluster assignments generated during each LOO iteration were compared to those obtained using the full dataset. The percentage of genes that maintained their final trajectory assignment was calculated per replicate, and the percentage of low confidence genes generated in each iteration is also reported. (B) Percentage of WT genes that maintained their trajectory assignment per trajectory across LOO iterations. Most trajectories showed high retention, indicating robust and reproducible trajectory structure. (C) Validation of WT trajectory structure using two independent, orthogonal RNA-seq datasets. Trajectories defined in this study were compared with published data from Jhanwar et al. and Onimaru et al.[1,2]. Of the 4,329 genes analyzed in Jhanwar et al., 2,445 had sufficient timepoint overlap for comparison. Among these, 77% showed concordant trajectory structure between studies, while 23% did not. Discordant trajectories were cross-checked against the Onimaru et al. dataset, revealing that approximately half of the genes that disagreed with Jhanwar et al. were concordant with Onimaru et al. (50.6%, 286 genes). Discrepancies in trajectory assignments were consistent with differences in genome annotation, sequencing depth, and timepoint variability between studies[1,2]. (D-F) Comparison of representative gene expression trajectories associated with known developmental processes between published data and this study[2]. Early limb patterning genes (D), regulators of progenitor cell fate (E), and chondrogenic markers (F) exhibit highly similar temporal expression profiles across datasets, supporting the reproducibility of the present dataset and the biological relevance of the identified trajectories. (G) Schematic overview of LOO analysis used to assess reproducibility of final mutant trajectory assignments. (H) Percentage of genes that maintained their mutant trajectory assignment per trajectory across LOO iterations. Most mutant trajectories showed high retention, indicating robust and reproducible trajectory structure.

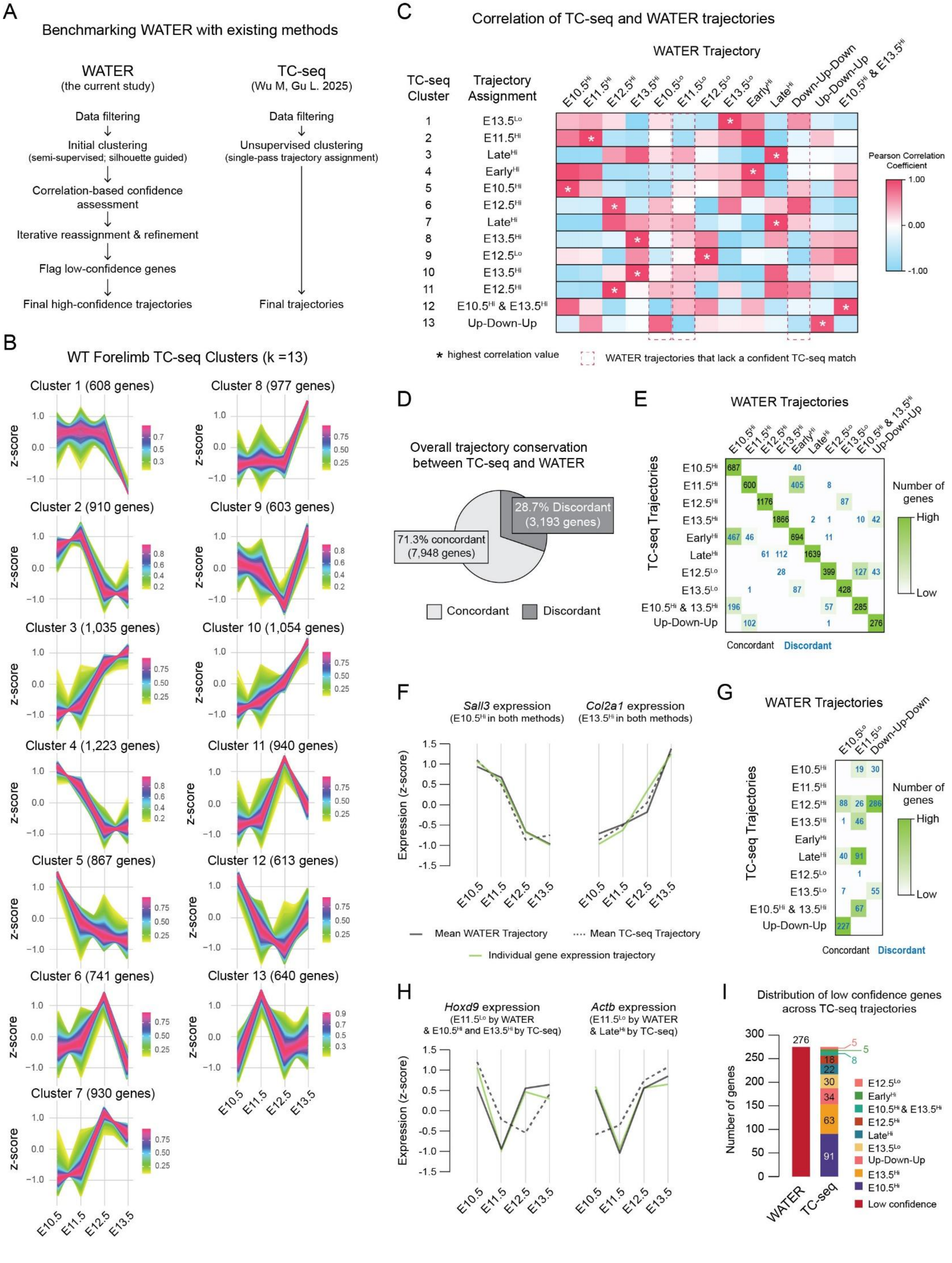

A
Benchmarking WATER with existing methods
WATER
(the current study)
Data filtering
Initial clustering
(semi-supervised; silhouette guided)
Correlation-based confidence assessment
Iterative reassignment & refinement
Flag low-confidence genes
Final high-confidence trajectories
TC-seq
(Wu M, Gu L. 2025)
Data filtering
Unsupervised clustering
(single-pass trajectory assignment)
Final trajectories
B
WT Forelimb TC-seq Clusters (k =13)
Cluster 1 (608 genes)
Cluster 2 (910 genes)
Cluster 3 (1,035 genes)
Cluster 4 (1,223 genes)
Cluster 5 (867 genes)
Cluster 6 (741 genes)
Cluster 7 (930 genes)
Cluster 8 (977 genes)
Cluster 9 (603 genes)
Cluster 10 (1,054 genes)
Cluster 11 (940 genes)
Cluster 12 (613 genes)
Cluster 13 (640 genes)
z-score
E10.5
E11.5
E12.5
E13.5
C
Correlation of TC-seq and WATER trajectories
WATER Trajectory
TC-seq Cluster
Trajectory Assignment
E10.5Hi
E11.5Hi
E12.5Hi
E13.5Hi
E10.5Lo
E11.5Lo
E12.5Lo
E13.5Lo
EarlyHi
LateHi
Down-Up-Down
Up-Down-Up
E10.5Hi & E13.5Hi
1 E13.5Lo
2 E11.5Hi
3 LateHi
4 EarlyHi
5 E10.5Hi
6 E12.5Hi
7 LateHi
8 E13.5Hi
9 E12.5Lo
10 E13.5Hi
11 E12.5Hi
12 E10.5Hi & E13.5Hi
13 Up-Down-Up
Pearson Correlation Coefficient
1.00
0.00
-1.00
* highest correlation value
WATER trajectories that lack a confident TC-seq match
D
Overall trajectory conservation between TC-seq and WATER
28.7% Discordant (3,193 genes)
71.3% concordant (7,948 genes)
Concordant
Discordant
E
WATER Trajectories
TC-seq Trajectories
Number of genes
High
Low
Concordant
Discordant
F
Sall3 expression
(E10.5Hi in both methods)
Col2a1 expression
(E13.5Hi in both methods)
Expression (z-score)
Mean WATER Trajectory
Mean TC-seq Trajectory
Individual gene expression trajectory
G
WATER Trajectories
TC-seq Trajectories
Number of genes
High
Low
Concordant
Discordant
H
Hoxd9 expression
(E11.5Lo by WATER & E10.5Hi and E13.5Hi by TC-seq)
Actb expression
(E11.5Lo by WATER & LateHi by TC-seq)
Expression (z-score)
I
Distribution of low confidence genes across TC-seq trajectories
Number of genes
WATER
TC-seq
276
91
63
34
30
22
18
8
5
5
E12.5Lo
EarlyHi
E10.5Hi & E13.5Hi
E12.5Hi
LateHi
E13.5Lo
Up-Down-Up
E13.5Hi
E10.5Hi
Low confidence

**Figure S10. Benchmarking WATER against TC-seq for high confidence temporal trajectory inference, related to Figure 1.** (A) Schematic comparison of WATER and TC-seq workflows[3]. WATER utilizes semi-supervised clustering, correlation-based confidence assessment, iterative refinement, and tagging of low confidence genes to generate high confidence temporal trajectories. In contrast, TC-seq applies unsupervised single-pass clustering to assign genes to temporal expression clusters. (B) WT forelimb TC-seq clusters (k = 13). Z-scored gene expression across E10.5 to E13.5 for each TC-seq cluster. Each line represents the expression profile of an individual gene, with colors indicating membership value (i.e., the strength of a gene's association with a given cluster). Higher membership values denote stronger cluster assignment. (C) Pearson correlation analysis between TC-seq and WATER trajectories. The highest correlation for each TC-seq cluster is indicated by (*). Correlation values were used to match TC-seq and WATER trajectories. Red dashed boxes indicate WATER trajectories that lack a confident TC-seq match. (D) Overall trajectory conservation between TC-seq and WATER shows most genes (7,948 genes; 71.3%) are concordant between the two methods. Conversely, 28.7% (3,193 genes) were discordant between TC-seq and WATER. (E) Confusion matrix showing gene overlap between TC-seq clusters and WATER trajectories. Numbers indicate the number of genes per comparison; empty boxes denote no genes in that category. Heatmap coloring was scaled per column (green, many genes; white, no genes). Gene sets considered discordant between the two methods are colored in blue. (F) Representative expression trajectories for two genes, *Sall3* and *Col2a1*, whose trajectories are concordant between WATER and TC-seq. Plots compare the mean WATER (solid line) and TC-seq (dashed line) trajectory profiles for the cluster assigned to each gene, with the observed gene expression overlaid (green line). (G) Confusion matrix for WATER trajectories lacking a confident TC-seq match, highlighting the distribution of these genes across multiple TC-seq clusters. Numbers indicate the number of genes per comparison; empty boxes denote no genes in that category. Heatmap coloring was scaled per column (green, many genes; white, no genes). Gene sets considered discordant between the two methods are colored in blue. (H) Representative expression trajectories for *Hoxd9* and *Actb*, whose trajectories are discordant between WATER and TC-seq. Plots compare the mean WATER (solid line) and TC-seq (dashed line) trajectory profiles for the cluster assigned to each gene, with the observed gene expression overlaid (green line). (I) Distribution of low confidence genes across TC-seq trajectories, highlighting the TC-seq clustering of genes flagged as low confidence by WATER. See also Table S2.

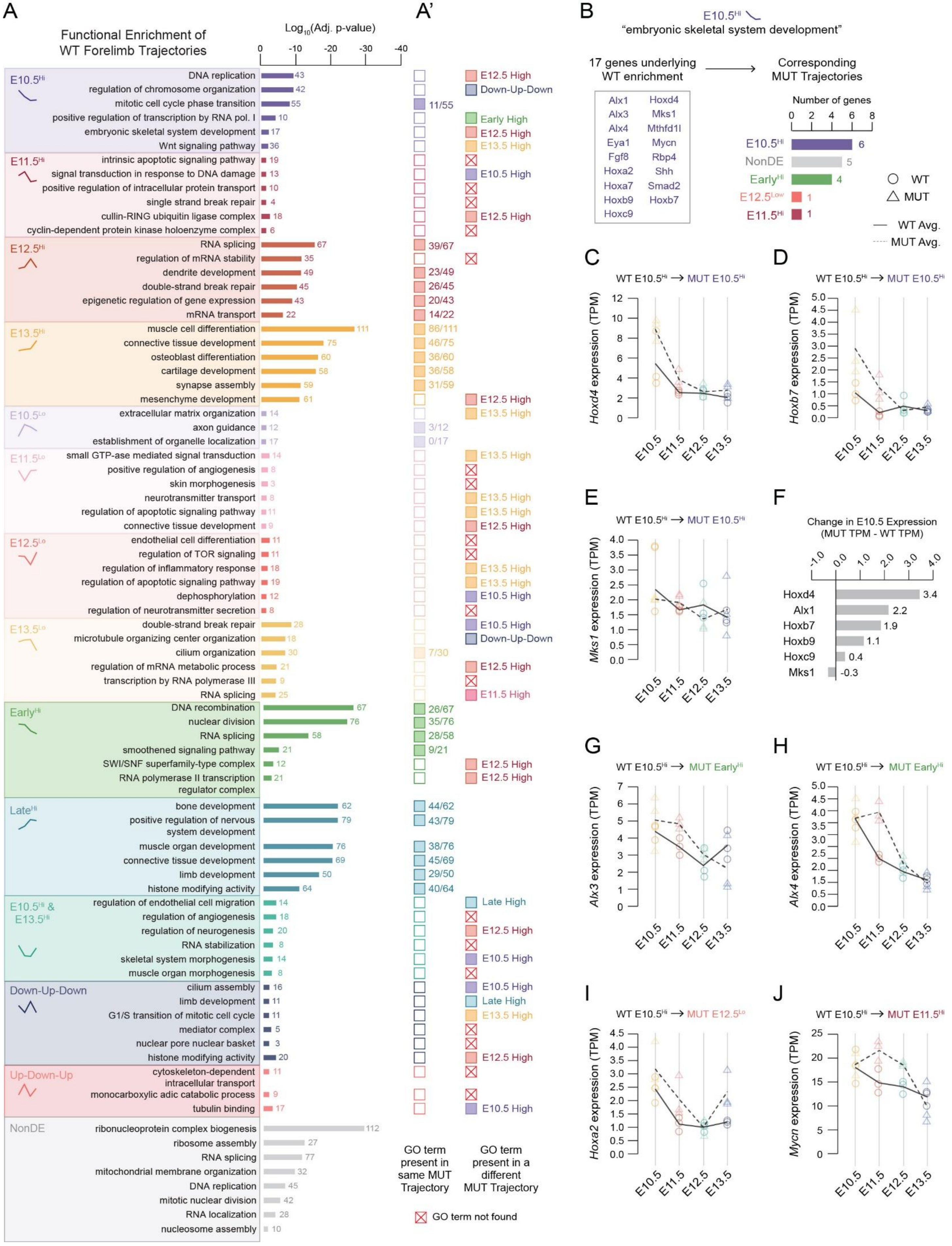
A
Functional Enrichment of WT Forelimb Trajectories
Log10(Adj. p-value)
E10.5Hi
DNA replication
regulation of chromosome organization
mitotic cell cycle phase transition
positive regulation of transcription by RNA pol. I
embryonic skeletal system development
Wnt signaling pathway
E11.5Hi
intrinsic apoptotic signaling pathway
signal transduction in response to DNA damage
positive regulation of intracellular protein transport
single strand break repair
cullin-RING ubiquitin ligase complex
cyclin-dependent protein kinase holoenzyme complex
E12.5Hi
RNA splicing
regulation of mRNA stability
dendrite development
double-strand break repair
epigenetic regulation of gene expression
mRNA transport
E13.5Hi
muscle cell differentiation
connective tissue development
osteoblast differentiation
cartilage development
synapse assembly
mesenchyme development
E10.5Lo
extracellular matrix organization
axon guidance
establishment of organelle localization
E11.5Lo
small GTP-ase mediated signal transduction
positive regulation of angiogenesis
skin morphogenesis
neurotransmitter transport
regulation of apoptotic signaling pathway
connective tissue development
E12.5Lo
endothelial cell differentiation
regulation of TOR signaling
regulation of inflammatory response
regulation of apoptotic signaling pathway
dephosphorylation
regulation of neurotransmitter secretion
E13.5Lo
double-strand break repair
microtubule organizing center organization
cilium organization
regulation of mRNA metabolic process
transcription by RNA polymerase III
RNA splicing
EarlyHi
DNA recombination
nuclear division
RNA splicing
smoothened signaling pathway
SWI/SNF superfamily-type complex
RNA polymerase II transcription regulator complex
LateHi
bone development
positive regulation of nervous system development
muscle organ development
connective tissue development
limb development
histone modifying activity
E10.5Hi & E13.5Hi
regulation of endothelial cell migration
regulation of angiogenesis
regulation of neurogenesis
RNA stabilization
skeletal system morphogenesis
muscle organ morphogenesis
Down-Up-Down
cilium assembly
limb development
G1/S transition of mitotic cell cycle
mediator complex
nuclear pore nuclear basket
histone modifying activity
Up-Down-Up
cytoskeleton-dependent intracellular transport
monocarboxylic adic catabolic process
tubulin binding
NonDE
ribonucleoprotein complex biogenesis
ribosome assembly
RNA splicing
mitochondrial membrane organization
DNA replication
mitotic nuclear division
RNA localization
nucleosome assembly
A'
GO term present in same MUT Trajectory
GO term present in a different MUT Trajectory
GO term not found
B
E10.5Hi
"embryonic skeletal system development"
17 genes underlying WT enrichment
Corresponding MUT Trajectories
Alx1 Hoxd4 Alx3 Mks1 Alx4 Mthfd1l Eya1 Mycn Fgf8 Rbp4 Hoxa2 Shh Hoxa7 Smad2 Hoxb9 Hoxb7 Hoxc9
Number of genes
E10.5Hi 6
NonDE 5
EarlyHi 4
E12.5Low 1
E11.5Hi 1
WT
MUT
WT Avg.
MUT Avg.
C
WT E10.5Hi → MUT E10.5Hi
Hoxd4 expression (TPM)
D
WT E10.5Hi → MUT E10.5Hi
Hoxb7 expression (TPM)
E
WT E10.5Hi → MUT E10.5Hi
Mks1 expression (TPM)
F
Change in E10.5 Expression (MUT TPM - WT TPM)
Hoxd4 3.4
Alx1 2.2
Hoxb7 1.9
Hoxb9 1.1
Hoxc9 0.4
Mks1 -0.3
G
WT E10.5Hi → MUT EarlyHi
Alx3 expression (TPM)
H
WT E10.5Hi → MUT EarlyHi
Alx4 expression (TPM)
I
WT E10.5Hi → MUT E12.5Lo
Hoxa2 expression (TPM)
J
WT E10.5Hi → MUT E11.5Hi
Mycn expression (TPM)
E10.5 E11.5 E12.5 E13.5

**Figure S11. Functional enrichment of expression trajectories reveals differences between WT and mutant forelimbs, related to Figure 1.** (A) Gene Ontology (GO; BP, CC, and MF) enrichment analysis for each WT expression trajectory. Bar plots show significantly enriched terms (adjusted $p < 0.05$), grouped by trajectory and colored to match corresponding WT expression patterns. Bar length indicates $\log_{10}$(adjusted p-value), and the number of genes underlying each enrichment is indicated at the end of the corresponding bar. (A') Filled squares indicate the same GO term is enriched in the corresponding mutant trajectory, whereas open squares indicate that the term was not enriched in that corresponding mutant trajectory. When a GO term is enriched in the mutant, the number of genes shared between the WT and mutant is shown. If the term is enriched in the mutant but assigned to a different trajectory, the corresponding mutant trajectory is indicated next to the open square to denote redistribution. Red squares marked with an "X" denote GO terms that were not enriched in any mutant trajectory. Together, this panel illustrates redistribution of functional programs across temporal expression trajectories following U11 loss. (B) Genes underlying WT enrichment for the GO term "embryonic skeletal system development" within the WT E10.5 high trajectory (17 genes). Bars indicate the number of these genes assigned to each mutant trajectory, demonstrating partial conservation of early skeletal development programming alongside redistribution to alternative temporal trajectories in the mutant. (C-E) Representative examples of genes that remain assigned to the E10.5 high trajectory in both WT and mutant. Lines show the mean gene expression (TPM) across time (E10.5-E13.5) for WT (solid line, circles) and mutant (dashed line, triangles), with biological replicates overlaid. (F) Change in E10.5 expression (MUT TPM – WT TPM) for genes that maintain an E10.5 high trajectory in both genotypes. Positive values indicate higher expression in the mutant, whereas negative values indicate higher expression in the WT, revealing changes in expression magnitude despite trajectory conservation. (G-J) Examples of skeletal development genes that shift from WT E10.5 high to mutant Early high (G-H), E12.5 Low (I), and E11.5 High (J). See also data S5-6.

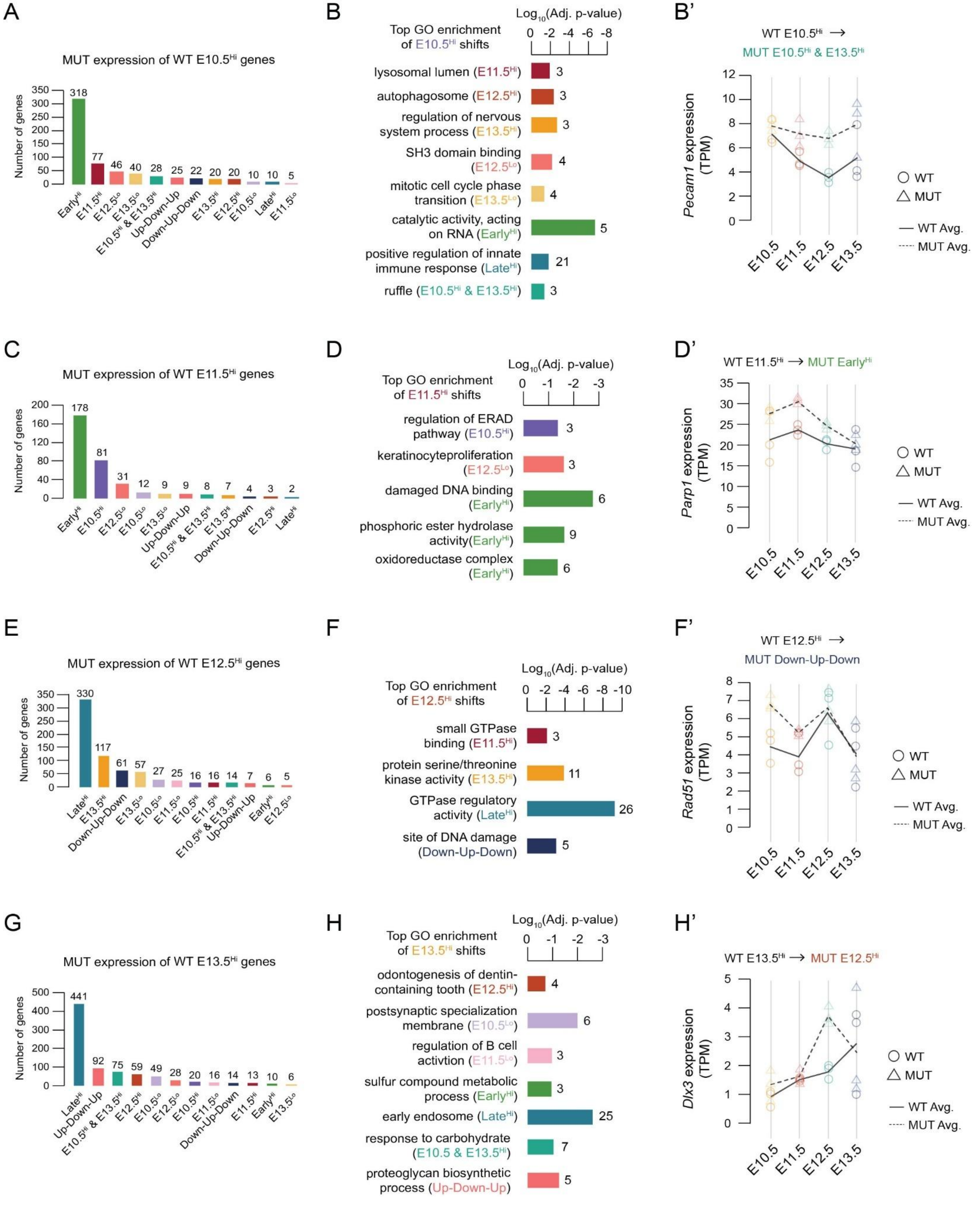
A
MUT expression of WT E10.5Hi genes
Number of genes
318 77 46 40 28 25 22 20 20 10 10 5
EarlyHi E11.5Hi E12.5Lo E13.5Lo E10.5Hi & E13.5Hi Up-Down-Up Down-Up-Down E13.5Hi E12.5Hi E10.5Lo LateHi E11.5Lo
B
Top GO enrichment of E10.5Hi shifts
Log10(Adj. p-value)
lysosomal lumen (E11.5Hi) 3
autophagosome (E12.5Hi) 3
regulation of nervous system process (E13.5Hi) 3
SH3 domain binding (E12.5Lo) 4
mitotic cell cycle phase transition (E13.5Lo) 4
catalytic activity, acting on RNA (EarlyHi) 5
positive regulation of innate immune response (LateHi) 21
ruffle (E10.5Hi & E13.5Hi) 3
B'
WT E10.5Hi → MUT E10.5Hi & E13.5Hi
Pecam1 expression (TPM)
WT
MUT
WT Avg.
MUT Avg.
E10.5 E11.5 E12.5 E13.5
C
MUT expression of WT E11.5Hi genes
Number of genes
178 81 31 12 9 9 8 7 4 3 2
EarlyHi E10.5Hi E12.5Lo E10.5Lo E13.5Lo Up-Down-Up E10.5Hi & E13.5Hi E13.5Hi Down-Up-Down E12.5Hi LateHi
D
Top GO enrichment of E11.5Hi shifts
Log10(Adj. p-value)
regulation of ERAD pathway (E10.5Hi) 3
keratinocyteproliferation (E12.5Lo) 3
damaged DNA binding (EarlyHi) 6
phosphoric ester hydrolase activity(EarlyHi) 9
oxidoreductase complex (EarlyHi) 6
D'
WT E11.5Hi → MUT EarlyHi
Parp1 expression (TPM)
WT
MUT
WT Avg.
MUT Avg.
E
MUT expression of WT E12.5Hi genes
Number of genes
330 117 61 57 27 25 16 16 14 7 6 5
LateHi E13.5Hi Down-Up-Down E13.5Lo E10.5Lo E11.5Lo E10.5Hi E11.5Hi E10.5Hi & E13.5Hi Up-Down-Up EarlyHi E12.5Lo
F
Top GO enrichment of E12.5Hi shifts
Log10(Adj. p-value)
small GTPase binding (E11.5Hi) 3
protein serine/threonine kinase activity (E13.5Hi) 11
GTPase regulatory activity (LateHi) 26
site of DNA damage (Down-Up-Down) 5
F'
WT E12.5Hi → MUT Down-Up-Down
Rad51 expression (TPM)
WT
MUT
WT Avg.
MUT Avg.
G
MUT expression of WT E13.5Hi genes
Number of genes
441 92 75 59 49 28 20 16 14 13 10 6
LateHi Up-Down-Up E10.5Hi & E13.5Hi E12.5Hi E10.5Lo E12.5Lo E10.5Hi E11.5Lo Down-Up-Down E11.5Hi EarlyHi E13.5Lo
H
Top GO enrichment of E13.5Hi shifts
Log10(Adj. p-value)
odontogenesis of dentin-containing tooth (E12.5Hi) 4
postsynaptic specialization membrane (E10.5Lo) 6
regulation of B cell activtion (E11.5Lo) 3
sulfur compound metabolic process (EarlyHi) 3
early endosome (LateHi) 25
response to carbohydrate (E10.5 & E13.5Hi) 7
proteoglycan biosynthetic process (Up-Down-Up) 5
H'
WT E13.5Hi → MUT E12.5Hi
Dlx3 expression (TPM)
WT
MUT
WT Avg.
MUT Avg.

**Figure S12. Redistribution of High WT-only expression trajectories following U11 loss, related to Figure 2.** (A) Distribution of mutant trajectory assignments for genes that exclusively follow the WT E10.5 high trajectory. Bars indicate the number of genes reassigned to each mutant trajectory. (B) Top GO enrichment results for each E10.5 high WT-to-MUT trajectory shift. The absence of a trajectory indicates that shift had no functional enrichment results. Bars represent significantly enriched GO terms (BP, CC, MF; adjusted $p < 0.05$), with bar length indicating the $\log_{10}$(adjusted p-value) and numbers denoting the number of genes underlying enrichment of each term. (B') Example expression profile for *Pecam1*, underlying the GO term "ruffle" in panel B. Lines show the mean expression (TPM) across time for WT (solid line, circles) and mutant (dashed line, triangles), with biological replicates overlaid. (C) Distribution of mutant trajectory assignments for genes that exclusively follow the WT E11.5 high trajectory. (D) Top GO enrichment results for each E11.5 high WT-to-MUT trajectory shift. (D') Example expression profile for *Parp1*, underlying the GO term "damaged DNA binding" in panel D. (E) Distribution of mutant trajectory assignments for genes that exclusively follow the WT E12.5 high trajectory. (F) Top GO enrichment results for each E12.5 high WT-to-MUT trajectory shift. (F') Example expression profile for *Rad51*, underlying the GO term "site of DNA damage" in panel F. (G) Distribution of mutant trajectory assignments for genes that exclusively follow the WT E13.5 high trajectory. (H) Top GO enrichment results for each E13.5 high WT-to-MUT trajectory shift. (H') Example expression profile for *Dlx3*, underlying the GO term "odontogenesis of dentin-containing tooth" in panel H.

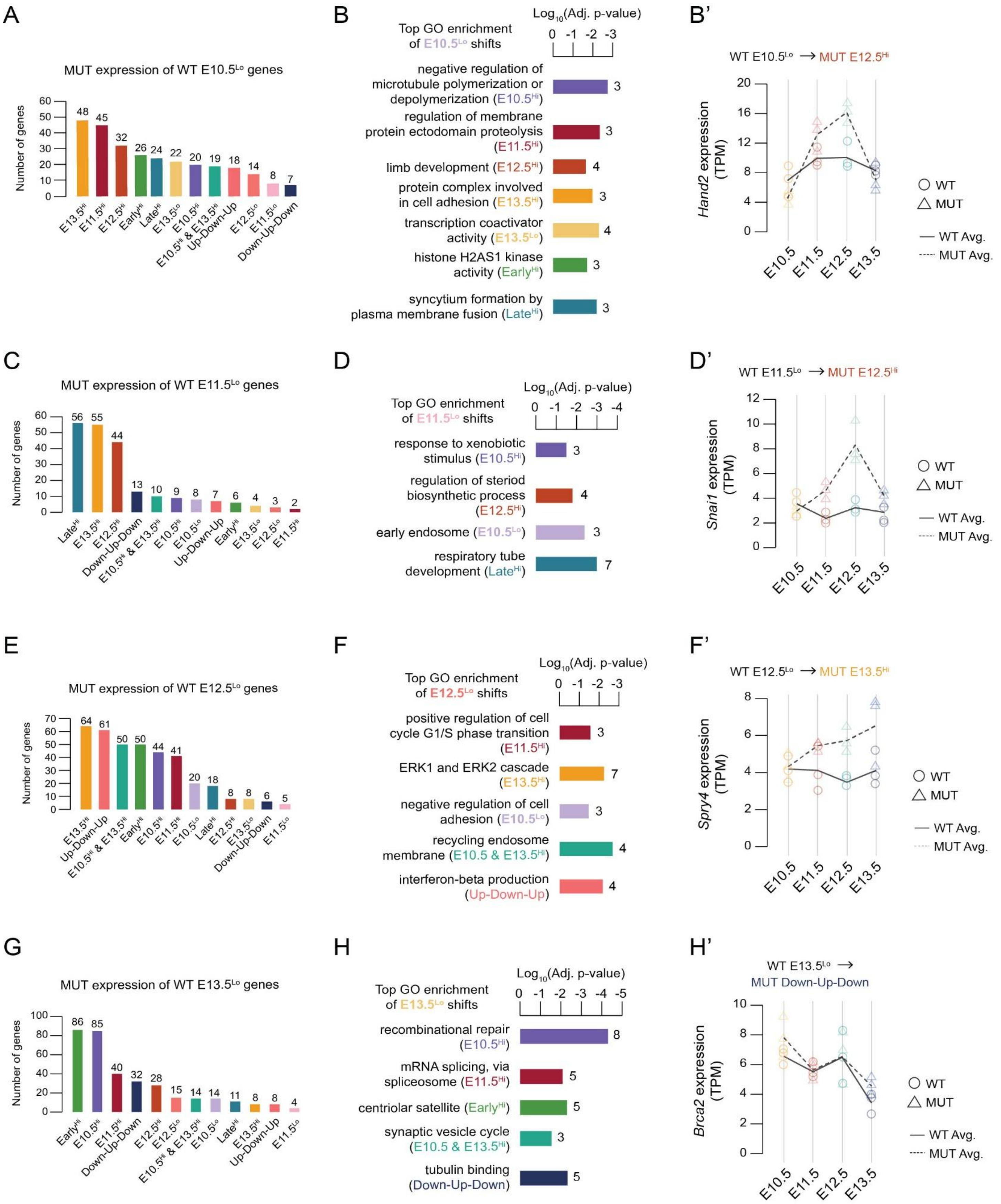
A
MUT expression of WT E10.5Lo genes
Number of genes
48 45 32 26 24 22 20 19 18 14 8 7
E13.5Hi E11.5Hi E12.5Hi EarlyHi LateHi E13.5Lo E10.5Hi E10.5Hi & E13.5Hi Up-Down-Up E12.5Lo E11.5Lo Down-Up-Down
B
Top GO enrichment of E10.5Lo shifts
Log10(Adj. p-value)
negative regulation of microtubule polymerization or depolymerization (E10.5Hi)
regulation of membrane protein ectodomain proteolysis (E11.5Hi)
limb development (E12.5Hi)
protein complex involved in cell adhesion (E13.5Hi)
transcription coactivator activity (E13.5Lo)
histone H2AS1 kinase activity (EarlyHi)
syncytium formation by plasma membrane fusion (LateHi)
B'
WT E10.5Lo → MUT E12.5Hi
Hand2 expression (TPM)
WT
MUT
WT Avg.
MUT Avg.
C
MUT expression of WT E11.5Lo genes
Number of genes
56 55 44 13 10 9 8 7 6 4 3 2
LateHi E13.5Hi E12.5Hi Down-Up-Down E10.5Hi & E13.5Hi E10.5Hi E10.5Lo Up-Down-Up EarlyHi E13.5Lo E12.5Lo E11.5Hi
D
Top GO enrichment of E11.5Lo shifts
Log10(Adj. p-value)
response to xenobiotic stimulus (E10.5Hi)
regulation of steriod biosynthetic process (E12.5Hi)
early endosome (E10.5Lo)
respiratory tube development (LateHi)
D'
WT E11.5Lo → MUT E12.5Hi
Snai1 expression (TPM)
E
MUT expression of WT E12.5Lo genes
Number of genes
64 61 50 50 44 41 20 18 8 8 6 5
E13.5Hi Up-Down-Up E10.5Hi & E13.5Hi EarlyHi E10.5Hi E11.5Hi E10.5Lo LateHi E12.5Hi E13.5Lo Down-Up-Down E11.5Lo
F
Top GO enrichment of E12.5Lo shifts
Log10(Adj. p-value)
positive regulation of cell cycle G1/S phase transition (E11.5Hi)
ERK1 and ERK2 cascade (E13.5Hi)
negative regulation of cell adhesion (E10.5Lo)
recycling endosome membrane (E10.5 & E13.5Hi)
interferon-beta production (Up-Down-Up)
F'
WT E12.5Lo → MUT E13.5Hi
Spry4 expression (TPM)
G
MUT expression of WT E13.5Lo genes
Number of genes
86 85 40 32 28 15 14 14 11 8 8 4
EarlyHi E10.5Hi E11.5Hi Down-Up-Down E12.5Hi E12.5Lo E10.5Hi & E13.5Hi E10.5Lo LateHi E13.5Hi Up-Down-Up E11.5Lo
H
Top GO enrichment of E13.5Lo shifts
Log10(Adj. p-value)
recombinational repair (E10.5Hi)
mRNA splicing, via spliceosome (E11.5Hi)
centriolar satellite (EarlyHi)
synaptic vesicle cycle (E10.5 & E13.5Hi)
tubulin binding (Down-Up-Down)
H'
WT E13.5Lo → MUT Down-Up-Down
Brca2 expression (TPM)
E10.5 E11.5 E12.5 E13.5

**Figure S13. Redistribution of Low WT-only expression trajectories following U11 loss, related to Figure 2.** (A) Distribution of mutant trajectory assignments for genes that exclusively follow the WT E10.5 low trajectory. Bars indicate the number of genes reassigned to each mutant trajectory. (B) Top GO enrichment results for each E10.5 low WT-to-MUT trajectory shift. The absence of a trajectory indicates that shift had no functional enrichment results. Bars represent significantly enriched GO terms (BP, CC, MF; adjusted $p < 0.05$), with bar length indicating the $\log_{10}$(adjusted p-value) and numbers denoting the number of genes underlying enrichment of each term. (B') Example expression profile for *Hand2*, underlying the GO term "limb development" in panel B. Lines show the mean expression (TPM) across time for WT (solid line, circles) and mutant (dashed line, triangles), with biological replicates overlaid. (C) Distribution of mutant trajectory assignments for genes that exclusively follow the WT E11.5 low trajectory. (D) Top GO enrichment results for each E11.5 low WT-to-MUT trajectory shift. (D') Example expression profile for *Snai1*, underlying the GO term "regulation of steroid biosynthetic process" in panel D. (E) Distribution of mutant trajectory assignments for genes that exclusively follow the WT E12.5 low trajectory. (F) Top GO enrichment results for each E12.5 low WT-to-MUT trajectory shift. (F') Example expression profile for *Spry4*, underlying the GO term "ERK1 and ERK2 cascade" in panel F. (G) Distribution of mutant trajectory assignments for genes that exclusively follow the WT E13.5 low trajectory. (H) Top GO enrichment results for each E13.5 low WT-to-MUT trajectory shift. (H') Example expression profile for *Brca2*, underlying the GO term "tubulin binding" in panel H.

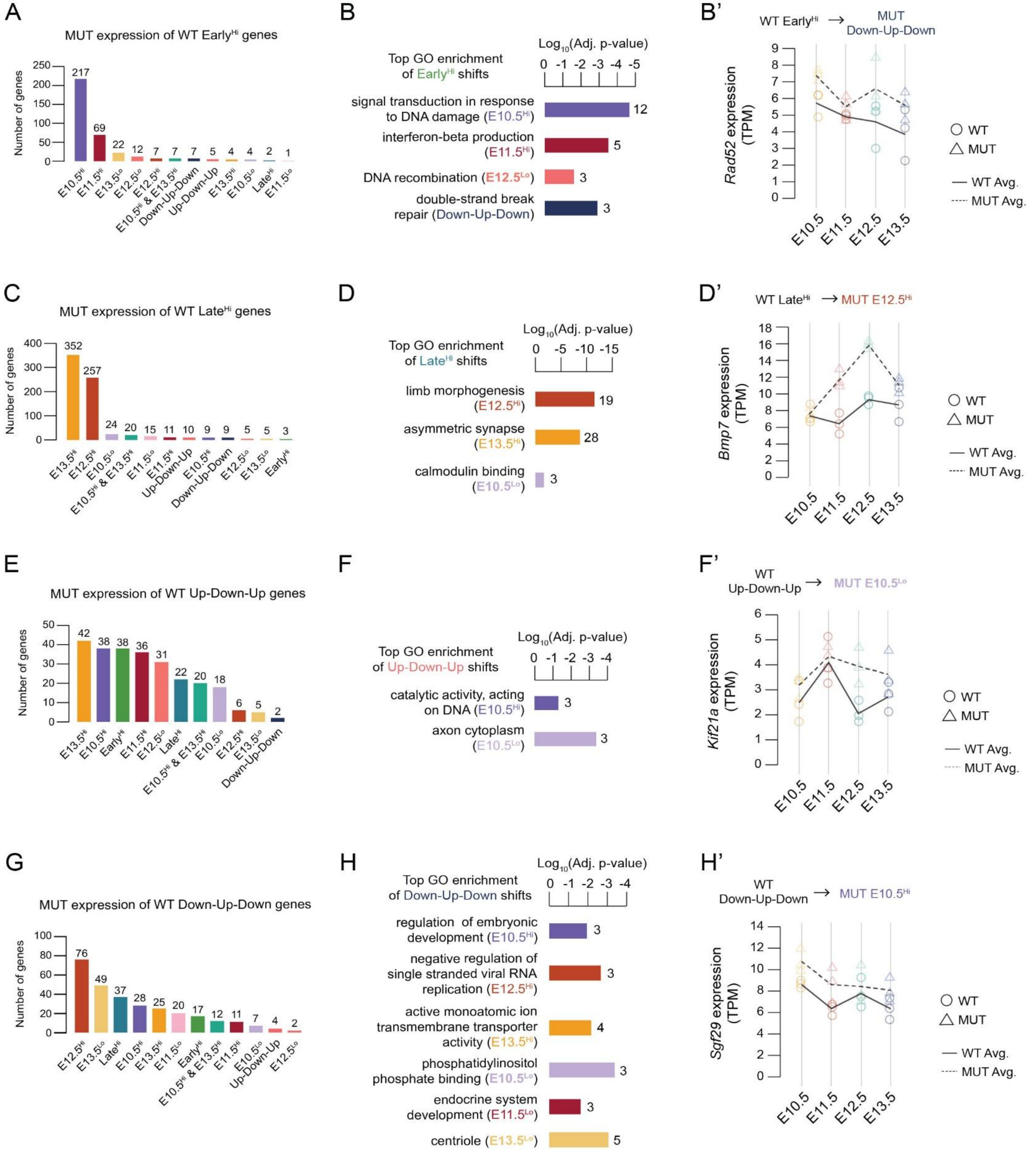
A
MUT expression of WT Early^Hi genes
Number of genes
250
200
150
100
50
0
217
69
22
12
7
7
7
5
4
4
2
1
E10.5^Hi
E11.5^Hi
E13.5^Lo
E12.5^Lo
E12.5^Hi
E10.5^Hi & E13.5^Hi
Down-Up-Down
Up-Down-Up
E13.5^Hi
E10.5^Lo
Late^Hi
E11.5^Lo
B
Top GO enrichment of Early^Hi shifts
Log10(Adj. p-value)
0 -1 -2 -3 -4 -5
signal transduction in response to DNA damage (E10.5^Hi)
12
interferon-beta production (E11.5^Hi)
5
DNA recombination (E12.5^Lo)
3
double-strand break repair (Down-Up-Down)
3
B'
WT Early^Hi → MUT Down-Up-Down
Rad52 expression (TPM)
9
8
7
6
5
4
3
2
1
0
E10.5
E11.5
E12.5
E13.5
WT
MUT
WT Avg.
MUT Avg.
C
MUT expression of WT Late^Hi genes
Number of genes
400
300
200
100
0
352
257
24
20
15
11
10
9
9
5
5
3
E13.5^Hi
E12.5^Hi
E10.5^Lo
E10.5^Hi & E13.5^Hi
E11.5^Lo
E11.5^Hi
Up-Down-Up
E10.5^Hi
Down-Up-Down
E12.5^Lo
E13.5^Lo
Early^Hi
D
Top GO enrichment of Late^Hi shifts
Log10(Adj. p-value)
0 -5 -10 -15
limb morphogenesis (E12.5^Hi)
19
asymmetric synapse (E13.5^Hi)
28
calmodulin binding (E10.5^Lo)
3
D'
WT Late^Hi → MUT E12.5^Hi
Bmp7 expression (TPM)
18
16
14
12
10
8
6
4
2
0
E10.5
E11.5
E12.5
E13.5
WT
MUT
WT Avg.
MUT Avg.
E
MUT expression of WT Up-Down-Up genes
Number of genes
50
40
30
20
10
0
42
38
38
36
31
22
20
18
6
5
2
E13.5^Hi
E10.5^Hi
Early^Hi
E11.5^Hi
E12.5^Lo
Late^Hi
E10.5^Hi & E13.5^Hi
E10.5^Lo
E12.5^Hi
E13.5^Lo
Down-Up-Down
F
Top GO enrichment of Up-Down-Up shifts
Log10(Adj. p-value)
0 -1 -2 -3 -4
catalytic activity, acting on DNA (E10.5^Hi)
3
axon cytoplasm (E10.5^Lo)
3
F'
WT Up-Down-Up → MUT E10.5^Lo
Kif21a expression (TPM)
6
5
4
3
2
1
0
E10.5
E11.5
E12.5
E13.5
WT
MUT
WT Avg.
MUT Avg.
G
MUT expression of WT Down-Up-Down genes
Number of genes
100
80
60
40
20
0
76
49
37
28
25
20
17
12
11
7
4
2
E12.5^Hi
E13.5^Lo
Late^Hi
E10.5^Hi
E13.5^Hi
E11.5^Lo
Early^Hi
E10.5^Hi & E13.5^Hi
E11.5^Hi
E10.5^Lo
Up-Down-Up
E12.5^Lo
H
Top GO enrichment of Down-Up-Down shifts
Log10(Adj. p-value)
0 -1 -2 -3 -4
regulation of embryonic development (E10.5^Hi)
3
negative regulation of single stranded viral RNA replication (E12.5^Hi)
3
active monoatomic ion transmembrane transporter activity (E13.5^Hi)
4
phosphatidylinositol phosphate binding (E10.5^Lo)
3
endocrine system development (E11.5^Lo)
3
centriole (E13.5^Lo)
5
H'
WT Down-Up-Down → MUT E10.5^Hi
Sgf29 expression (TPM)
14
12
10
8
6
4
2
0
E10.5
E11.5
E12.5
E13.5
WT
MUT
WT Avg.
MUT Avg.

**Figure S14. Redistribution of Early High, Late High, and biphasic WT-only expression trajectories following U11 loss, related to Figure 2.** (A) Distribution of mutant trajectory assignments for genes that exclusively follow the WT Early High trajectory. Bars indicate the number of genes reassigned to each mutant trajectory. (B) Top GO enrichment results for each Early High WT-to-MUT trajectory shift. The absence of a trajectory indicates that shift had no functional enrichment results. Bars represent significantly enriched GO terms (BP, CC, MF; adjusted $p < 0.05$), with bar length indicating the $\log_{10}$(adjusted p-value) and numbers denoting the number of genes underlying enrichment of each term. (B') Example expression profile for *Rad52*, underlying the GO term "double-strand break repair" in panel B. Lines show the mean expression (TPM) across time for WT (solid line, circles) and mutant (dashed line, triangles), with biological replicates overlaid. (C) Distribution of mutant trajectory assignments for genes that exclusively follow the WT Late High trajectory. (D) Top GO enrichment results for each Late High WT-to-MUT trajectory shift. (D') Example expression profile for *Bmp7*, underlying the GO term "limb morphogenesis" in panel D. (E) Distribution of mutant trajectory assignments for genes that exclusively follow the WT Up-Down-Up trajectory. (F) Top GO enrichment results for each Up-Down-Up WT-to-MUT trajectory shift. (F') Example expression profile for *Kif21a*, underlying the GO term "axon cytoplasm" in panel F. (G) Distribution of mutant trajectory assignments for genes that exclusively follow the WT Down-Up-Down trajectory. (H) Top GO enrichment results for each Down-Up-Down WT-to-MUT trajectory shift. (H') Example expression profile for *Sgf29* underlying the GO term "regulation of embryonic development" in panel H.

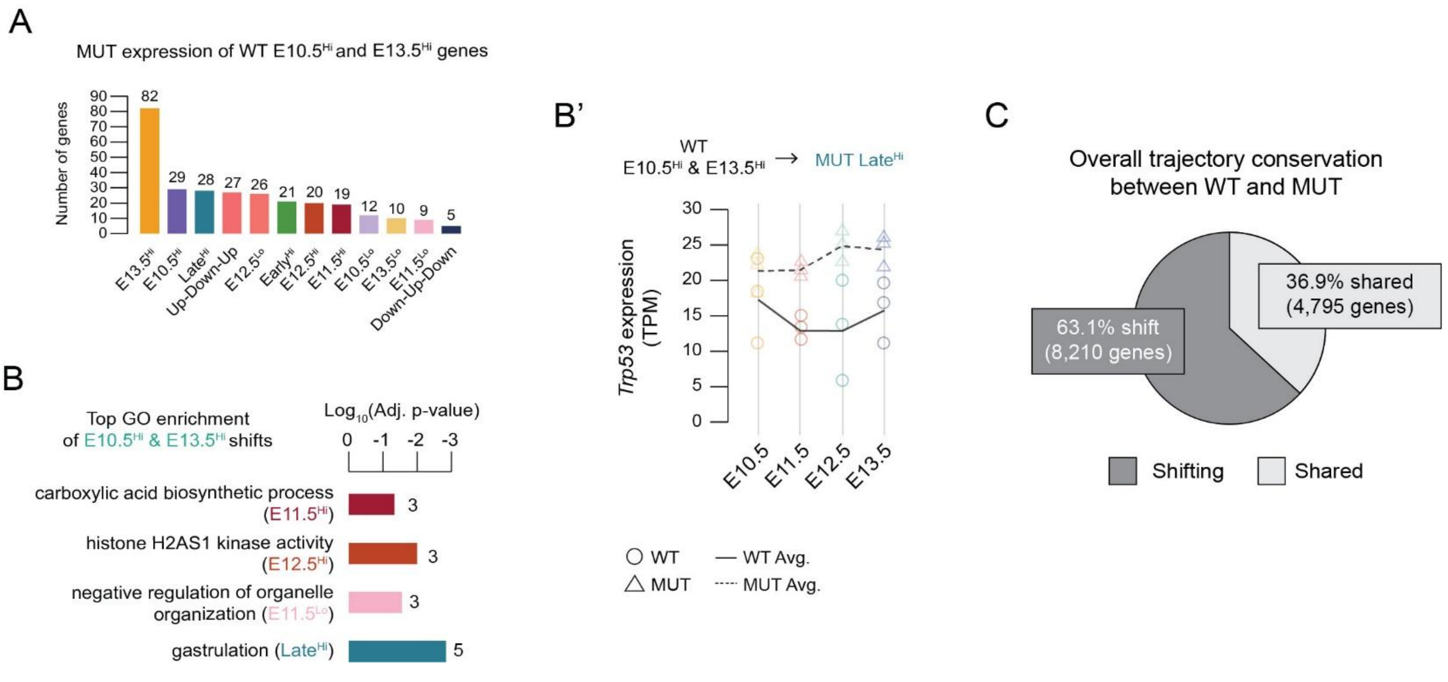


**Figure S15. Redistribution of E10.5 and E13.5 High WT-only trajectories following U11 loss, related to Figure 2.** (A) Distribution of mutant trajectory assignments for genes that exclusively follow the WT E10.5 and E13.5 High trajectory. Bars indicate the number of genes reassigned to each mutant trajectory. (B) Top GO enrichment results for each E10.5 and E13.5 High WT-to-MUT trajectory shift. The absence of a trajectory indicates that shift had no functional enrichment results. Bars represent significantly enriched GO terms (BP, CC, MF; adjusted $p < 0.05$), with bar length indicating the $log_{10}$(adjusted p-value) and numbers denoting the number of genes underlying enrichment of each term. (B') Example expression profile for *Trp53*, underlying the GO term "gastrulation". Lines show the mean expression (TPM) across time for WT (solid line, circles) and mutant (dashed line, triangles), with biological replicates overlaid. (C) Overall trajectory conservation between WT and mutant datasets. Pie chart depicts the percentage of genes sharing trajectory assignments (Shared; 36.9%, 4,795 genes) compared to those exhibiting trajectory shifts following U11 loss (Shifting; 63.1%, 8,210 genes).

Superimposing static differential expression on trajectory conservation

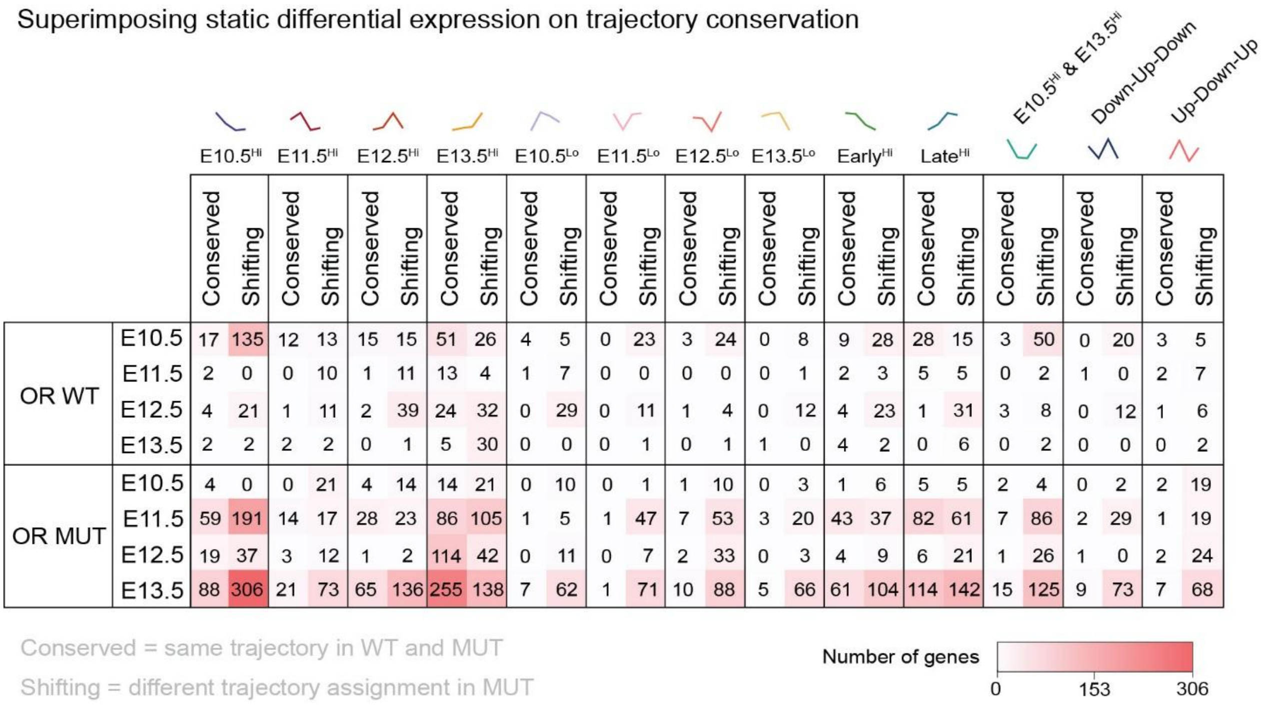


| | | $E10.5^{Hi}$ | | $E11.5^{Hi}$ | | $E12.5^{Hi}$ | | $E13.5^{Hi}$ | | $E10.5^{Lo}$ | | $E11.5^{Lo}$ | | $E12.5^{Lo}$ | | $E13.5^{Lo}$ | | $Early^{Hi}$ | | $Late^{Hi}$ | | $E10.5^{Hi}$ & $E13.5^{Hi}$ | | Down-Up-Down | | Up-Down-Up | |
|---|---|---|---|---|---|---|---|---|---|---|---|---|---|---|---|---|---|---|---|---|---|---|---|---|---|---|---|
| | | Conserved | Shifting | Conserved | Shifting | Conserved | Shifting | Conserved | Shifting | Conserved | Shifting | Conserved | Shifting | Conserved | Shifting | Conserved | Shifting | Conserved | Shifting | Conserved | Shifting | Conserved | Shifting | Conserved | Shifting | Conserved | Shifting |
| OR WT | E10.5 | 17 | 135 | 12 | 13 | 15 | 15 | 51 | 26 | 4 | 5 | 0 | 23 | 3 | 24 | 0 | 8 | 9 | 28 | 28 | 15 | 3 | 50 | 0 | 20 | 3 | 5 |
| | E11.5 | 2 | 0 | 0 | 10 | 1 | 11 | 13 | 4 | 1 | 7 | 0 | 0 | 0 | 0 | 0 | 1 | 2 | 3 | 5 | 5 | 0 | 2 | 1 | 0 | 2 | 7 |
| | E12.5 | 4 | 21 | 1 | 11 | 2 | 39 | 24 | 32 | 0 | 29 | 0 | 11 | 1 | 4 | 0 | 12 | 4 | 23 | 1 | 31 | 3 | 8 | 0 | 12 | 1 | 6 |
| | E13.5 | 2 | 2 | 2 | 2 | 0 | 1 | 5 | 30 | 0 | 0 | 0 | 1 | 0 | 1 | 1 | 0 | 4 | 2 | 0 | 6 | 0 | 2 | 0 | 0 | 0 | 2 |
| OR MUT | E10.5 | 4 | 0 | 0 | 21 | 4 | 14 | 14 | 21 | 0 | 10 | 0 | 1 | 1 | 10 | 0 | 3 | 1 | 6 | 5 | 5 | 2 | 4 | 0 | 2 | 2 | 19 |
| | E11.5 | 59 | 191 | 14 | 17 | 28 | 23 | 86 | 105 | 1 | 5 | 1 | 47 | 7 | 53 | 3 | 20 | 43 | 37 | 82 | 61 | 7 | 86 | 2 | 29 | 1 | 19 |
| | E12.5 | 19 | 37 | 3 | 12 | 1 | 2 | 114 | 42 | 0 | 11 | 0 | 7 | 2 | 33 | 0 | 3 | 4 | 9 | 6 | 21 | 1 | 26 | 1 | 0 | 2 | 24 |
| | E13.5 | 88 | 306 | 21 | 73 | 65 | 136 | 255 | 138 | 7 | 62 | 1 | 71 | 10 | 88 | 5 | 66 | 61 | 104 | 114 | 142 | 15 | 125 | 9 | 73 | 7 | 68 |

**Figure S16. Superimposing static differential gene expression and expression trajectories reveals widespread changes in magnitude independent of trajectory shifts, related to Figure 2.** Heatmap superimposing static WT-MUT differential expression onto expression trajectory behavior. Genes are grouped by their WT trajectory assignment and classified as either conserved (same trajectory assignment in WT and mutant) or shifting (different trajectory assignment in mutant). Rows indicate the developmental stages at which genes are OR in WT or mutant, and color intensity reflects the number of genes in each category. This analysis demonstrates that differences in expression magnitude occur across both conserved and shifting trajectories, consistent with prior analyses showing magnitude differences both in the absence of trajectory changes and in conjunction with trajectory shifts. See also Figures S11-S15.

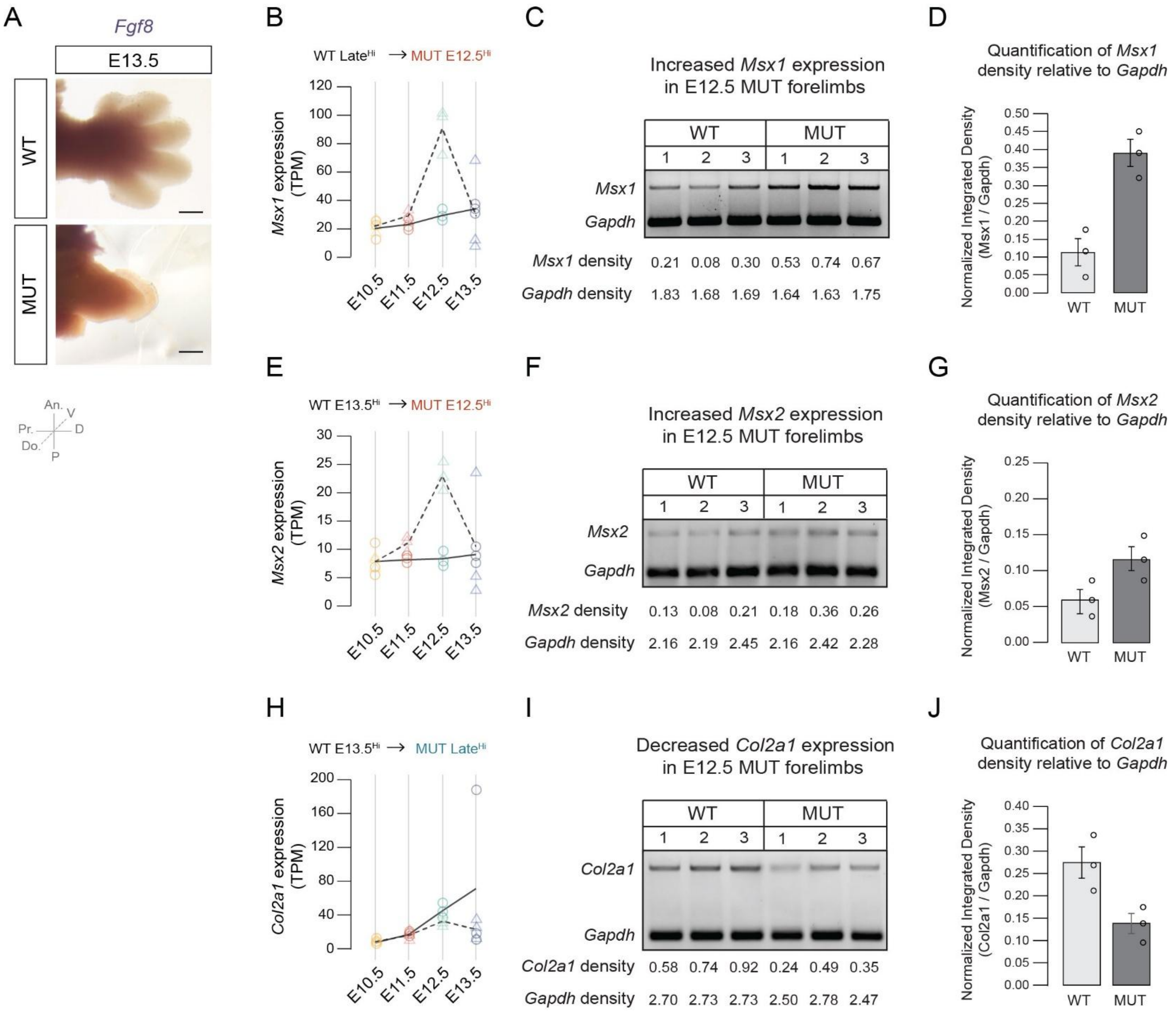
A
Fgf8
E13.5
WT
MUT
An.
V
Pr.
D
Do.
P
B
WT Late^Hi → MUT E12.5^Hi
Msx1 expression (TPM)
E10.5
E11.5
E12.5
E13.5
C
Increased Msx1 expression in E12.5 MUT forelimbs
WT
MUT
1 2 3 1 2 3
Msx1
Gapdh
Msx1 density 0.21 0.08 0.30 0.53 0.74 0.67
Gapdh density 1.83 1.68 1.69 1.64 1.63 1.75
D
Quantification of Msx1 density relative to Gapdh
Normalized Integrated Density (Msx1 / Gapdh)
WT
MUT
E
WT E13.5^Hi → MUT E12.5^Hi
Msx2 expression (TPM)
F
Increased Msx2 expression in E12.5 MUT forelimbs
Msx2
Gapdh
Msx2 density 0.13 0.08 0.21 0.18 0.36 0.26
Gapdh density 2.16 2.19 2.45 2.16 2.42 2.28
G
Quantification of Msx2 density relative to Gapdh
Normalized Integrated Density (Msx2 / Gapdh)
H
WT E13.5^Hi → MUT Late^Hi
Col2a1 expression (TPM)
I
Decreased Col2a1 expression in E12.5 MUT forelimbs
Col2a1
Gapdh
Col2a1 density 0.58 0.74 0.92 0.24 0.49 0.35
Gapdh density 2.70 2.73 2.73 2.50 2.78 2.47
J
Quantification of Col2a1 density relative to Gapdh
Normalized Integrated Density (Col2a1 / Gapdh)

**Figure S17. Validation of trajectory shifts by whole mount in situ hybridization and RT-PCR analysis, related to Figure 2.** (A) Whole mount in situ hybridization (WISH) for *Fgf8* in E13.5 WT and mutant forelimbs. Orientation schematic indicates the axes of development. (B) RNA-seq expression trajectories for *Msx1* (WT Late high → MUT E12.5 high). Lines show the mean expression (TPM) across time for WT (solid line, circles) and mutant (dashed line, triangles), with biological replicates overlaid. (C) RT-PCR analysis of *Msx1* expression in E12.5 WT and mutant forelimbs. *Msx1* (top band) and *Gapdh* (bottom band) were amplified in the same PCR reaction (duplex PCR) to control for loading variability and enable normalization of band intensity. Three biological replicates are shown per genotype. (D) Quantification of *Msx1* band intensity normalized to *Gapdh*. Bars represent mean $\pm$ SEM; points indicate biological replicates. (E) RNA-seq expression trajectories for *Msx2* (WT E13.5 high → MUT E12.5 high). Lines show the mean expression (TPM) across time for WT (solid line, circles) and mutant (dashed line, triangles), with biological replicates overlaid. (F) RT-PCR analysis of *Msx2* expression in E12.5 WT and mutant forelimbs. Duplex PCR results for *Msx2* (top band) and *Gapdh* (bottom band) with three biological replicates shown per genotype. (G) Quantification of *Msx2* band intensity normalized to *Gapdh*. Bars represent mean $\pm$ SEM; points indicate biological replicates. (H) RNA-seq expression trajectories for *Col2a1* (WT E13.5 high → MUT Late high). Lines show the mean expression (TPM) across time for WT (solid line, circles) and mutant (dashed line, triangles), with biological replicates overlaid. (I) RT-PCR analysis of *Col2a1* expression in E12.5 WT and mutant forelimbs. Duplex PCR results for *Col2a1* (top band) and *Gapdh* (bottom band) with three biological replicates shown per genotype. (J) Quantification of *Col2a1* band intensity normalized to *Gapdh*. Bars represent mean $\pm$ SEM; points indicate biological replicates. An.; anterior, P; posterior, D; distal, Pr.; proximal, Do.; dorsal, V; ventral. Scale bars = 0.5 mm. See also Table S5.

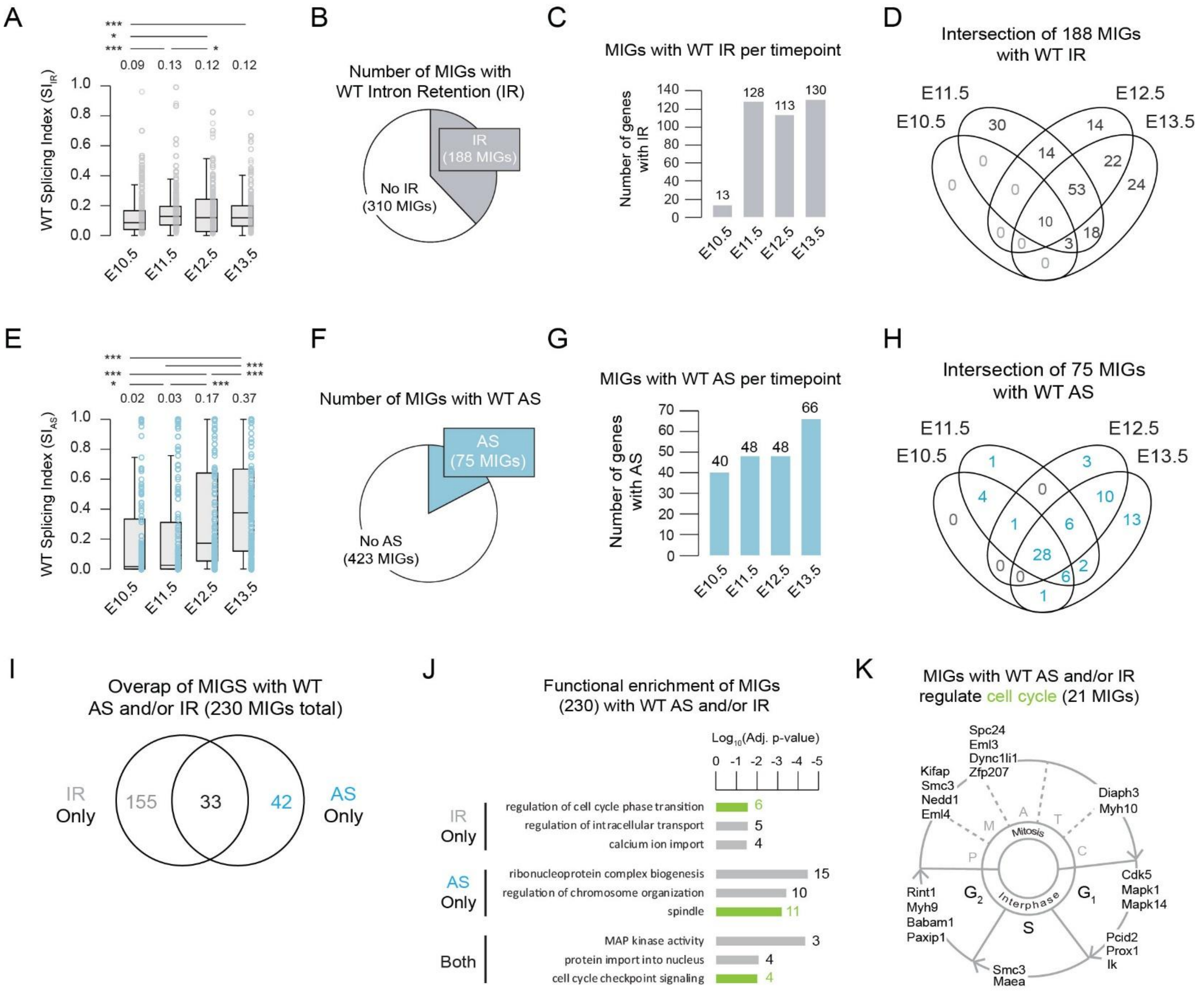


**Figure S18. MIG splicing is dynamically regulated across developmental time in the WT forelimb, related to Figure 3.** (A) Distribution of intron retention splicing index ($SI_{IR}$) values for minor intron-containing genes (MIGs) across WT forelimb development (E10.5-E13.5). Boxplots summarize the $SI_{IR}$ values for all expressed MIGs that pass filtering in all timepoints, with individual introns shown as points. Median $SI_{IR}$ values for each timepoint are indicated above each boxplot. Boxes represent the interquartile range (25$^{th}$ to 75$^{th}$ percentile), center lines indicate the median, and whiskers denote 1.5x the interquartile range. Statistical significance across developmental timepoints was assessed using a Friedman test, treating each intron as a repeated measure across time. Post hoc pairwise Wilcoxon signed-rank tests were performed between timepoints with Benjamini Hochberg correction. (B) Proportion of MIGs exhibiting detectable intron retention (IR) in WT forelimbs. (C) Number of MIGs with WT intron retention per timepoint. (D) Intersection of MIGs exhibiting intron retention across timepoints, illustrating both temporally restricted and common IR events across time. (E) Distribution of alternative splicing, splicing index ($SI_{AS}$) values for MIGs across WT forelimb development (E10.5-E13.5). Boxplots summarize the $SI_{AS}$ values for all expressed MIGs that pass filtering in all timepoints, with individual introns shown as points. Median $SI_{AS}$ values for each timepoint are indicated above each boxplot. Statistical significance was determined as previously described for panel A. (F) Proportion of MIGs exhibiting detectable alternative splicing (AS) in WT forelimbs. (G) Number of MIGs with WT alternative splicing per timepoint. (H) Intersection of MIGs exhibiting alternative splicing across timepoints, illustrating both temporally restricted and common AS events across time. (I) Overlap between MIGs with AS and/or IR. (J) GO enrichment of MIGs exhibiting WT AS and/or IR. (K) Diagram highlighting MIGs with WT AS and/or IR that regulate cell-cycle associated processes.

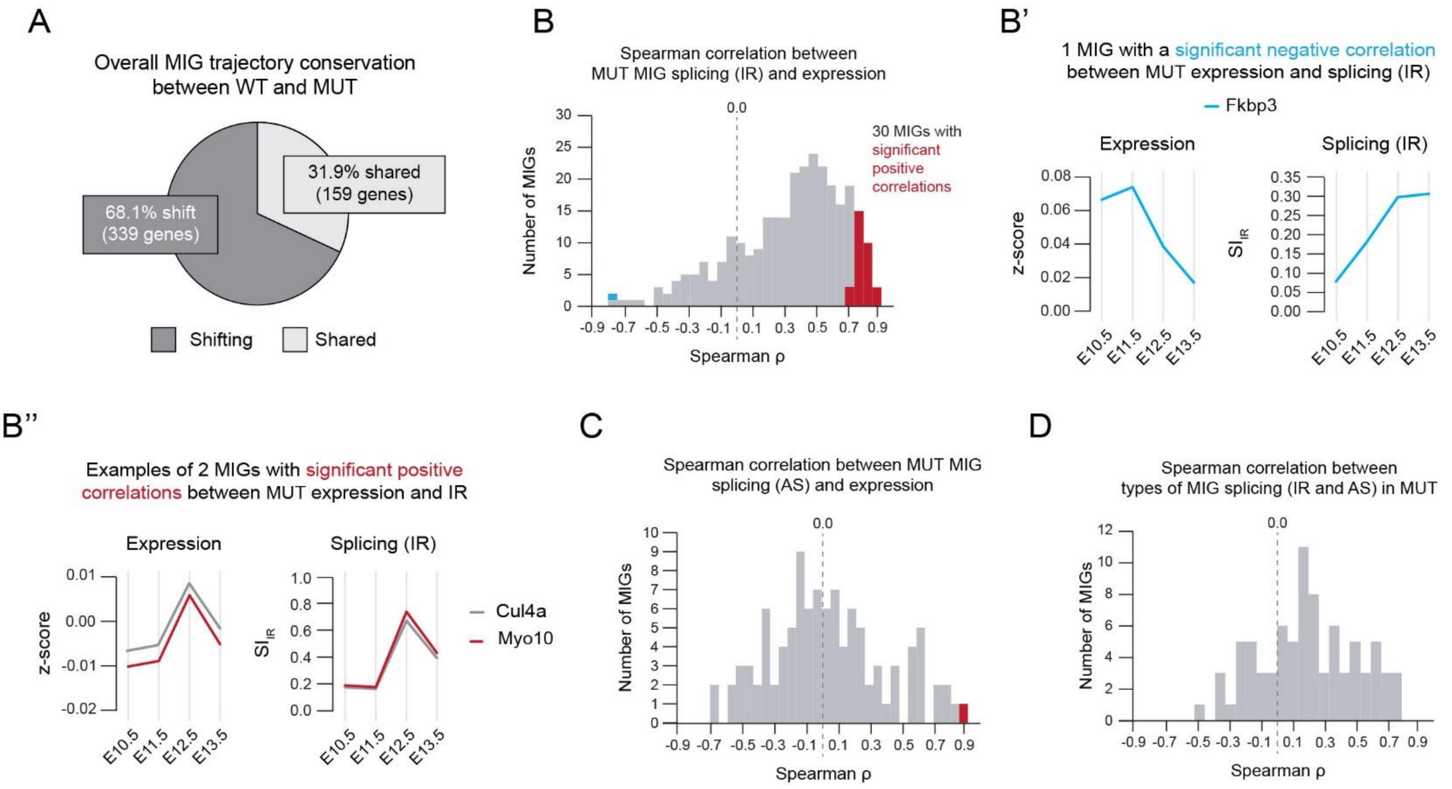


**Figure S19. Altered MIG splicing upon U11 loss does not globally correlate with shifts in MIG expression, related to Figure 3.** (A) Overall conservation of MIG expression trajectories between WT and mutant forelimbs. Pie chart shows the percentage of MIGs that retain the same trajectory (shared) or are reassigned (shifting) between genotypes. (B) Distribution of Spearman correlation coefficients (ρ) between mutant MIG expression and intron retention (IR) across developmental time (E10.5-E13.5). Significant positive and negative correlations (Benjamini-Hochberg adjusted $p < 0.05$) are highlighted in red and blue, respectively. (B') Z-scored expression and splicing index ($SI_{IR}$) values for the single MIG, *Fkbp3*, exhibiting a significant negative correlation between expression and IR. (B'') Examples of z-scored expression and $SI_{IR}$ values for two MIGs, *Cul4a and Myo10*, exhibiting a significant positive correlation between expression and IR. (C) Distribution of Spearman correlation coefficients (ρ) between mutant MIG expression and alternative splicing (AS; $SI_{AS}$) across developmental time. Significant positive correlations (Benjamini-Hochberg adjusted $p < 0.05$) are highlighted in red. (D) Distribution of Spearman correlation coefficients (ρ) between mutant MIG IR and AS across developmental time.

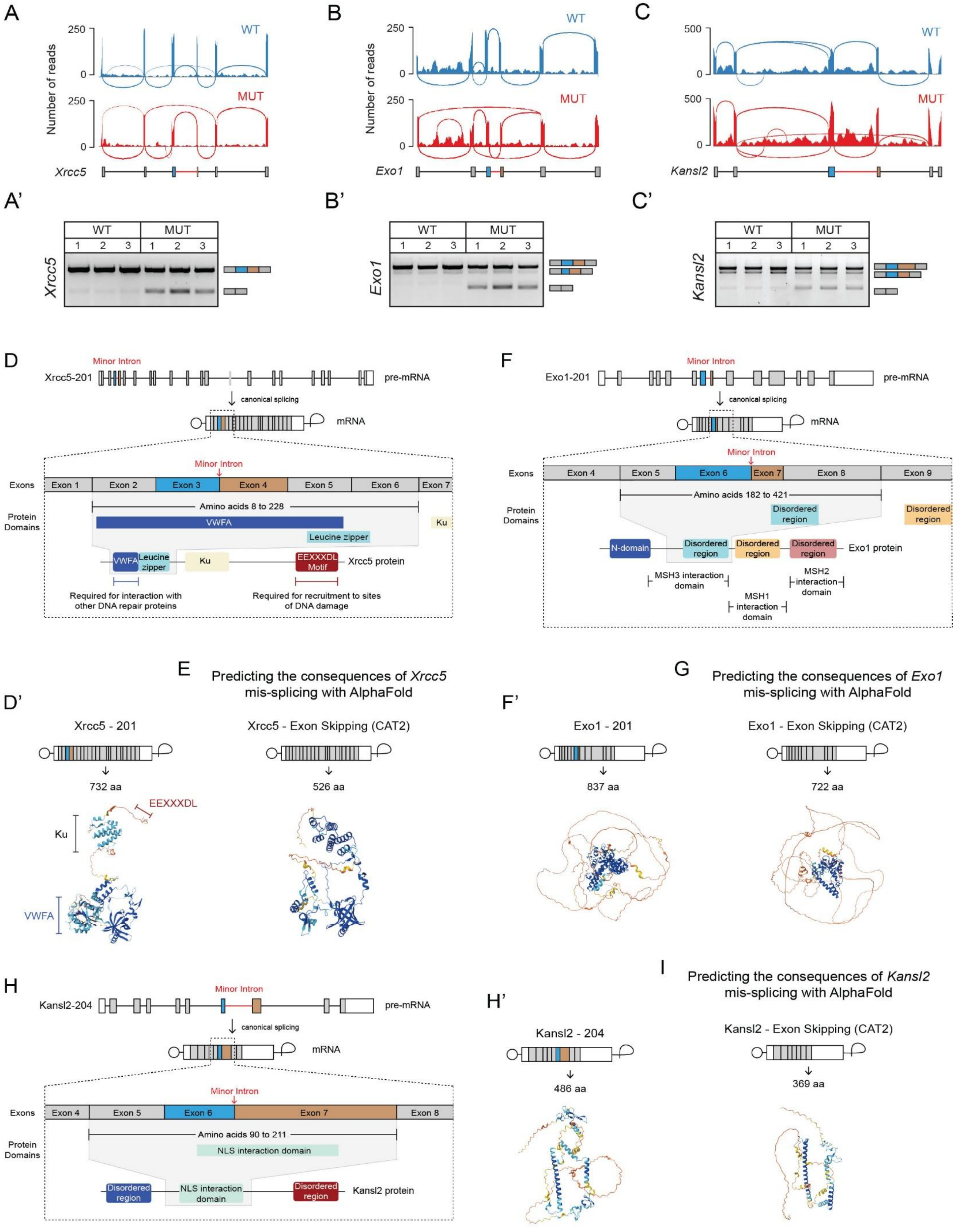
A
B
C
WT
MUT
Number of reads
Xrcc5
Exo1
Kansl2
A'
B'
C'
D
Minor Intron
Xrcc5-201
pre-mRNA
canonical splicing
mRNA
Exons
Exon 1
Exon 2
Exon 3
Exon 4
Exon 5
Exon 6
Exon 7
Protein Domains
Amino acids 8 to 228
VWFA
Leucine zipper
Ku
EEXXXDL Motif
Xrcc5 protein
Required for interaction with other DNA repair proteins
Required for recruitment to sites of DNA damage
F
Exo1-201
Exon 8
Exon 9
Amino acids 182 to 421
Disordered region
N-domain
Exo1 protein
MSH3 interaction domain
MSH2 interaction domain
MSH1 interaction domain
E
Predicting the consequences of Xrcc5 mis-splicing with AlphaFold
D'
Xrcc5 - 201
Xrcc5 - Exon Skipping (CAT2)
732 aa
526 aa
EEXXXDL
G
Predicting the consequences of Exo1 mis-splicing with AlphaFold
F'
Exo1 - 201
Exo1 - Exon Skipping (CAT2)
837 aa
722 aa
H
Kansl2-204
Amino acids 90 to 211
NLS interaction domain
Kansl2 protein
I
Predicting the consequences of Kansl2 mis-splicing with AlphaFold
H'
Kansl2 - 204
Kansl2 - Exon Skipping (CAT2)
486 aa
369 aa

**Figure S20. Predicted consequences of minor intron mis-splicing in DNA repair and chromatin-associated MIGs, related to Figure 3.** (A-C) Sashimi plots showing RNA-seq read coverage and alternative splicing in mutant forelimbs relative to WT for *Xrcc5* (A), *Exo1* (B), and *Kansl2* (C). Gene schematics for the region of interest are shown below each track. (A'-C') RT-PCR validation of alternative splicing events observed in panels A-C for *Xrcc5* (A, A'), *Exo1* (B, B'), and *Kansl2* (C, C') in WT and mutant forelimbs (n = 3 biological replicates per genotype). Schematics to the right of each gel image indicate canonical and alternative splicing events. (D) Transcript and protein domain organization of *Xrcc5* show that the minor intron resides between exons 3 and 4. The four exons surrounding the minor intron encode the VWFA domain, which is critical for interaction with other DNA repair proteins. The canonical isoform of *Xrcc5* (*Xrcc5-201*) encodes a 732 amino acid (aa) protein containing VWFA domain, Ku domain, EEXXXDL motif, and several structural domains (D'). (E) AlphaFold structural prediction of the 526 aa protein predicted to result from exon skipping (CAT2) in *Xrcc5*. The predicted model suggests potential loss of VWFA domain structure. (F) Transcript and protein domain organization of *Exo1* show that the minor intron resides between exons 6 and 7. The four exons surrounding the minor intron encode a disordered region and overlap with a known MSH3 interaction site. The canonical isoform of *Exo1* (*Exo1-201*) encodes an 837 aa protein containing an N-terminal domain, three disordered regions, and sites for interaction with Msh1/2/3 (F'). (G) AlphaFold structural prediction of the 722 aa protein predicted to result from exon skipping (CAT2) in *Exo1*. The predicted model suggests potential disruption of the region required for Msh3 interaction. (H) Transcript and protein domain organization of *Kansl2* show that the minor intron resides between exons 6 and 7. The four exons surrounding the minor intron encode a region required for interaction with other members of the NSL complex. The canonical isoform of *Kansl2* (*Kansl2-204*) encodes a 486 aa protein containing two disordered regions and the region required for NSL complex interaction (H'). (I) AlphaFold structural prediction of the 369 aa protein predicted to result from exon skipping (CAT2) in *Kansl2*. The predicted model suggests truncation of structural regions and potential disruption of the region required for NSL complex interaction. See also Table S5.

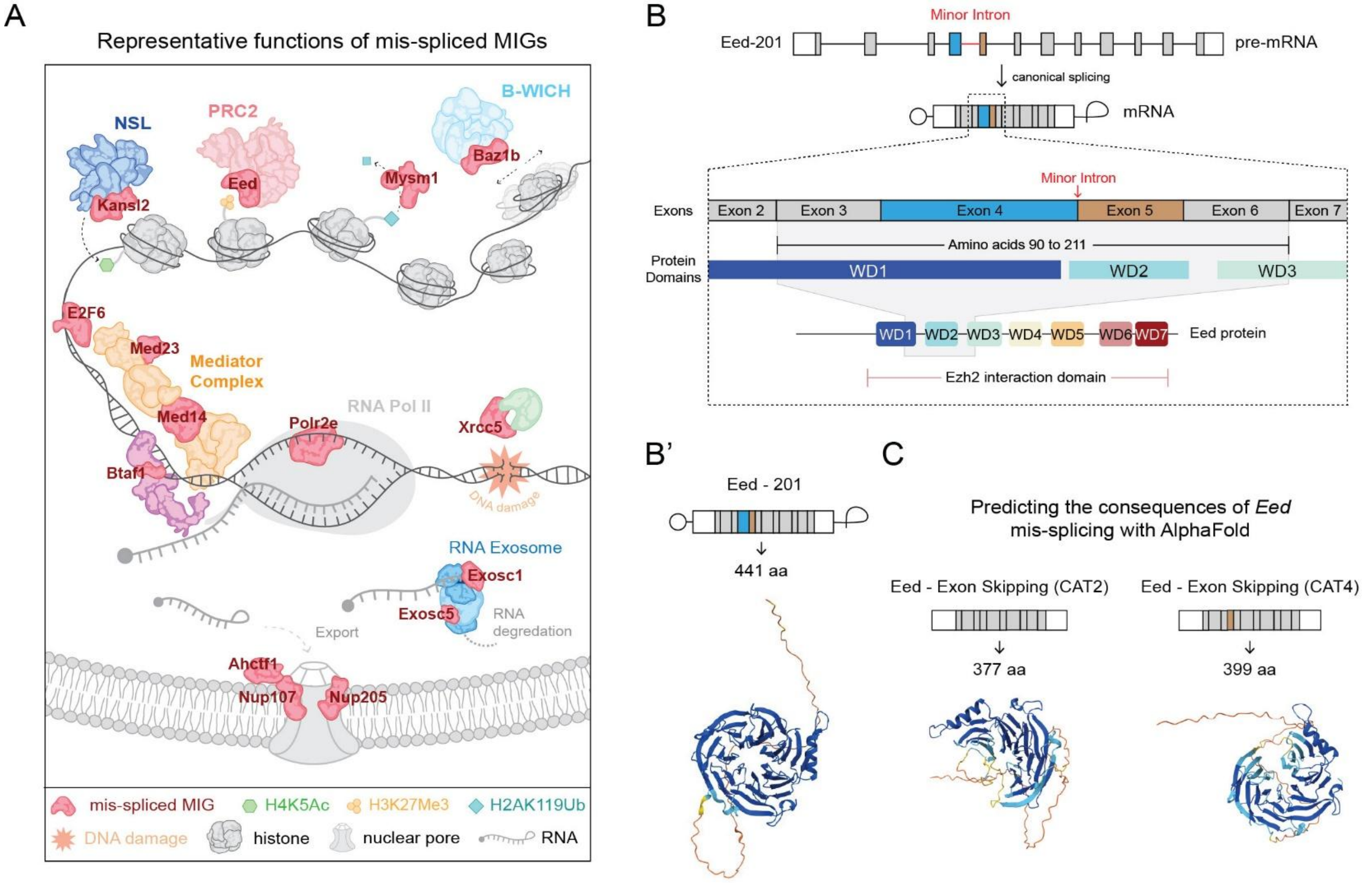


**Figure S21. Minor intron mis-splicing converges on critical MIG nodes within diverse regulatory networks, including epigenetic modifiers such as Eed, related to Figure 3.** (A) Schematic representation of the diverse regulatory pathways in which mis-spliced MIGs (red) reside, including multiple chromatin-associated and transcriptional regulatory complexes such as NSL (*Kansl2*), PRC2 (*Eed*), B-WICH complex (*Baz1b*), Mediator complex (*Med14*, *Med23*), and RNA polymerase II associated factors (*Polr2e, E2F6, Btaf1*). Additional MIGs involved in histone modification (*Mysm1*), RNA degradation (*Exosc1*, *Exosc5*), nuclear transport (*Ahctf1*, *Nup107*, *Nup205*), and DNA repair (*Xrcc5*) are also indicated. Histone modifications (H4K5Ac, H3K27Me3, H2AK119Ub) are shown to illustrate the connection between mis-spliced MIGs and epigenetic regulation. This schematic represents only a subset of the mis-spliced MIGs identified in this study and their associated functions. (B) Transcript and protein domain organization of *Eed* showing that the minor intron resides between exons 4 and 5. The four exons surrounding the minor intron encode three WD40 repeat domains, including WD1, which is essential for Eed interaction with the of PRC2 complex. The canonical isoform (*Eed-201*) encodes a 441 aa protein containing seven WD40 repeat domains that facilitate Eed's interaction with Ezh2 (B'). (C) AlphaFold structural prediction of the 377 aa and 399 aa proteins predicted to result from exon skipping (CAT2 and CAT4) in *Eed*. The predicted model suggests potential disruption of WD40 repeat architecture, including the WD1 domain critical for PRC2 interaction.

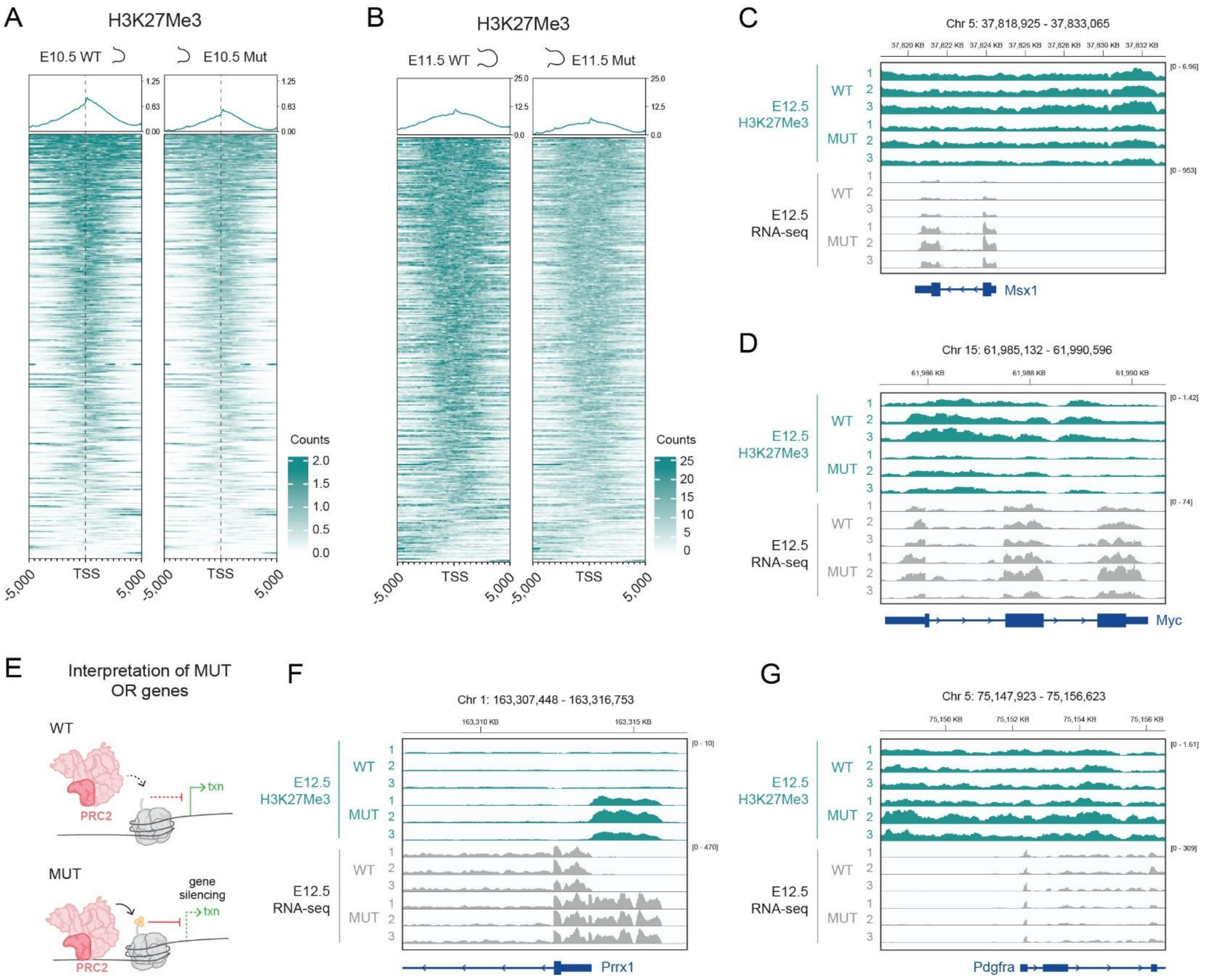


**Figure S22. Extended analysis of H3K27Me3 dynamics across developmental time in U11 mutant forelimbs, related to Figure 4.** (A-B) Heatmaps showing H3K27Me3 signal intensity centered on transcription start sites (TSS ±5 kb) for the top 10,000 WT H3K27Me3- enriched sites at E10.5 (A) and E11.5 (B) in WT and mutant forelimbs. The same WT-ranked sites depicted in Figure 4C are displayed for both genotypes and timepoints. Average H3K27Me3 signal profiles are shown above each heatmap. (C-D) IGV tracks illustrating the biological replicate data for the representative loci, *Msx1* (C) and *Myc* (D), shown in Figure 4E-F. (E) Schematic illustrating the interpretation of genes overrepresented in mutant relative to WT. (F-G) Additional representative loci illustrating increased H3K27Me3 deposition and expression changes at E12.5, including *Prrx1* (F) and *Pdgfra* (G). Tracks display H3K27Me3 CUT&RUN signal (teal) and RNA-seq coverage (grey) for WT and mutant forelimbs.

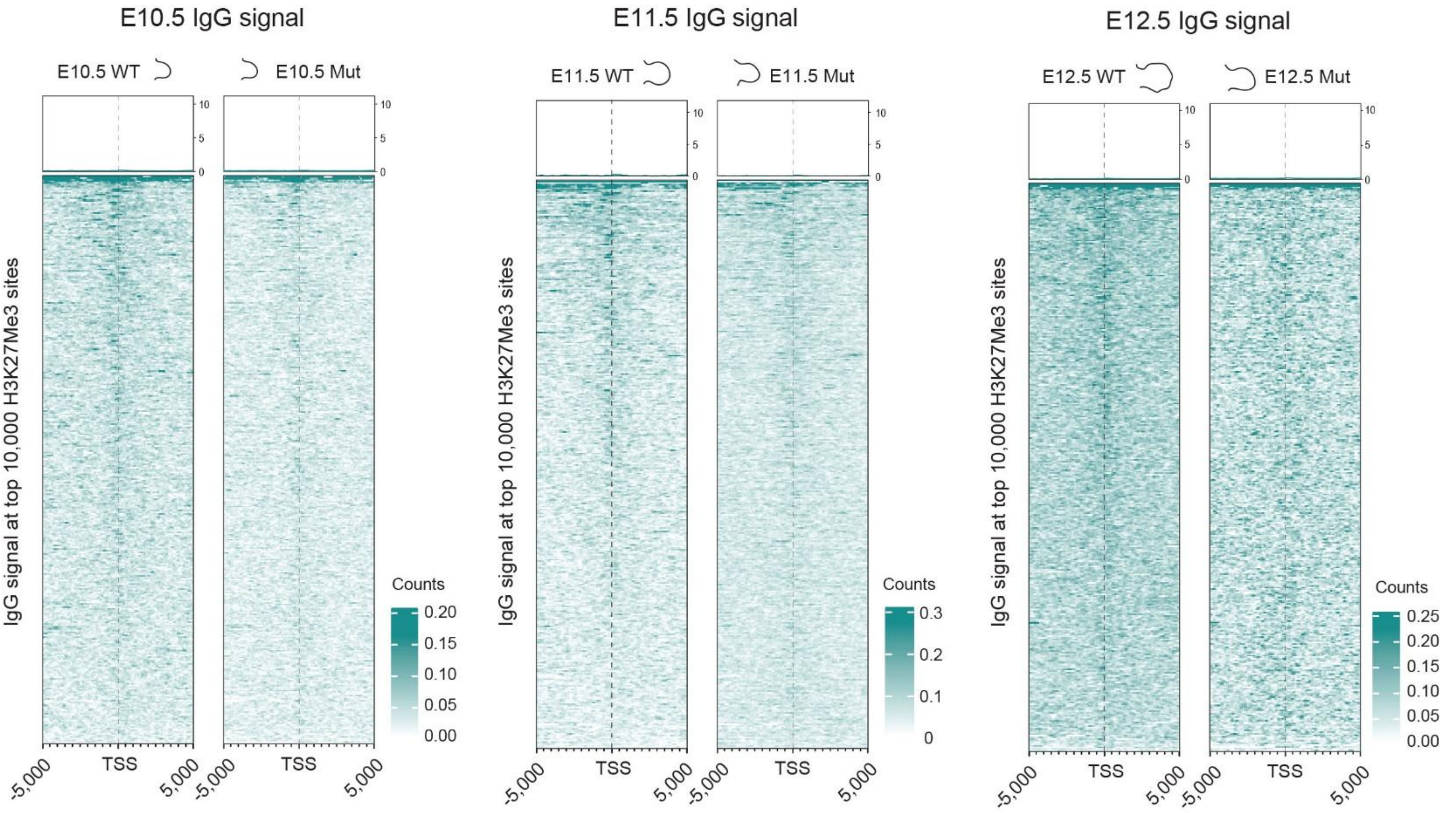


**Figure S23. IgG control profiles for H3K27Me3 CUT&RUN experiments, related to Figure 4.** Heatmaps showing IgG control signal centered around transcription start sites (TSS +5 kb) for the same top 10,000 H3K27Me3-enriched sites analyzed in Figure 4C and Figure S22A-B at E10.5, E11.5, and E12.5 for WT and mutant. Average signal is shown above each heatmap. Minimal enrichment and the absence of a strong TSS-centered signal in IgG confirms the specificity of H3K27Me3 CUT&RUN signal across genotypes and timepoints.

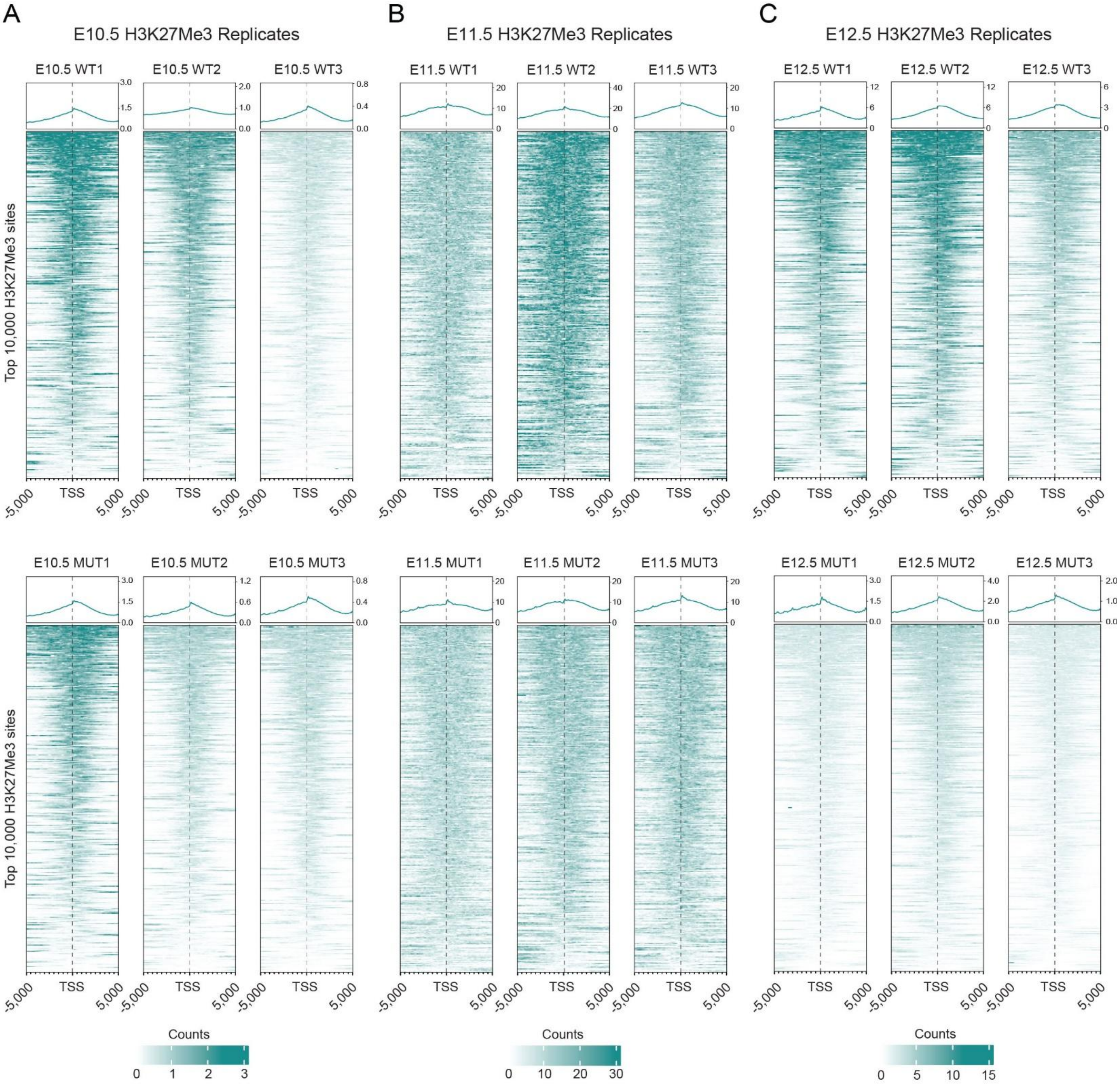


**Figure S24. Replicate concordance and H3K27Me3 signal profiles across developmental timepoints, related to Figure 4.** (A-C) Heatmaps showing H3K27Me3 signal centered around transcription start sites (TSS +5 kb) for the same top 10,000 H3K27Me3-enriched sites analyzed in Figure 4C and Figure S22A-B at E10.5 (A), E11.5 (B), and E12.5 (C) for WT and mutant. Individual biological replicates for WT (top row) and mutant (bottom row) are shown separately and average signal profiles for each replicate are shown above the corresponding heatmap. Consistent TSS-centered enrichment within each genotype and timepoint demonstrates reproducibility of H3K27Me3 across biological replicates.

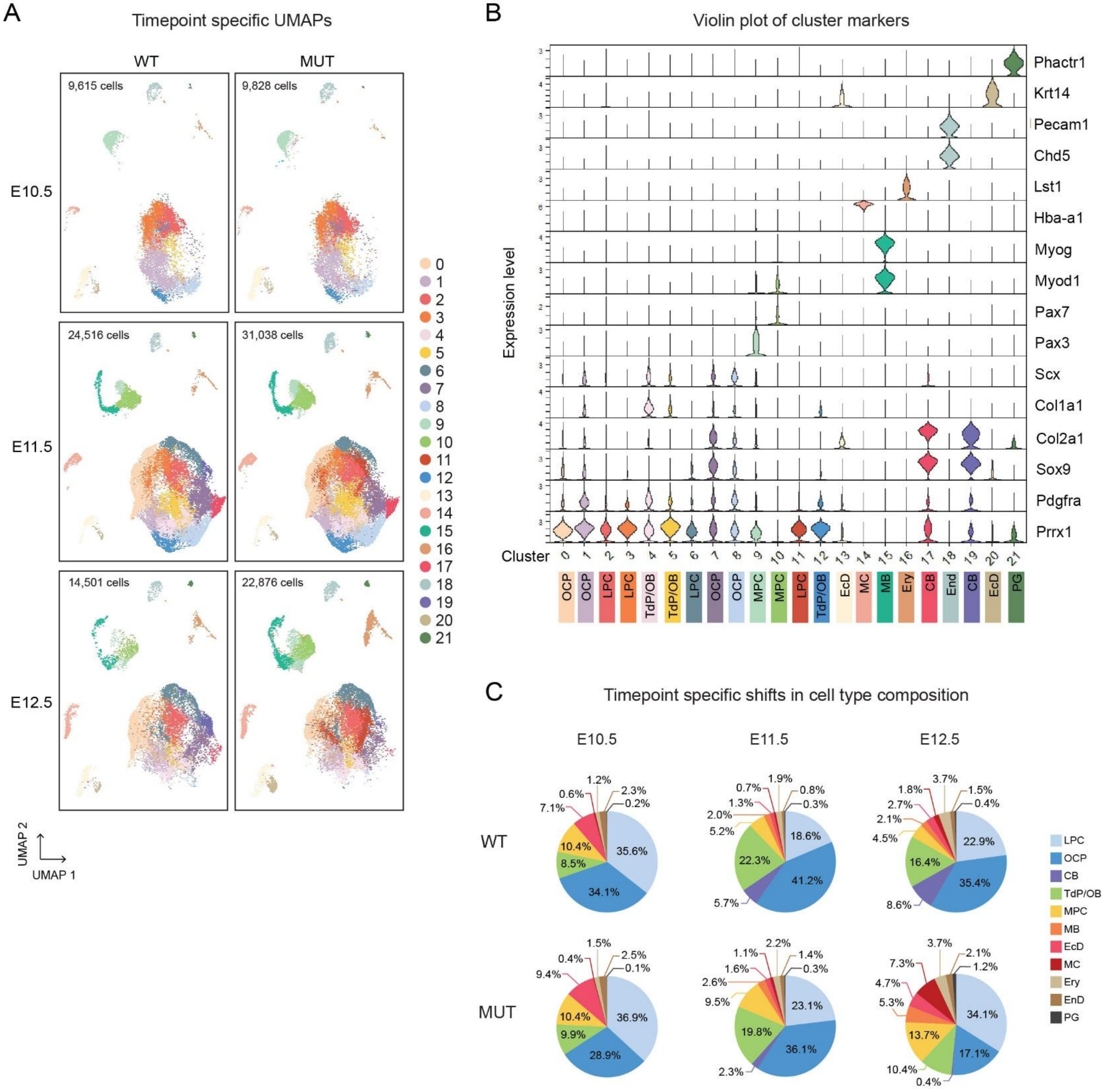


**Figure S25. Single-cell cluster annotation across developmental timepoints, related to Figure 6.** (A) Timepoint-specific UMAP visualization of integrated WT and mutant single cell RNA sequencing datasets. Cells are colored by cluster identity. The number of cells captured per genotype and timepoint is indicated in each panel. (B) Violin plots showing expression of marker genes used to annotate clusters. Maker expression supports identification of major cell types including limb progenitor cells (LPC), osteochondroprogenitor cells (OCP), chondroblasts (CB), tendon progenitor cell/osteoblast progenitors (Tdp/OB), muscle progenitor cells (MPC), myoblasts (MB), ectoderm (EcD), myeloid cells (MC), erythroid cells (Ery), endothelial cells (EnD), and peripheral glia (PG). (C) Timepoint specific cell type composition. Pie charts display proportional representation of major cell types at E10.5, E11.5, and E12.5 for each genotype. See also Data S15.

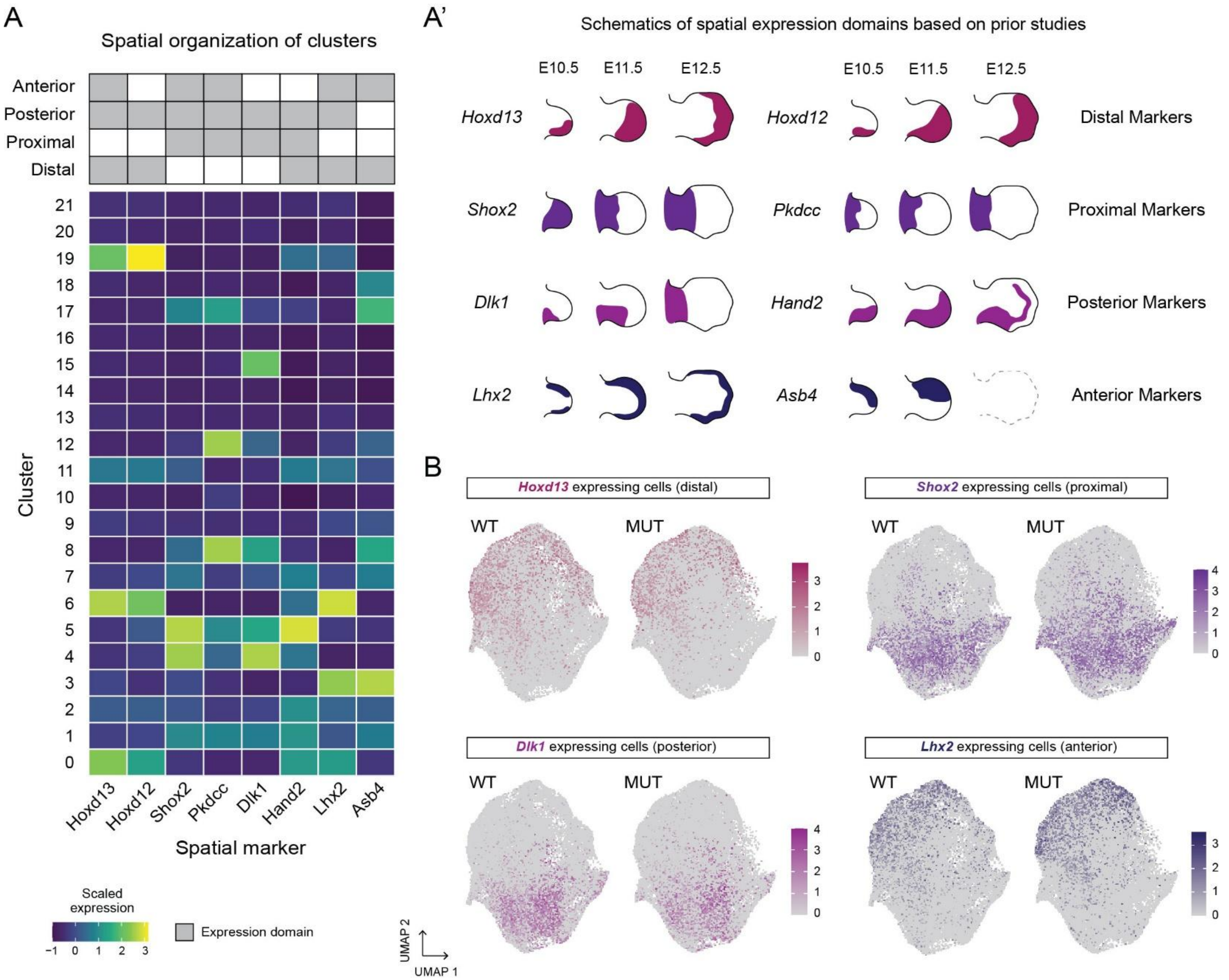


**Figure S26. Spatial marker-based organization of single-cell clusters, related to Figure 6.** (A) Heatmap showing scaled expression of established spatial markers across single-cell clusters. Markers associated with distal (*Hoxd13*, *Hoxd12*), proximal (*Shox2*, *Pkdcc*), posterior (*Dlk1*, *Hand2*), and anterior (*Lhx2*, *Asb4*) limb domains are shown. (A') Schematics illustrating spatial expression domains for each marker from E10.5-E12.5 based on prior studies[4–17]. (B) Feature plots showing expression of representative distal (*Hoxd13*), proximal (*Shox2*), posterior (*Dlk1*), and anterior (*Lhx2*) markers projected into integrated UMAP space. Spatially restricted marker expression supports subdivision of related cell types into distinct clusters based on differential anterior-posterior and proximal-distal identity. See also Data S16.

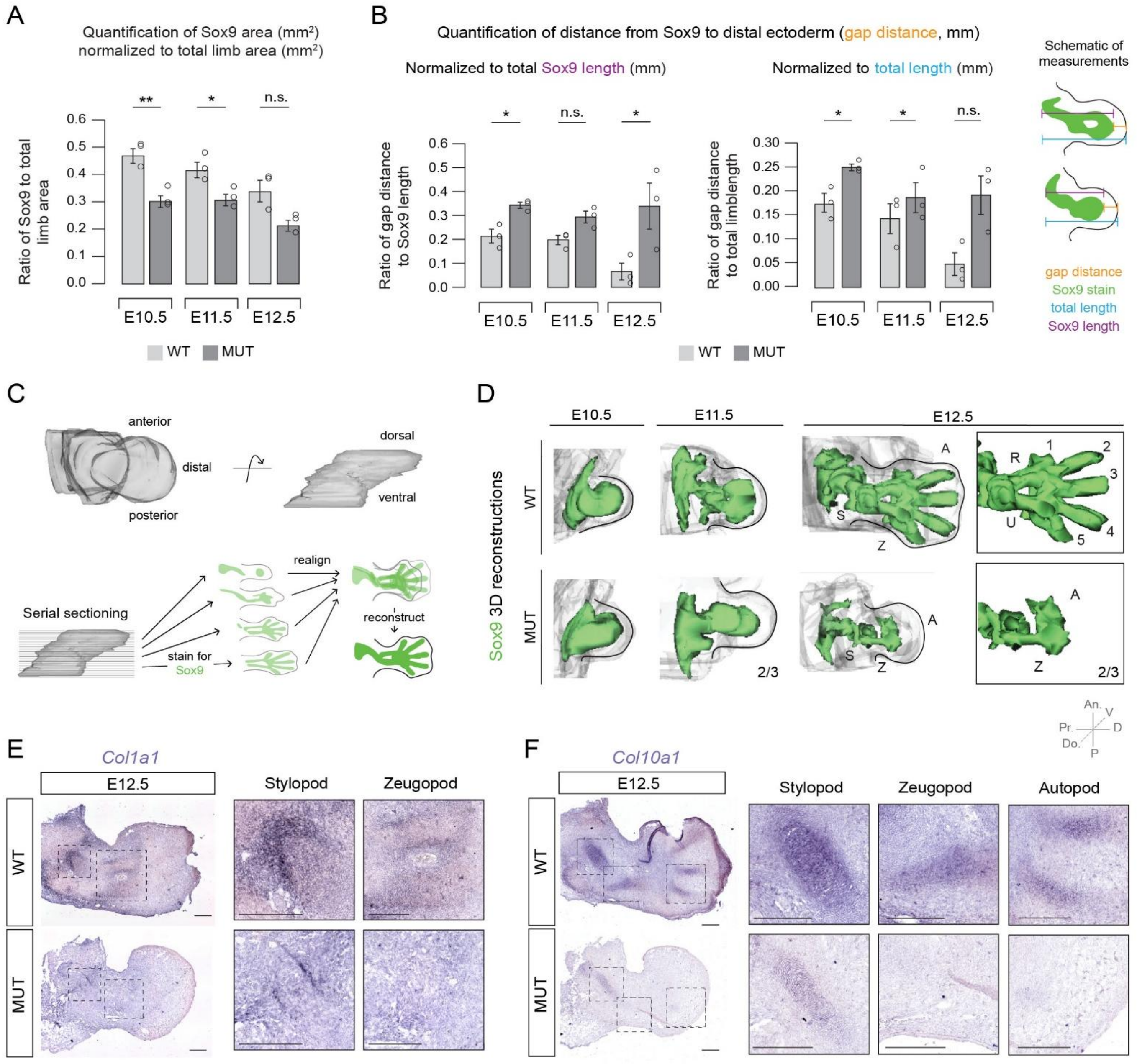


**Figure S27. Histological validation of chondrogenesis defects in mutant forelimbs, related to Figure 6.** (A) Quantification of Sox9-positive area ($mm^2$) normalized to total limb area ($mm^2$) at E10.5 – E12.5 in WT and mutant forelimbs. Bars represent mean $\pm$ SEM. Statistical significance was determined using two-tailed Welch's t-tests (n=3 biological replicates per genotype and timepoint). U11 mutant limbs exhibit a reduction in Sox9-positive tissue at early timepoints, consistent with impaired chondrogenesis. (B) Quantification of the distance between the distal ectoderm and the distal boundary of the Sox9 domain ("gap distance"). Left, gap distance normalized to total Sox9 domain length. Right, gap distance normalized to total limb length. Increased normalized gap distance in mutant limbs indicates delay or failure to initiate distal chondrogenic condensation. Schematic illustrates measurement strategy. (C) Experimental workflow for 3D reconstruction of Sox9 expression from serial sections. (D) 3D reconstructions of Sox9 domains in WT and mutant forelimbs at E10.5, E11.5, and E12.5. Mutant limbs display a reduction in distal chondrogenic condensations (2/3) along with the absence of a zeugopodial element (2/3). (E-F) In situ hybridization for *Col1a1* (E) and *Col10a1* (F) at E12.5 in WT and mutant forelimbs. Insets show higher-magnification views of stylopod, zeugopod, and autopod regions. S; stylopod, Z; zeugopod, A; autopod, R; radius, U; ulna, An.; anterior, P; posterior, D; distal, Pr.; proximal, Do.; dorsal, V; ventral. Scale bars, 200 µm. See also Table S4.

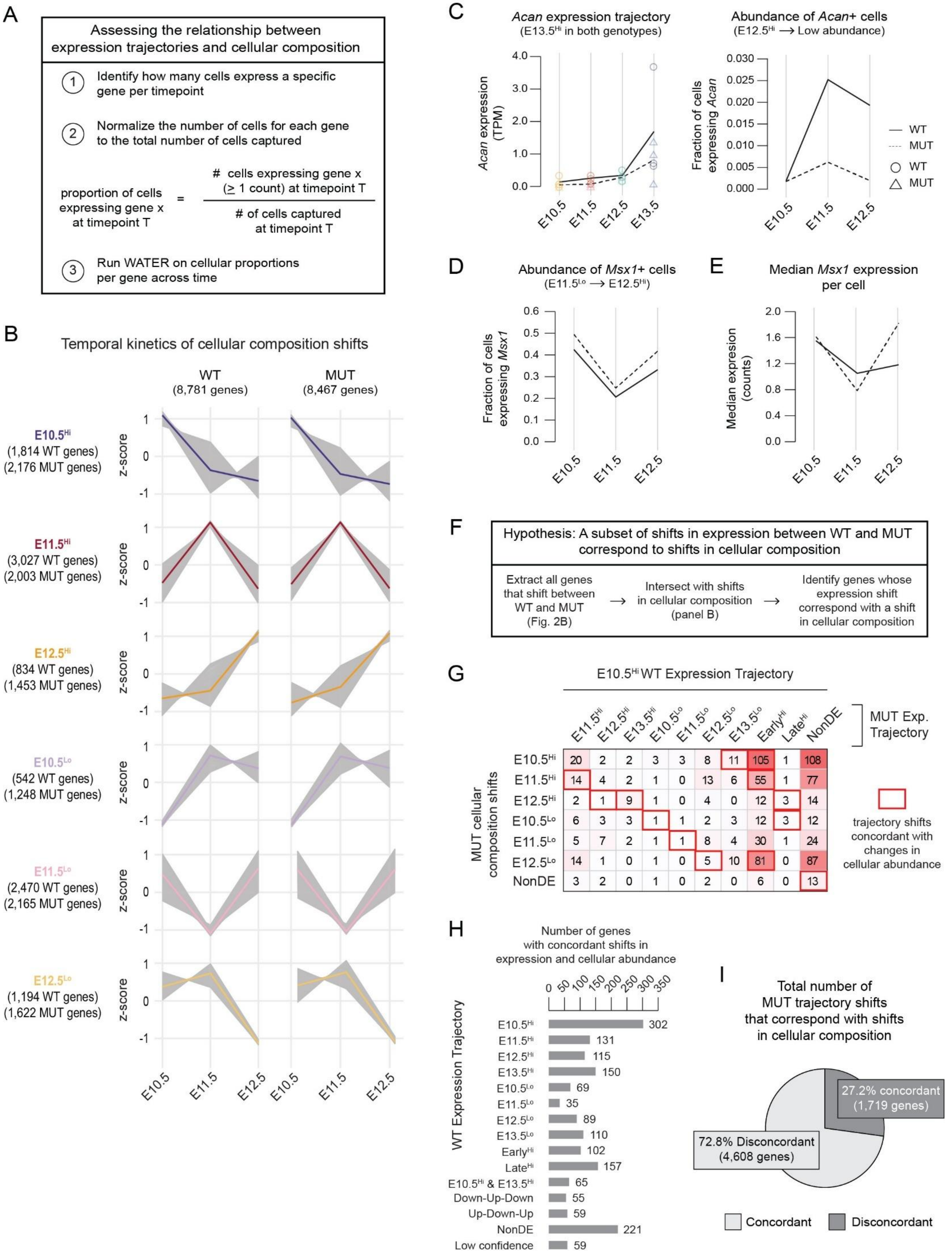
A
Assessing the relationship between expression trajectories and cellular composition
1 Identify how many cells express a specific gene per timepoint
2 Normalize the number of cells for each gene to the total number of cells captured
proportion of cells expressing gene x at timepoint T = # cells expressing gene x (≥ 1 count) at timepoint T / # of cells captured at timepoint T
3 Run WATER on cellular proportions per gene across time
B
Temporal kinetics of cellular composition shifts
WT (8,781 genes)
MUT (8,467 genes)
E10.5Hi (1,814 WT genes) (2,176 MUT genes)
E11.5Hi (3,027 WT genes) (2,003 MUT genes)
E12.5Hi (834 WT genes) (1,453 MUT genes)
E10.5Lo (542 WT genes) (1,248 MUT genes)
E11.5Lo (2,470 WT genes) (2,165 MUT genes)
E12.5Lo (1,194 WT genes) (1,622 MUT genes)
z-score
E10.5
E11.5
E12.5
C
Acan expression trajectory (E13.5Hi in both genotypes)
Acan expression (TPM)
Abundance of Acan+ cells (E12.5Hi ⟶ Low abundance)
Fraction of cells expressing Acan
WT
MUT
WT
MUT
E10.5
E11.5
E12.5
E13.5
D
Abundance of Msx1+ cells (E11.5Lo ⟶ E12.5Hi)
Fraction of cells expressing Msx1
E
Median Msx1 expression per cell
Median expression (counts)
F
Hypothesis: A subset of shifts in expression between WT and MUT correspond to shifts in cellular composition
Extract all genes that shift between WT and MUT (Fig. 2B) ⟶ Intersect with shifts in cellular composition (panel B) ⟶ Identify genes whose expression shift correspond with a shift in cellular composition
G
E10.5Hi WT Expression Trajectory
MUT Exp. Trajectory
MUT cellular composition shifts
E11.5Hi E12.5Hi E13.5Hi E10.5Lo E11.5Lo E12.5Lo E13.5Lo EarlyHi LateHi NonDE
E10.5Hi 20 2 2 3 3 8 11 105 1 108
E11.5Hi 14 4 2 1 0 13 6 55 1 77
E12.5Hi 2 1 9 1 0 4 0 12 3 14
E10.5Lo 6 3 3 1 1 2 3 12 3 12
E11.5Lo 5 7 2 1 1 8 4 30 1 24
E12.5Lo 14 1 0 1 0 5 10 81 0 87
NonDE 3 2 0 1 0 2 0 6 0 13
trajectory shifts concordant with changes in cellular abundance
H
Number of genes with concordant shifts in expression and cellular abundance
WT Expression Trajectory
E10.5Hi 302
E11.5Hi 131
E12.5Hi 115
E13.5Hi 150
E10.5Lo 69
E11.5Lo 35
E12.5Lo 89
E13.5Lo 110
EarlyHi 102
LateHi 157
E10.5Hi & E13.5Hi 65
Down-Up-Down 55
Up-Down-Up 59
NonDE 221
Low confidence 59
I
Total number of MUT trajectory shifts that correspond with shifts in cellular composition
27.2% concordant (1,719 genes)
72.8% Disconcordant (4,608 genes)
Concordant
Disconcordant

**Figure S28. Dissecting the relationship between gene expression trajectories and shifts in cellular abundance in WT and mutant forelimbs, related to Figure 6.** (A) Workflow for assessing whether temporal gene expression trajectories reflect changes in cellular composition. For each gene, the proportion of cells expressing the gene ($\geq$ 1 count) at each timepoint was calculated and normalized to the total number of cells captured at that timepoint. The temporal proportions were then analyzed using WATER to define cellular abundance trajectories. (B) Temporal kinetics of cellular composition shifts in WT and mutant limbs. WATER was applied to gene-specific cellular proportions across time, revealing six temporal abundance patterns in WT and mutant datasets. The number of genes per genotype and trajectory is indicated to the left. (C) Example of a gene (*Acan*) whose expression trajectory and cellular abundance are concordant. Left, bulk expression trajectory for WT (solid line, circles) and mutant (dashed line, triangles) with biological replicates overlaid. Right, fraction of *Acan*-expressing cells in WT (solid line) and mutant (dashed line). (D-E) Example of a gene (*Msx1*) previously identified as shifting from WT Late high to mutant E12.5 High. Despite this shift in bulk expression trajectory, the temporal pattern of cellular abundance is largely conserved between WT and mutant (D), although the overall proportion of *Msx1*-expressing cells is elevated in the mutant. This suggests that the observed shift in *Msx1* expression cannot be attributed to changes in cell abundance alone. Consistently, median *Msx1* expression per cell is higher in the mutant (E), supporting a per-cell increase in RNA abundance rather than a shift in cell composition as the primary driver. (F) Analytical framework to test the hypothesis that a subset of expression shifts between WT and mutant correspond to shifts in cellular composition. (G) Intersection of mutant expression trajectories and mutant cellular abundance trajectories for genes that shift from WT E10.5 high to alternative trajectories in the mutant. Genes whose shifted expression trajectories are concordant with corresponding changes in cellular abundance are highlighted with red boxes. (H) Number of genes with expression shifts that are concordant with cellular abundance trajectories, subset by WT trajectory. (I) Percentage of total mutant trajectory shifts that correspond to shifts in cellular composition. Approximately 27.2% of mutant trajectory shifts are concordant with changes in cellular abundance, while 72.8% cannot be explained by cellular composition alone. See also Data S17.

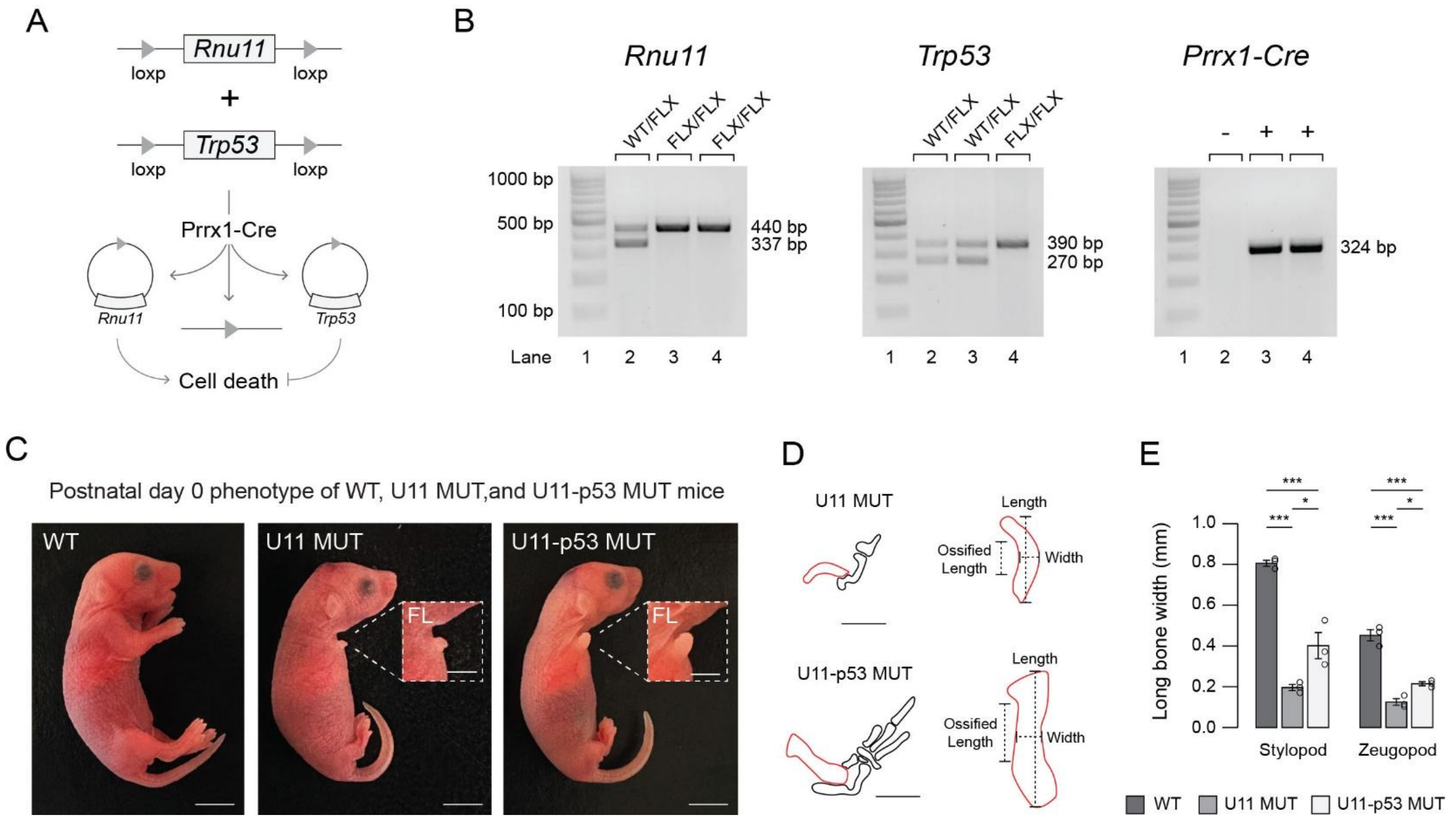


**Figure S29. Phenotypic analysis of U11-p53 mutant limbs, related to Figure 7.** (A) Schematic of the genetic strategy used to conditionally delete Rnu11 and Trp53 from the developing limb using *Prrx1*-Cre. (B) PCR-based genotyping of *Rnu11*, *Trp53*, and *Prrx1*-Cre alleles. Band sizes corresponding to wildtype and floxed alleles are indicated. (C) Representative images of postnatal day 0 (P0) WT, U11 mutant, and U11-p53 mutant mice. Insets highlight forelimbs (FL), illustrating partial restoration of limb size in U11-p53 mutant relative to U11 mutant. (D) Schematic representation of measurement strategy for long bone length, width, and ossified length in U11 mutant and U11-p53 mutant forelimbs. (E) Quantification of stylopod and zeugopod long bone width across genotypes. Each point represents a biological replicate, and bars indicate mean + SEM. Statistical significance was determined by a one-way ANOVA followed by post-hoc Tukey's test. * $p < 0.05$, *** $p < 0.001$.

**Supplementary Tables**

**Table S1. Summary statistics from limb surface area quantifications, related to Figure 1.**

| **Comparison** | | | **Adj. p-value** | **Significance** |
|---|---|---|---|---|
| E10.5 WT | vs. | E11.5 WT | 1.00E-07 | *** |
| E10.5 WT | vs. | E12.5 WT | 0.00E+00 | *** |
| E10.5 WT | vs. | E13.5 WT | 0.00E+00 | *** |
| E11.5 WT | vs. | E12.5 WT | 1.00E-07 | *** |
| E11.5 WT | vs. | E13.5 WT | 0.00E+00 | *** |
| E12.5 WT | vs. | E13.5 WT | 2.55E-05 | ** |
| E10.5 MUT | vs. | E11.5 MUT | 3.27E-05 | *** |
| E10.5 MUT | vs. | E12.5 MUT | 4.00E-07 | *** |
| E10.5 MUT | vs. | E13.5 MUT | 1.20E-06 | *** |
| E11.5 MUT | vs. | E12.5 MUT | 2.29E-01 | n.s. |
| E11.5 MUT | vs. | E13.5 MUT | 7.76E-01 | n.s. |
| E12.5 MUT | vs. | E13.5 MUT | 9.47E-01 | n.s. |
| Significance codes: n.s. >0.05, * <0.05, ** <0.01, *** <0.001 | | | | |

**Table S2. Comparison of WATER with other analysis methods for temporal RNA-seq data, related to Figure S10.***

| **Capability** | **WATER (this study)** | **TC-Seq (Wu & Simko, Bioconductor)**[3] | **Mfuzz (Kumar et al. 2007)**[18] | **ImpulseDE2 (Fischer et al. 2018)**[19] | **maSigPro (Nueda et al. 2014)**[20] |
|---|---|---|---|---|---|
| Uses full time-vector clustering | No | Yes | Yes | No | No |
| Overlapping window comparisons | Yes | Optional support | No | No | No |
| Detection of transient/biphasic expression patterns | Yes (via window comparisons and iterative refinement) | potential detection via soft clustering | potential detection via soft clustering | Yes (via impulse model fitting) | Yes (via polynomial regression) |
| Cross-condition trajectory mapping | Yes | No | No | Optional condition comparison without trajectory mapping | Optional multi-condition regression modeling |
| Explicit modeling of gene expression as continuous functions | No | No | No | Yes (via impulse modeling) | Yes (via polynomial regression) |
| Statistical testing for differential expression across time | No (performed upstream) | Limited support | No | Yes | Yes |
| Sensitivity to number of timepoints | Low-Moderate | Moderate | Moderate | High | Moderate-High |

* Methods differ substantially in their intended analytical goals. This table highlights methodological distinctions rather than performance comparisons.

**Table S3. Redistribution of functional modules across temporal trajectories following minor spliceosome inhibition, related to Figure 2. See also Data S7.**

| GO Term | WT Enriched Trajectory | # WT Genes | Mut Enriched Trajectories | # Genes per Mut Trajectory | Interpretation |
|---|---|---|---|---|---|
| DNA replication | E10.5$^{Hi}$ | 43 | Early$^{Hi}$ / E11.5$^{Hi}$ / NonDE | 18 / 15 / 10 | Redistribution of early proliferative programs into sustained or unresolved states Persistence of cell cycle programs |
| Mitotic cell cycle phase transition | E10.5$^{Hi}$ | 36 | Early$^{Hi}$ / E11.5$^{Hi}$ / NonDE | 14 / 11 / 11 | |
| Nuclear Division | E10.5$^{Hi}$ | 29 | Early$^{Hi}$ / NonDE | 17 / 12 | Prolonged usage of early proliferative modules |
| Chromosome Organization | E10.5$^{Hi}$ | 24 | Early$^{Hi}$ / E11.5$^{Hi}$ | 13 / 11 | Disrupted early chromatin organization |
| RNA processing | E11.5$^{Hi}$ | 38 | Early$^{Hi}$ / Late$^{Hi}$ / NonDE | 16 / 12 / 10 | Loss or flattening of transitional regulatory phase |
| RNA splicing | E11.5$^{Hi}$ | 34 | Early$^{Hi}$ / Late$^{Hi}$ | 18 / 16 | |
| Ribonucleoprotein complex biogenesis | E11.5$^{Hi}$ | 27 | Early$^{Hi}$ / NonDE | 14 / 13 | |
| Histone modifying activity | E12.5$^{Hi}$ | 31 | Late$^{Hi}$ / E13.5$^{Hi}$ | 17 / 14 | Delayed or prolonged epigenetic gating |
| Polycomb group (PcG) protein complex | E12.5$^{Hi}$ | 14 | Late$^{Hi}$ / NonDE | 9 / 5 | Altered temporal positioning of repressive chromatin complexes |
| SWI/SNF superfamily-type complex | E12.5$^{Hi}$ | 19 | Late$^{Hi}$ / E13.5$^{Hi}$ | 11 / 8 | Redistribution of chromatin remodeling modules to later states |
| Regulation of cellular response to growth factor stimulus | E12.5$^{Hi}$ | 22 | Late$^{Hi}$ | 22 | Preserved signaling with shifted temporal deployment |
| Skeletal system morphogenesis | E13.5$^{Hi}$ | 41 | Late$^{Hi}$ / E13.5$^{Hi}$ | 21 / 20 | Delayed or redistributed differentiation-associated programs |
| Osteoblast differentiation | E13.5$^{Hi}$ | 26 | Late$^{Hi}$ | 26 | Differentiation programs retained but temporally shifted |
| Cartilage development | E13.5$^{Hi}$ | 33 | Late$^{Hi}$ / NonDE | 19 / 14 | Partial redistribution of lineage-specific modules |
| Muscle cell differentiation | E13.5$^{Hi}$ | 29 | Late$^{Hi}$ | 29 | Temporal compression or delayed differentiation |
| Extracellular matrix organization | E13.5$^{Hi}$ | 35 | Late$^{Hi}$ / E13.5$^{Hi}$ | 17 / 18 | Redistribution of structural maturation programs |

**Table S4. Summary statistics from quantification of distal ectoderm-Sox9 gap distance and Sox9-positive area quantifications, related to Figure S27.**

| | **Sox9-positive area** | | **Distal ectoderm-Sox9 gap distance (normalized to total Sox9 length)** | | **Distal ectoderm-Sox9 gap distance (normalized to total limb length)** | |
|---|---|---|---|---|---|---|
| | **p-value** | **Significance** | **p-value** | **Significance** | **p-value** | **Significance** |
| **E10.5** | 0.009 | ** | 0.047 | * | 0.030 | * |
| **E11.5** | 0.040 | * | 0.379 | n.s. | 0.041 | * |
| **E12.5** | 0.068 | n.s. | 0.048 | * | 0.092 | n.s. |
| Significance codes: n.s. >0.05, * <0.05, ** <0.01, *** <0.001 | | | | | | |

**Table S5. RT-PCR primer sequences, related to Figure 2, 3, 6, S17, S20, S27 and S29.**

| Gene | Strand | Sequence | Usage | Fig. |
|---|---|---|---|---|
| Fgf8 | Fwd | CAGGTCCTGGCCAACAAG | In situ probe preparation | 2G |
| | Rvs | AGCTCCCGCTGGATTCCT | | |
| Eed | Fwd | GGAGACCCTCTGGTGTTTGCAACT | AS validation | 3I |
| | Rvs | GAGAAGGTTTGGGTCTCGTGGG | | |
| Chd4 | Fwd | GAGGAGGAGAAAAAGGACGTGATGCTT | AS validation | 3I |
| | Rvs | ACATCAGCTTTCATGTCACTCAACAGTTC | | |
| Actl6a | Fwd | CAGAGTTGATGTTTGAGCACTACAGCAT | IR validation | 3I |
| | Rvs | CACTGGCACATGTGCGCTGC | | |
| Ints4 | Fwd | CTTATGCTGCACTAATGTCTCAACC | IR validation | 3I |
| | Rvs | CGCCCATCTGAACTAAGCATAGAG | | |
| Ints7 | Fwd | TTCAGAGGTGCGGCCGGATCT | IR validation | 3I |
| | Rvs | CTAGGGGCTGTGGCAGTTCTTCT | | |
| Lsm5 | Fwd | ACCACGAACCCGTCCCAACTC | AS validation | 3I |
| | Rvs | TACTTCAGGCCCTTCTCCTCCG | | |
| Med23 | Fwd | AGGTTTTTAGAGTTGCTTCCAGTGTCCAA | IR validation | 3I |
| | Rvs | GTTGGCCTAAGCAACTCAAAATTAAAAATGA | | |
| Kansl2 (IR) | Fwd | GTCTCCTGACTGGCCCTGAG | IR validation | 3I |
| | Rvs | AGTAACCAGGATCTTGAGTGAAACTGAC | | |
| Kansl2 (AS) | Fwd | GTCTCCTGACTGGCCCTGAG | AS validation | S20C’ |
| | Rvs | TCTCCGGCACTTTCTGTGACTGAAT | | |
| Exo1 | Fwd | GCATTCTGTCCGGCTGTGACTA | AS validation | S20B’ |
| | Rvs | ACACTGTTCACCTCAAGTCTAGGAC | | |
| Xrcc5 | Fwd | AGTGATGACTATGTTTGTCCAACGACAG | AS validation | S20A’ |
| | Rvs | CAGAGAGATGCCAGACTTCTTCAAG | | |
| Msx1 | Fwd | ACTAGATCGGACCCCGTGGATG | Gene expression validation and in situ probe preparation | S17C & 6K |
| | Rvs | GTGGTACATGCTGTAGCCTACATG | | |
| Msx2 | Fwd | GTGGATACAGGAGCCCGGCA | Gene expression validation | S17F |
| | Rvs | AGTTGATAGGGAAGGGCAGACTG | | |
| Col2a1 | Fwd | ACGTGGAGGTGGACGCTACACTCA | Gene expression validation and in situ probe preparation | S17I & 6F |
| | Rvs | AGCCATCCTTCAGGGCAGTGTATG | | |
| Col1a1 | Fwd | ATGACTTCAGCTTCCTGCCTCAGC | In situ probe preparation | S27E |
| | Rvs | GGTGCTGTAGGTGAAGCGACTGT | | |
| Col10a1 | Fwd | TTCCTGGAGAGAAGGGTGCACAAG | In situ probe preparation | S27F |
| | Rvs | ACCCGGTTCTCCAGGATGACCT | | |
| Rnu11 (genotyping) | Fwd | ATAGTGCATGAATAGAGCACCCTCCC | Genotyping | S29B |
| | Rvs | AGGCTGCTACAGGATGACTCTGTCTTCT | | |
| Prrx1-Cre genotyping | Fwd | TATCCAGCAACATTTGGGCCAGCT | Genotyping | S29B |
| | Rvs | AAC ATT CTC CCA CCG TCA GTA CGT GA | | |
| Trp53 (genotyping) | Fwd | GGTTAAACCCAGCTTGACCA | Genotyping | S29B |
| | Rvs | GGAGGCAGAGACAGTTGGAG | | |
| Gapdh | Fwd | GACAACTTTGGCATTGTGGAAGGG | Gene expression validation | S17C, F, I |
| | Rvs | TGGATGCAGGGATGATGTTCTGG | | |
| AS; Alternative Splicing, IR; Intron Retention | | | | |
